# *Fusobacterium nucleatum* produces previously unappreciated AHR-activating metabolites and promotes CRC cell proliferation via the AHR-TERT axis

**DOI:** 10.1101/2025.09.25.678601

**Authors:** Kyoung-Jae Won, Pei-Ru Jin, Lydia Davis, Jimmy Orjala, Abigail T. Armstrong, Wen- Hung Wang, Harish Kothandaraman, Sagar M. Utturkar, Dawon Bae, Byeong-Seon Jeong, Chenggang Wu, Iain A. Murray, Gary H. Perdew, Chijian Xiang, Jianing Li, Hyunwoo Lee, Hyunyoung Jeong

## Abstract

*Fusobacterium nucleatum* (*Fn*) is prevalently enriched in colorectal cancer (CRC), promoting CRC progression. However, *Fn*-derived small molecules and their host target pathways in CRC remain largely underexplored. Here, we identify that *Fn* produces previously unrecognized three indole-containing metabolites, fusotrisindoline (FTIN), streptindole (STIN), and trisindoline (TIN). Using AHR reporter, *CYP1A1* mRNA induction, CYP1A enzyme assay, photoaffinity AHR-ligand competition, and *in silico* docking, we establish that these metabolites are bona fide agonists of the aryl hydrocarbon receptor (AHR), with FTIN being the most potent AHR ligand (EC_50_ ∼41 nM in HepG2 cells). Genetic and pharmacological perturbations demonstrate that FTIN-activated AHR promotes proliferation, migration, and invasion in EGFR blockade-responsive CRC cell lines, such as SNU-C4, but not in EGFR blockade-resistant cell lines, such as HCT116. We show that FTIN-activated AHR signaling is critical for *Fn*-mediated promotion of CRC cell growth. A Δ*tnaA* mutant defective in indole production lacked FTIN/STIN/TIN production and was unable to activate AHR or enhance CRC cell growth *in vitro* and SNU-C4 xenograft growth *in vivo*. RNA-seq and follow-up functional validation identified TERT as a key downstream effector of FTIN-activated AHR signaling. FTIN upregulated TERT transcription and telomerase activity, and TERT knockdown abrogated FTIN-promoted CRC cell proliferation. FTIN production was conserved across *Fn* subspecies, as well as other *Fusobacterium* species, and detected in additional CRC-associated genera. In 30 patient pairs of CRC-matched normal colon tissues, FTIN was quantifiable in most CRC tissues and significantly enriched relative to matched normal colon tissues, whereas STIN/TIN were undetectable. Our findings reveal a metabolite-centered FTIN-AHR-TERT axis by which intratumoral bacteria, such as *Fn,* accelerate CRC growth and uncover FTIN as a potential biomarker in CRC.

## INTRODUCTION

*Fusobacterium nucleatum* (*Fn*) is a Gram-negative anaerobic bacterium that is typically found in the oral cavity of humans.^1^ While *Fn* has generally been considered an oral commensal and opportunistic periodontal pathogen, its detection in extra-oral sites such as the intestine and uterus has been associated with extra-oral diseases in humans and animal models. For example, intrauterine infection of *Fn* is implicated to be a cause of term stillbirth^2^ and ovarian endometriosis,^3^ and the gut colonization of *Fn* is shown to worsen ulcerative colitis^4^ or rheumatoid arthritis^5^ by promoting gut microbiota dysbiosis and triggering inflammatory responses. Moreover, many studies have now established *Fn* as an intratumoral bacterium that promotes cancer development and progression in various cancer types,^6–10^ including esophageal squamous cell carcinoma (ESCC)^11,12^ and colorectal cancer (CRC).^13,14^ These broad spectra of *Fn* infection may suggest versatile pathogenic mechanisms of *Fn*, deploying host cell type-specific, as well as universal, virulence effectors that distinctly affect host cell physiology.

*Fn* is one of the most extensively studied intratumoral bacteria in CRC. Comparative analysis of microbiota from CRC tissues versus adjacent normal mucosa^15–27^ and from CRC patient versus healthy stool samples^28–34^ consistently shows *Fn* enrichment in CRC. Earlier mechanistic studies have identified that *Fn* employs multiple macromolecular adhesins (CbpF,^35,36^ FadA,^37^ Fap2,^38–41^ and RadD^41,42^) for CRC colonization and manipulation of the pathophysiology of CRC and immune cells. On the other hand, more recent studies have increasingly uncovered *Fn*-produced small-molecule metabolites that are potentially involved in CRC carcinogenesis, activate oncogenic signaling for CRC proliferation and metastasis, and promote CRC chemoresistance. Two *Fn* metabolites, DL-homocystine and allantoic acid, are shown to cause DNA damage in the SW480 human CRC cell line and in a mouse model.^43^ Adenosine-diphosphate-D-glycero-β-D-manno-heptose (ADP-heptose) and butyrate synergistically activate nuclear factor kappa-light-chain-enhancer of activated B cells (NF-κB) signaling, promoting HT-29 human CRC cell proliferation and chemoresistance to 5- fluorouracil.^44^ *Fn*-produced succinic acid induces resistance of MC38 and CT26 mouse CRC cell lines to immune checkpoint blockade.^45^ Hydrogen sulfide, a product of *Fn* metabolism of L-cysteine, enhances the migration of HCT116 and DLD-1 but not RKO cell lines.^46^ Finally, two *Fn* metabolites impact CRC cell physiology via aryl hydrocarbon receptor (AHR) signaling: Formate-mediated enhancement of AHR signaling increases cancer stemness in a mouse xenograft model using HT-29 cells.^29^ Indolepropionic acid (IPA), proposed to be an *Fn* metabolite, activates the AHR signaling in macrophages, which subsequently stimulates HCT116 cell proliferation.^47^ These findings indicate that *Fn* shapes CRC progression through a repertoire of protein adhesins and bioactive metabolites.

AHR is a ligand-activated transcription factor,^48^ and it senses structurally diverse (endogenous and exogenous) compounds as AHR agonists.^49^ While the AHR primarily regulates the expression of genes involved in xenobiotic metabolism (by inducing CYP1A1 expression) and immune and epithelial cell differentiation during homeostasis,^50–52^ *Fn*- mediated AHR activation plays a critical role in cancer progression. Besides the CRC mentioned above, *Fn* also promotes ESCC proliferation via AHR signaling,^53^ suggesting that AHR-activating *Fn* metabolites (i.e., formate and IPA) found to affect CRC physiology may be involved in the interaction of *Fn* with ESCC. On the other hand, how formate enhances the AHR signaling is unknown^29^; and although IPA is proposed to be an *Fn* metabolite,^47^ its biosynthetic pathway does not appear to exist in the genome of *Fn*.^54^ Moreover, dysregulated AHR activation in CRC cells manifests context-dependent, often opposite, phenotypes. For example, the well-known AHR agonist TCDD (2,3,7,8-tetrachlorodibenzo-*p*-dioxin) promotes the proliferation of both H508 and SNU-C4 human CRC cell lines^55^; however, it suppresses RKO cell proliferation.^56^ Therefore, whether and how *Fn* activates the AHR signaling and the role of *Fn*-mediated AHR activation in CRC remain to be defined.

In this study, we identify three indole-containing metabolites previously unknown to be produced by *Fn*, and show that they are bona fide AHR agonists. We demonstrate that *Fn*- mediated AHR activation differentially regulates *TERT* expression across CRC cell lines and promotes CRC cell proliferation via an AHR-TERT axis in a cell line-dependent manner. Moreover, we show that the most potent AHR-activating *Fn* metabolite is significantly more abundant in human CRC tissues relative to adjacent normal colon tissues and is also produced by other CRC-associated bacteria. Our findings suggest that intratumoral bacteria-mediated AHR activation likely has a broader impact on CRC progression.

## RESULTS

### *Fusobacterium nucleatum* (*Fn*) produces AHR-activating metabolites

To gain insights into the potential of the gut microbiota to activate the AHR, we selected 10 gut bacteria that represent six major phyla in the human gut microbiota. Respective cultures of these bacteria grown in brain-heart infusion containing 0.3% L-cysteine (BHIc) for 2-3 days were extracted with ethyl acetate and tested for AHR activation using a luciferase-based AHR reporter assay in HepG2 cells, where the promoter of a representative AHR target gene *Cyp1a1* drives luciferase expression. A known AHR ligand, 3-methylcholanthrene (3-MC), served as a positive control. None of the bacterial extracts used in the AHR reporter assay was cytotoxic against HepG2 cells under our tested conditions (**Figure S1A**). Interestingly, all 10 bacteria induced *Cyp1a1* promoter activities to some extent, while *Fn* exhibited the most potent activation under our culture and test conditions (**Figure 1A**). We focused on characterizing *Fn* as an AHR-activating bacterium.

**Figure 1.**
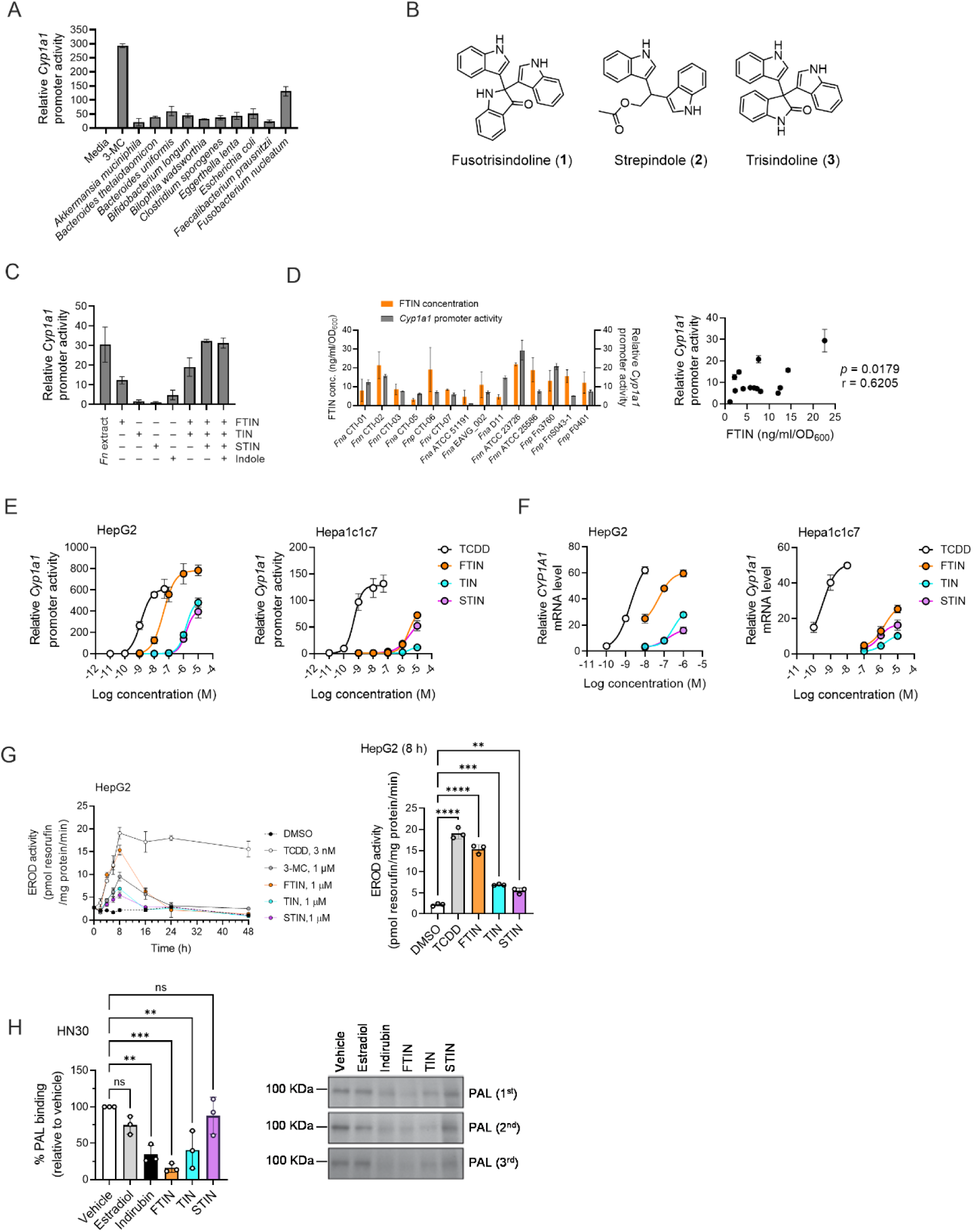
Identification of *Fusobacterium nucleatum* metabolites as AHR agonists (A) Induction of *Cyp1a1* promoter activity by gut bacterial culture extracts (*n* = 3). HepG2 cells transfected with a luciferase-based AHR-reporter system were treated with organic extracts of respective culture supernatants and 3MC (a known AHR agonist as a positive control, 3 μM) for 24 h. (B) Chemical structures of three *Fn* metabolites identified. (C) Induction of *Cyp1a1* promoter activity by *Fn* culture extracts and respective *Fn* metabolites alone or in combination (*n* = 3). FTIN (7.8 nM), STIN (1.2 nM), TIN (2.7 nM), and indole (25 µM). The *Fn* culture extract and the *Fn* metabolite concentrations tested were equivalent to 10% of the *Fn* culture. (D) Correlation between FTIN levels in the culture of *Fn* strains and *Cyp1a1* promoter activity induced by the culture extracts of *Fn* strains. Overnight cultures of respective strains grown in BHIc medium were used for FTIN measurement and the AHR reporter assays. FTIN measurement, with a limit of quantification (LOQ) of 0.2 ng/ml, was from two independent experiments (*n* = 2), and the AHR reporter assay was performed in triplicate with extract samples from a single experiment (Figure 1D, left). The correlation coefficient and associated *p*-value were obtained by Pearson correlation analysis (Figure 1D, right). (E) Induction of *Cyp1a1* promoter activity in human hepatoma HepG2 cells or murine hepatoma Hepa1c1c7 cells by synthesized *Fn* metabolites (*n* = 3). Respective cell lines transfected with the AHR reporter system were treated with varying concentrations of synthetic FTIN, TIN, STIN, or the positive control TCDD (a known AHR agonist) for 4 h. (F) Induction of *CYP1A1* mRNA expression in HepG2 cells and *Cyp1a1* mRNA expression in Hepa1c1c7 cells treated with synthetic FTIN, TIN, STIN, or TCDD (positive control) for 4 h (*n* = 3). (G) Time-course measurement of CYP1A enzymes’ ethoxyresorufin-*O*-deethylase (EROD) activity in HepG2 cells treated with FTIN (1 µM), TIN (1 µM), STIN (1 µM), 3-MC (a transient AHR agonist, 1 µM), or TCDD (a sustained AHR agonist, 3 nM). Shown is a representative of two independent experiments (*n* = 2) performed in triplicate with similar results. (H) The binding of *Fn* metabolites to the AHR was determined using an AHR photoaffinity ligand (PAL) competition assay in human quamous cell carcinoma HN30 (*n* = 3). HN30 cells were treated with respective *Fn* metabolites (FTIN, STIN, and TIN at 10 µM). Indirubin (a known AHR ligand, 100 nM) and estradiol (10 µM) were used as positive and negative controls, respectively. Data (right) represent mean percent PAL binding ± SD (normalized to AHR) relative to vehicle-treated controls. Data are represented as mean ± standard deviation (SD), where appropriate. Statistical significance was assessed by ordinary one-way ANOVA with Dunnett’s multiple comparisons test (G) and with Šídák’s multiple comparisons test (H): **, *p* < 0.01; ***, *p* < 0.001; ****, *p* < 0.0001; ns, not significant.

Because BHIc is a rich medium whose complex ingredients might interfere with the isolation of putative AHR-activating metabolite(s) produced by *Fn* and to streamline the purification of the *Fn* metabolites for structural determination, we tested whether a *Fn* culture grown in a modified M9 minimal medium containing 0.3% cysteine and 1% tryptone (M9CT) also induces *Cyp1a1* promoter activity using the AHR reporter assay. Similar to the *Fn* BHIc culture extracts, the *Fn* M9CT culture extracts increased *Cyp1a1* promoter activity (**Figure S1B**), and further examination of the *Fn* M9CT culture indicated that putative AHR-activating metabolite(s) are primarily present in culture supernatant (rather than in the bacterial cell pellets), reaching highest levels at the stationary phase (**Figure S1C**). We therefore purified AHR-activating metabolite(s) from the culture supernatant of *Fn* grown in M9CT media using chromatographic methods, determined their chemical identities using high-resolution mass spectrometry (HRMS) and nuclear magnetic resonance (NMR) spectroscopic methods, and validated their structures with custom- and in-house synthesized compounds (see **Supplemental Texts** for details). As a result, we uncovered three indole-containing metabolites previously unknown to be produced by *Fn* (**Figure 1B**). While these metabolites are new to *Fn*, each of the three metabolites has previously been isolated from facultative anaerobic bacteria or chemically synthesized in earlier studies.^57–60^ While compounds **2** and **3** were previously given traditional names streptindole (STIN)^60^ and trisindoline (TIN),^59^ respectively, compound **1** was abbreviated as BII for 2,2-*b*is(3′-*i*ndolyl)*i*ndoxyl.^57,58^ We named compound **1** as fusotrisindoline (FTIN) to emphasize its primary role in AHR activation by *Fn,* as demonstrated in this study.

In typical *Fn* cultures grown overnight in M9CT media, FTIN, STIN, and TIN were detected at concentrations of ∼80 nM (∼30 ng/mL), ∼12 nM (∼4 ng/mL), and ∼27 nM (∼10 ng/mL), respectively. We examined whether combining three metabolites accounts for the full extent of AHR activation by the whole *Fn* culture using the AHR reporter assay. We also included indole, as it is known to be a weak AHR activator^61^ and produced by *Fn*.^62^ Using high-performance liquid chromatography-ultraviolet (HPLC-UV) analysis, we determined a concentration of indole to be ∼250 µM in *Fn* cultures where FTIN, STIN, and TIN concentrations were measured. The four *Fn* metabolites (indole, FTIN, STIN, and TIN) alone or in varying combinations, at one-tenth of the respective concentrations measured in the *Fn* culture supernatant, induced *Cyp1a1* promoter activity to different extents (**Figure 1C**). FTIN was the most potent inducer, followed by indole. While STIN and TIN were poor activators individually, each metabolite additively enhanced FTIN-mediated induction of the *Cyp1a1* promoter. Interestingly, the combination of three metabolites (FTIN/STIN/TIN) with or without indole induced a similar magnitude of *Cyp1a1* promoter activity by the whole *Fn* culture extracts, suggesting a minimal role of indole in *Fn*-mediated AHR activation.

Given the finding that FTIN is produced at the highest amounts in our model strain *F. nucleatum* subspecies *nucleatum* (*Fnn*) ATCC 23726, and it is the most potent inducer of *Cyp1a1* promoter activity, we wondered whether other subspecies and strains of *Fn* also produce FTIN. A total of 14 *Fn* strains (*Fnn* ATCC 23726 as a control), including other subspecies (*animalis*, *Fna*; *polymorphum*, *Fnp*; and *vincentii*, *Fnv*),^63,64^ were grown overnight in BHIc medium, and we measured FTIN levels in their culture supernatants using LC-MS/MS and determined the induction of *Cyp1a1* promoter activity by the culture extracts using the AHR reporter assay. We found that all tested *Fn* strains produced FTIN, albeit at different levels under our culture conditions, and induced *Cyp1a1* promoter activities (**Figure 1D**), and observed a strong correlation between FTIN levels produced and *Cyp1a1* promoter-inducing activity (**Figure 1D**). Together, these results uncovered previously undescribed AHR-activating metabolites (FTIN, STIN, and TIN) produced by *Fn* and suggested that FTIN production is conserved in *Fusobacterium nucleatum*.

### *Fn*-produced indole-containing metabolites are AHR agonists

To fully determine the potency of FTIN, TIN, and STIN as AHR activators, we performed the AHR reporter assays in mouse and human cell lines, using 2,3,7,8-tetrachlorodibenzo-*p*-dioxin (TCDD; a well-known AHR agonist) as a positive control. All three *Fn* metabolites at the tested concentrations exhibited no to minimal cytotoxicity against HepG2 (human hepatoma cells) and Hepa1c1c7 (mouse hepatoma cells), respectively, under our reporter assay conditions (**Figure S1D**). While all three metabolites increased the AHR-reporter activity in both human and mouse cell lines, they exhibited ∼100-fold stronger potency in the human HepG2 cells than in the mouse Hepa1c1c7 cells (**Figure 1E**), in line with a previous report of interspecies differences between human and mouse AHRs, each binding the same AHR ligand with a different binding affinity.^65^ In HepG2 cells, FTIN was the most potent *Fn* metabolite for AHR activation, with a half-maximal effective concentration (EC_50_) of ∼41 nM. While the EC_50_ values of STIN and TIN could not be calculated due to their cytotoxicity at >10 µM, their potency appeared to be about 100-fold weaker than that of FTIN. In Hepa1c1c7 cells, FTIN and TIN induced the *Cyp1a1* promoter activity and mRNA levels, respectively, to similar extents, while STIN showed much weaker induction at concentrations up to 10 µM. We observed similar patterns of increases in the *CYP1A1* mRNA level and CYP1A1 enzyme activity in metabolite-treated HepG2 cells (**Figures 1F and 1G**), consistent with the results from the AHR reporter assays (**Figure 1E**). Different AHR agonists exhibit varying half-lives, depending on their susceptibility to host metabolism. 3-MC is an AHR ligand that is metabolized (i.e., transient) by host cells, whereas TCDD is not metabolized (i.e., sustained).^66^ To determine whether *Fn* metabolites are transient or sustained AHR ligand, we measured CYP1A enzyme activities over time in *Fn* metabolite-treated HepG2 cells. We found that CYP1A enzyme activity peaked at 8 h and returned to baseline in 24-48 h in *Fn* metabolite- treated HepG2 cells (**Figure 1G**), a pattern resembling that in 3-MC-treated HepG2 cells, suggesting a potential host metabolism of *Fn* metabolites.

Certain compounds can activate AHR indirectly without binding to the ligand-binding pocket of the AHR.^67^ We examined whether FTIN, STIN, or TIN binds to the AHR using [^125^I]- photoaffinity AHR ligand (PAL) competition assays in human HN30 cells.^68^ Indirubin (a potent AHR ligand)^69^ and estradiol (a non-AHR ligand) served as positive and negative controls, respectively. Cells were treated with respective metabolites and control compounds (at 10 μM) or vehicle, followed by the addition of [^125^I]-PAL. After 1 h, the cells were exposed to UV light to induce the covalent binding of [^125^I]-PAL to the AHR and analyzed for the signal intensity of a radioactive AHR by autoradiography. Western blot analysis demonstrated that treatment with respective metabolites and control compounds did not affect overall AHR expression in HN30 cells (**Figure S1E**). FTIN-treated cells exhibited the most significant reduction of [^125^I]-PAL-bound AHR, followed by TIN (≈ indirubin) > STIN (≈ estradiol) (**Figure 1H**). To further examine the binding of the *Fn* metabolites to the AHR, we performed *in silico* docking to the ligand-binding PER-ARNT-SIM (PAS)-B domain of the human AHR. To this end, we used the recently reported cryo-electron microscopic (cryo-EM) structure of the ternary AHR complex (Hsp90-XAP2-AHR) bound to indirubin.^70^ The docking scores of FTIN, TIN, and STIN are -12.932 kcal/mol, -12.672 kcal/mol, and -10.652 kcal/mol, respectively, indicating that FTIN has the highest binding affinity with the AHR (**Figure S1F**). In the interaction of each metabolite with the AHR protein, several amino acid residues (H291, F295, G321, Y322, F324, F351, and/or Q383) were predicted to be important for binding *Fn* metabolites. As was the case in the cryo-EM structure of ternary AHR complex bound to indirubin, which contains two indole rings, the amino acid residues H291 and F351 appeared to be critical for forming a Pi-Pi interaction with each of the three metabolites. While the essentiality of these and other residues for binding each *Fn* metabolite remained to be determined, the modeling results further support the notion that all three *Fn* metabolites bind to the AHR, and FTIN is the most potent AHR ligand. Together, these results establish three newly identified *Fn* metabolites as bona fide AHR agonists.

### FTIN promotes CRC cell proliferation via AHR activation

While *Fn* has been known to enhance CRC cell proliferation,^37^ the impact of AHR-activating *Fn* metabolites on CRC cell physiology has been unknown. To examine the effects of AHR activation by the *Fn* metabolites on CRC cells, we tested FTIN as a representative of the AHR-activating *Fn* metabolites for its impact on two distinct groups of human CRC cell lines: EGFR blockade-responsive (H508, HCT-8, LIM1215, and SNU-C4) and EGFR blockade-resistant cell lines (HCT116 and HT-29).^71,72^ The expression of the AHR protein was detected in all six CRC cell lines (**Figure S2A**). We first confirmed whether FTIN at varying concentrations (up to 1 µM) activates the AHR in these CRC cell lines using the AHR reporter assays. FTIN, as well as the control compound TCDD, increased *Cyp1a1* promoter activities in a concentration-dependent manner in respective CRC cell lines (**Figure S2B**). While precise EC_50_ values for FTIN could not be obtained due to its cytotoxicity (at >1 µM), estimated EC_50_ values of 35 - 117 nM for FTIN in six different CRC cell lines, as well as those (0.56 – 1.1 nM) for the control TCDD, were within a similar range. This result indicated that all six different CRC cell lines harbor intact AHR signaling, which FTIN can activate.

To determine the effect of FTIN on CRC cell proliferation, respective CRC cell lines were treated with FTIN (100 nM), TCDD (1 nM), or vehicle (DMSO), and their proliferation was determined using sulforhodamine B (SRB) assays. We found that FTIN, as well as TCDD, significantly stimulated the proliferation of EGFR blockade-responsive CRC cell lines (H508, HCT-8, LIM1215, and SNU-C4) (**Figure 2A**). However, neither FTIN nor TCDD promoted the proliferation of EGFR blockade-resistant cell lines (HCT116 and HT-29).

**Figure 2.**
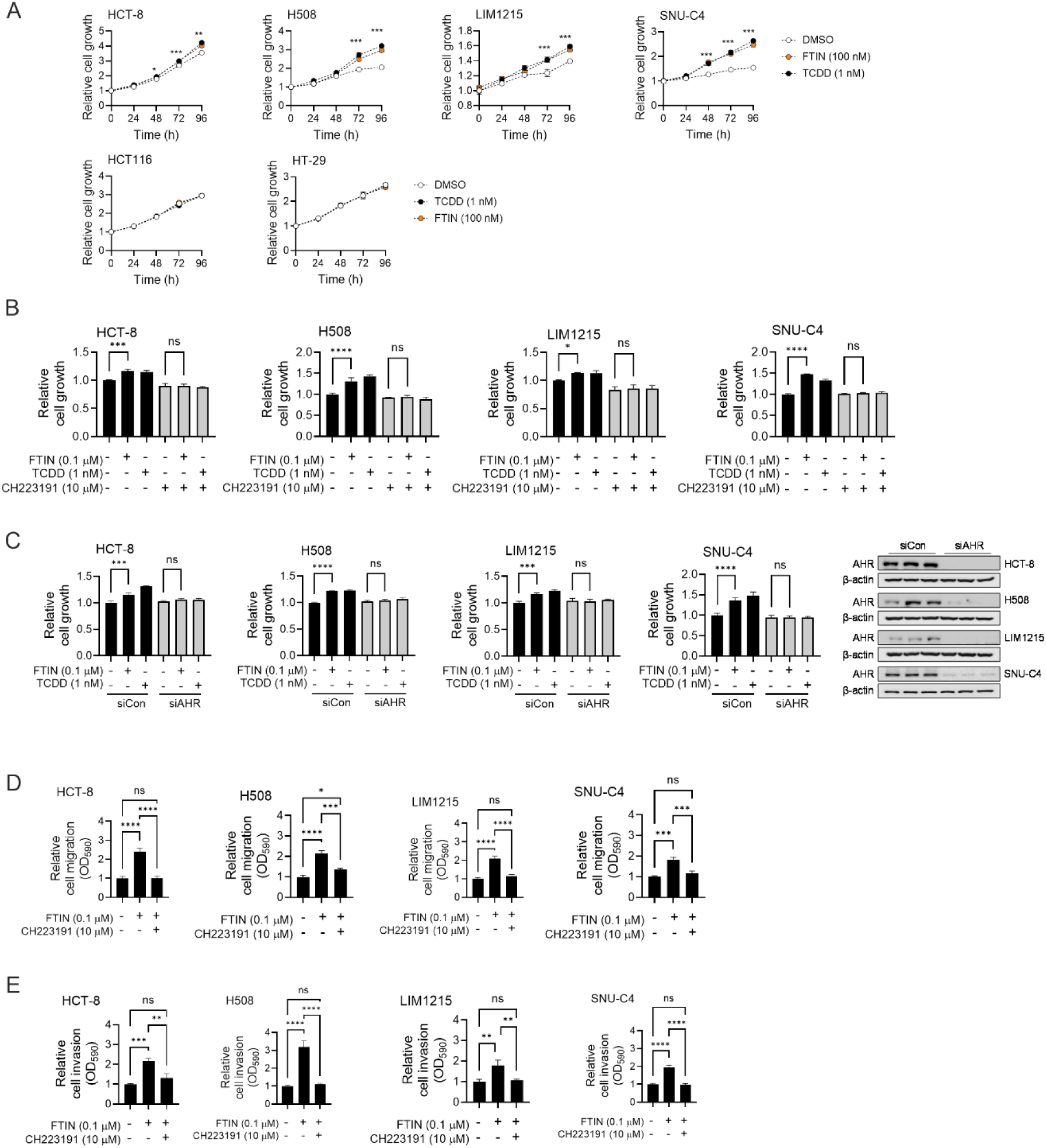
FTIN promotes CRC cell proliferation, migration, and invasion via AHR (A) FTIN promotes cell proliferation of HCT8, H508, LIM1215, and SNU-C4 CRC cell lines, but not HCT116 and HT-29 CRC cell lines (*n* = 3). Respective CRC cell lines were cultured in serum-free media containing FTIN (100 nM), TCDD (1 nM), or vehicle control (DMSO) for 96 h. The media were replaced, and cell growth was determined using the SRB assay every 24 h. (B) Treatment of the AHR antagonist CH223191 abolishes FTIN-promoted proliferation of CRC cell lines (HCT-8, H508, LIM1215, and SNU-C4) (*n* = 3). After respective CRC cell lines were pretreated with CH223191 (10 µM) for 1 h, cells were cultured in serum-free media containing FTIN (100 nM), TCDD (1 nM), or vehicle (DMSO) for 72 h. The media were replaced, and cell growth was determined using the SRB assay every 24 h. (C) AHR knockdown by siAHR abrogates FTIN-promoted proliferation of CRC cell lines (HCT-8, H508, LIM1215, and SNU-C4) (*n* = 3). Respective CRC cell lines were transfected with siAHR or siCon (control siRNA) (20 nM each) for 24 h and subsequently cultured in serum-free media containing FTIN (100 nM), TCDD (1 nM), or vehicle (DMSO) for 72 h. The media were replaced, and cell growth was determined using the SRB assay every 24 h. Shown on the right is the assessment of AHR protein expression at 72 h post-transfection of siAHR or siCon. (D, E) Cell migration (D) and invasion (E) were determined by the Transwell assay. Respective CRC cell lines were pretreated with CH223191 (10 µM) for 1 h, followed by FTIN (100 nM) or vehicle (DMSO) treatment for 24 h. Data are represented as mean ± SD, where appropriate. Statistical significance was assessed by a unpaired *t*-test (at the indicated time point in Figure 2A, DMSO *vs*. FTIN and DMSO *vs*. TCDD) and ordinary one-way ANOVA with Tukey’s multiple comparisons test (Figures 2B- 2E): *, *p* < 0.05; **, *p* < 0.01; ***, *p* < 0.001; ****, *p* < 0.0001; ns, not significant.

To determine whether FTIN-mediated promotion of CRC cell proliferation requires AHR signaling, we employed two different approaches: Chemical inhibition of the AHR signaling by the AHR antagonist CH223191 and AHR knockdown using an siRNA targeting AHR (i.e., siAHR). When each of the four CRC cell lines (H508, HCT-8, LIM1215, and SNU-C4) was co-treated with FTIN (or TCDD) and CH223191, we no longer observed FTIN- (or TCDD-) mediated promotion of CRC cell proliferation (**Figure 2B**). Consistently, we observed using AHR reporter assays that AHR activation in respective CRC cell lines (H508, HCT-8, LIM1215, and SNU-C4) by FTIN (or TCDD) was completely abolished by CH223191 treatment (**Figure S2C**). Moreover, respective CRC cell lines (H508, HCT-8, LIM1215, and SNU-C4) transfected with siAHR (which efficiently blocked the AHR expression) did not exhibit FTIN-mediated growth promotion (**Figure 2C**). We also observed that FTIN enhanced the migration and invasion of HCT-8, H508, LIM1215, and SNU-C4 CRC cell lines, respectively, which was abrogated when co-treated with CH223191 (**Figures 2D** and **2E**). Similar to the lack of FTIN effect on HCT116 cell proliferation, FTIN did not enhance the migration and invasion of HCT116 cells (**Figures S2D** and **S2E**), whereas TGF-β (used as a positive control) promoted HCT116 cell migration and invasion. Together, these results indicate that FTIN enhances the proliferation, migration, and invasion properties in EGFR blockade-responsive CRC cell lines through AHR activation.

### Production of AHR-activating metabolites requires indole as an intermediate in *Fn*

The three AHR-activating *Fn* metabolites contain two (i.e., STIN) or three (i.e., FTIN and TIN) indole moieties, suggesting indole as an intermediate in the biosynthesis of these metabolites in *Fn* (**Figure 3A**). In bacteria, indole can be produced from an indole-containing precursor (i.e., indole-3-glycerol phosphate) by tryptophan synthase during L-tryptophan biosynthesis.^73^ Alternatively, L-tryptophan is converted to indole by tryptophanase (encoded on the *tnaA* gene).^74^ The genome of a *Fn* strain ATCC 25586 has been predicted to lack the L-tryptophan biosynthetic pathway,^75^ and we confirmed the L-tryptophan auxotrophy of our model *Fn* strain ATCC 23726, as well as strain ATCC 25586 (**Figure S3A**), suggesting that the source of indole in *Fn* is L-tryptophan catabolism mediated by TnaA.

**Figure 3.**
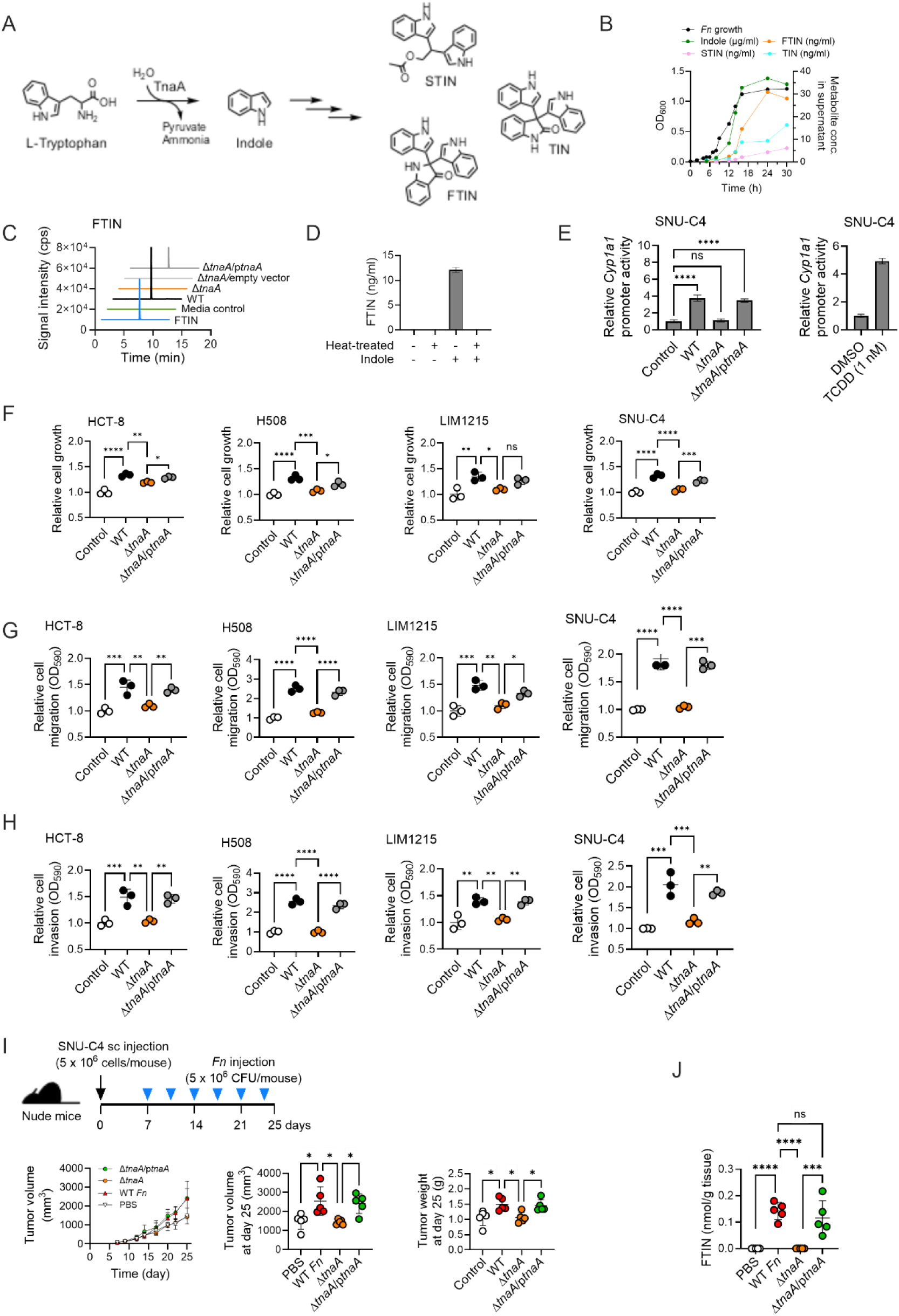
TnaA is required for FTIN production and for *Fn*-mediated promotion of CRC cell proliferation *in vitro* and tumor growth *in vivo* (A) Proposition of indole as an intermediate for the biosynthesis of FTIN, STIN, and TIN in *Fn*. The TnaA enzyme converts L-tryptophan into indole. (B) Time-course measurement of *Fn* metabolites (FTIN, STIN, and TIN) in the culture of the wild-type *Fn* ATCC 23726 strain grown in M9CT media. Shown is a representative of two independent experiments (*n* = 2) with similar results. OD_600_: optical density at 600 nm. (C) LC-MS/MS spectra of FTIN in cultures of wild-type (WT) *Fn*, Δ*tnaA*, Δ*tnaA*/empty vector, and Δ*tnaA*/p*tnaA* strains grown in M9CT media for 24 h. Shown is a representative LC-MS/MS spectrum from two independent experiments (*n* = 2) with similar results. (D) Production of FTIN in the Δ*tnaA* culture supplemented with indole (*n* = 2). Heat-treated or intact BHIc overnight culture of the Δ*tnaA* mutant was supplemented with indole (500 µM) and incubated for 4 h, and the culture supernatants were collected for FTIN measurement. (E) The Δ*tnaA* mutant is defective in inducing *Cyp1a1* promoter activity (*n* = 3). SNU-C4 CRC cells transfected with the AHR reporter system were co-incubated with the respective *Fn* strains (at MOI 300) or were treated with TCDD for 4 h. TCDD was used as a positive control for the AHR reporter assay. (F) The Δ*tnaA* mutant is impaired in promoting the proliferation of CRC cells (*n* = 3). Shown is the result at the 72 h time point (see Figure S3 for the result of the whole-time course up to 96 h). Respective CRC cell lines (HCT-8, H508, LIM1215, and SNU-C4) were co-incubated with an *Fn* strain (WT, Δ*tnaA*, or Δ*tnaA*/p*tnaA* at MOI 300) in serum-free media and cultured over time up to 96 h. Media containing an *Fn* strain were renewed, and cell growth was determined by the SRB assay every 24 h. (G and H) The Δ*tnaA* mutant is defective in promoting CRC cell migration (G) and invasion (H) (*n* = 3). The migration and invasion of respective CRC cell lines (HCT-8, H508, LIM1215, and SNU-C4) were determined by the Transwell assay. (I) Mouse xenografts with the SNU-C4 CRC cell line. Nude mice (*n* = 5 per group) were subcutaneously injected with SNU-C4 cells. After tumor formation, respective *Fn* strains (5×10^6^ colony-forming units (CFU)/injection) were intratumorally injected twice a week. Tumor volume was measured at the indicated time points, and tumor weight at the endpoint. (J) FTIN measurement in xenograft tumors (*n* = 5 per group). Two pieces of cuts from each tumor sample were processed for FTIN measurement as described in Experimental Method Details, and the average value of the two measurements was used. Data are represented as mean ± SD, where appropriate. Statistical significance was assessed by ordinary one-way ANOVA with Tukey’s multiple comparisons test (Figures 3E-3I): *, *p* < 0.05; **, *p* < 0.01; ***, *p* < 0.001; ****, *p* < 0.0001; ns, not significant.

To determine whether indole is required for the biosynthesis of FTIN, STIN, and/or TIN in *Fn*, we constructed a *tnaA* deletion mutant (Δ*tnaA*) and examined its production of FTIN, STIN, and TIN, as well as indole. In a typical wild-type *Fn* culture grown in M9CT media, indole production reached its highest level in the early stationary phase, followed by FTIN/STIN/TIN production (**Figure 3B**). The Δ*tnaA* mutant exhibited wild-type growth (**Figure S3B**); however, none of the four metabolites (indole, FTIN, STIN, and TIN) were detected in the culture of the *tnaA* mutant (**Figure 3C** and **Figure S3C**). The complementation of the Δ*tnaA* mutant with a plasmid expressing TnaA (i.e, Δ*tnaA*/p*tnaA*) restored the production of all four metabolites. Moreover, indole addition to growth media also restored FTIN production in the Δ*tnaA* mutant, but not in heat-killed bacteria (**Figure 3D**). These results indicate that indole serves as an intermediate for the biosynthesis of all three metabolites (FTIN, STIN, and TIN).

Given that we could not detect the three AHR-activating metabolites in the culture of the Δ*tnaA* mutant, we examined whether the Δ*tnaA* mutant activates AHR in CRC cells using the AHR reporter assays. We found that when co-incubated with respective CRC cell lines, the Δ*tnaA* mutant was unable to activate the AHR in SNU-C4 (**Figure 3E**) or H508 CRC cell lines (**Figure S3D**), compared with the wild-type *Fn*. A recent study reported that *Fn* produces 3- indolepropionic acid,^47^ a weak agonist of the AHR with an EC_50_ value of ∼100 µM in a luciferase-based AHR reporter assay similar to ours.^76,77^ However, we were unable to detect 3- indolepropionic acid (IPA) in the culture of the wild-type *Fn* grown in M9CT media, with a detection limit of IPA at 0.8 ng/ml (**Figure S3E**), ruling out the contribution of IPA to *Fn*- mediated AHR activation under our test conditions. Together, these results indicate that indole is an intermediate of all three AHR-activating metabolites (FTIN, STIN, and TIN) produced in *Fn* and demonstrate that TnaA, which converts L-tryptophan to indole, is required for the biosynthesis of FTIN, STIN, and TIN in *Fn*.

### The Δ*tnaA* mutant is unable to promote CRC cell proliferation *in vitro* and *in vivo*

Given that the Δ*tnaA* mutant was unable to activate the AHR, we examined the contribution of *Fn*-mediated AHR activation to the *Fn*-mediated promotion of CRC cell proliferation. Wild-type *Fn*, when co-incubated, promoted the proliferation of all four CRC cell lines (HCT-8, H508, LIM1215, and SNU-C4) (**Figure 3F** and **Figure S3F**). The Δ*tnaA* mutant, however, was unable to promote the proliferation of HCT-8, H508, LIM1215, or SNU-C4 cell lines, while the complemented Δ*tnaA*/p*tnaA* strain exhibited the wild-type promoting activity. By contrast, both the wild-type *Fn* and the Δ*tnaA* mutant promoted the proliferation of both HCT116 and HT-29 cells (**Figure S3G**). Similar to the differential effects of the Δ*tnaA* mutant on the proliferation of different CRC cell lines, we observed that the wild-type *Fn* enhanced the migration and invasion in HCT-8, H508, LIM1215, or SNU-C4 CRC cell lines, the Δ*tnaA* mutant lost such activity, and the complemented Δ*tnaA*/p*tnaA* strain restored the phenotypes (**Figures 3G** and **3H**). However, the Δ*tnaA* mutant exhibited wild-type *Fn*-mediated promotion of migration and invasion in both HCT116 and HT-29 cell lines (**Figure S3H**). These results indicate that TnaA-involved biosynthesis of AHR-activating metabolites is necessary for the *Fn*-mediated promotion of proliferation, migration, and invasion in EGFR blockade-responsive CRC cell lines, but not in EGFR blockade-resistant CRC cell lines.

We further examined the contribution of *Fn*-mediated AHR activation to the *Fn*-mediated promotion of CRC cell growth in a mouse xenograft model. Our preliminary study determined that HCT-8, H508, and LIM1215 CRC cell lines have low penetrance, being unable to develop metastasis in nude or NSG mice (data not shown), and SNU-C4 cells were known to have low metastatic potential and poor penetrance in mice.^78^ Thus, we focused on examining *Fn*- mediated promotion of SNU-C4 CRC cell growth using subcutaneous xenografts in mice. SNU-C4 cells were xenografted subcutaneously into nude mice, and the effects of intratumoral *Fn* injection on tumor growth were monitored. Consistent with the *in vitro* results (**Figure 3F**), we observed that the wild-type *Fn* significantly enhanced the tumor growth of SNU-C4 cells, the Δ*tnaA* mutant lost such enhancing activity, and the complemented Δ*tnaA*/p*tnaA* strain restored it (**Figure 3I**). The Δ*tnaA* mutant displayed wild-type growth *in vitro* (**Figure S3B**) and wild-type levels of attachment to and invasion into SNU-C4 CRC cells (**Figure S3I**), while a control Δ*fap2* mutant (which exhibits wild-type growth *in vitro*; **Figure S3B**) was defective in both CRC cell attachment and invasion as previously reported,^38^ indicating that the observed phenotype of the Δ*tnaA* mutant is neither due to its growth defect nor impaired interaction with CRC cells. Remarkably, we detected significant amounts of FTIN, but not STIN and TIN, in tumor tissues infected with either wild-type *Fn* or the complemented Δ*tnaA*/p*tnaA* strain, but not in those infected with the Δ*tnaA* mutant (**Figure 3J**). These results suggest that intratumoral *Fn* produces FTIN *in vivo* and that TnaA-involved biosynthesis of AHR-activating metabolites is essential for the *Fn*-mediated promotion of tumor growth in mice.

### *Fn*-mediated promotion of CRC cell proliferation requires AHR

To determine the requirement of the AHR in *Fn*-mediated promotion of CRC cell proliferation, we used two different approaches: the AHR antagonists (CH223191 and StemRegenin 1) and *AHR*-knockout CRC cell lines (SNU-C4^ΔAHR^ and HCT116^ΔAHR^) constructed using the CRISPR-Cas9 mutagenesis (**Figure S4A**). While the AHR antagonist CH223191 at the concentration (10 µM) used did not inhibit *Fn* growth (**Figure S4B**), its inhibition of AHR activity completely abrogated *Fn*-mediated induction of *Cyp1a1* promoter activity in SNU-C4 cells (**Figure S4C**) and partially attenuated *Fn*-mediated promotion of SNU-C4 cell proliferation (**Figure 4A**). We also obtained similar results with another AHR antagonist StemRegenin 1 (SR1) (**Figures S4D** and **S4E**). In the *AHR*-knockout cell line SNU-C4^ΔAHR^, we confirmed using the AHR reporter assays that neither FTIN, TCDD (a positive control), nor wild-type *Fn* induced *Cyp1a1* promoter activities (**Figure S4F**). When the wild-type *Fn* was co-incubated with respective *AHR*-knockout cell lines, we observed that *Fn* did not promote SNU-C4^ΔAHR^ cell proliferation (**Figure 4B**), whereas it did promote HCT116^ΔAHR^ cell proliferation (**Figure S4G**), compared with respective parent CRC cell lines. To further examine the *in vivo* relevance, parent (SNU-C4/gAHR) or *AHR*-knockout SNU-C4^ΔAHR^ cells were xenografted subcutaneously into nude mice, and the effect of intratumoral *Fn* injection on tumor growth was monitored. As expected, *Fn* enhanced the growth of parent SNU- C4/gAHR tumors compared with vehicle control, but not SNU-C4^ΔAHR^ tumors (**Figure 4C**). Similar to the xenograft experiment (wild-type *Fn vs*. Δ*tnaA*) above (**Figure 3J**), we detected significant amounts of FTIN in both *Fn*-infected SNU-C4/gAHR and SNU-C4^ΔAHR^ tumors, but not in PBS-treated SNU-C4/gAHR and SNU-C4^ΔAHR^ tumors (**Figure 4D**), indicating that the lack of *Fn*-mediated promotion of SNU-C4^ΔAHR^ tumor growth is not due to impaired FTIN production. Together, these results further confirm that *Fn*-mediated AHR activation has no impact on HCT116 cell proliferation and demonstrate that AHR is required for the *Fn*-mediated promotion of SNU-C4 cell proliferation *in vitro* and *in vivo*.

**Figure 4.**
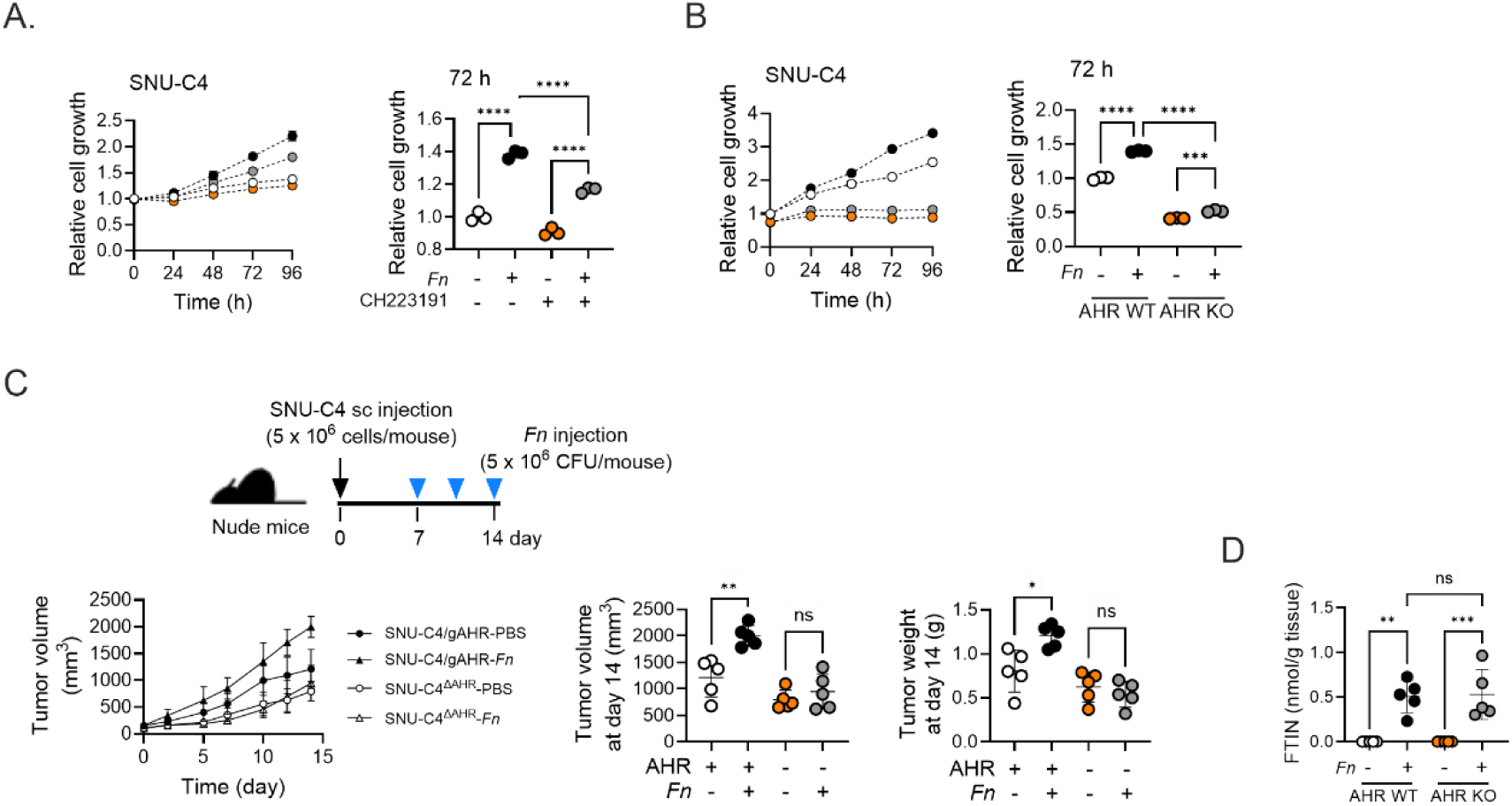
AHR is required for *Fn*-mediated promotion of CRC cell proliferation *in vitro* **and tumor growth *in vivo*** (A) *Fn*-mediated promotion of SNU-C4 CRC cell proliferation was partially abrogated by the AHR antagonist (*n* = 3). SNU-C4 cells pretreated with CH223191 (10 µM) for 1 h were co- incubated with *Fn* ATCC 23726 (at MOI 300) in serum-free media for 96 h. The media containing the *Fn* strain was replaced, and cell growth was determined using the SRB assay every 24 h. (B) *Fn* is unable to promote cell proliferation of SNU-C4^ΔAHR^ cells (*n* = 3). The parent SNU- C4/gAHR and SNU-C4^ΔAHR^ cells were co-incubated with *Fn* ATCC 23726 (at MOI 300) in serum-free media for 96 h. The media containing the *Fn* strain was replaced, and cell growth was determined using the SRB assay every 24 h. (C) Mouse xenografts with SNU-C4^ΔAHR^ CRC cell line. Nude mice (*n* = 5 per group) were subcutaneously injected with either the parent SNU-C4/gAHR or SNU-C4^ΔAHR^ cells. After tumor formation, wild-type *Fn* cells (5×10^6^ CFU/injection) were intratumorally injected twice a week. Tumor volume was measured at the indicated time points, and tumor weight at the endpoint. (D) FTIN measurement in xenograft tumors (*n* = 5 per group). Two pieces of cuts from each tumor sample were processed for FTIN measurement as described in Experimental Method Details, and the average value of the two measurements was used. Data are represented as mean ± SD, where appropriate. Statistical significance was assessed by ordinary one-way ANOVA with Tukey’s multiple comparisons test (Figures 4A-4D): *, *p* < 0.05; **, *p* < 0.01; ***, *p* < 0.001; ****, *p* < 0.0001; ns, not significant.

### FTIN promotes CRC cell proliferation via the genomic action of the AHR

The inactive AHR cytosolic complex consists of four proteins, including the Src kinase.^79^ Upon binding of an AHR agonist such as TCDD, the AHR may function as a transcription regulator in the nucleus (i.e., genomic action) and/or as a substrate-specific adaptor component of a ubiquitin ligase complex in the cytosol (i.e., non-genomic action).^80,81^ In addition, the Src kinase liberated from the AHR complex phosphorylates and activates its target proteins (i.e., an indirect effect of the ligand-dependent AHR activation).^79^ A previous study has reported that the AHR agonist TCDD stimulates the proliferation of SNU-C4 and H508 CRC cells via the Src kinase, which was proposed to phosphorylate and activate the EGFR-ERK kinase cascade for promotion of CRC cell proliferation.^55^ Given our observations that FTIN, as well as TCDD used as a control, promotes SNU-C4 and H508 cell proliferation (**Figure 2A**), we reasoned that FTIN would induce the Src-EGFR-ERK signaling pathway, accounting, at least in part, for the FTIN promotion of CRC cell proliferation. Surprisingly, however, neither TCDD nor FTIN activated the Src kinase in SNU-C4 or H508 CRC cells, whereas the positive control EGF stimulated the phosphorylation of EGFR as expected (**Figure S5A** and **Figure S5B**).^82^ These results ruled out the Src-EGFR-ERK kinase cascade being a downstream mediator of FTIN-activated AHR signaling in SNU-C4 and H508 CRC cells. Next, we examined whether FTIN-bound AHR functions as part of the ubiquitin ligase complex. A previous study has reported that the AHR agonist 3-MC induces AHR-dependent ubiquitylation and proteasomal degradation of β-catenin, thereby lowering β-catenin transcriptional activity in the DLD-1 CRC cell line,^81^ although this report could not be validated in another study.^83^ We performed a luciferase-based β-catenin reporter assay and observed that FTIN does not have any impact on β-catenin transcriptional activity, whereas the control CHIR99021 (GSK-3 inhibitor) induced β-catenin activation, ruling out the non-genomic action of FTIN-bound AHR (**Figure S5C**).

The ligand-bound AHR forms a heterodimer primarily with AHR nuclear translocator (ARNT) to become an active transcription regulator in the nucleus.^84^ To determine whether the genomic action of the AHR mediates the FTIN effect on CRC cells, we first examined the impact of a siRNA targeting ARNT (i.e., siARNT) on FTIN-activated AHR in SNU-C4 cells using the AHR reporter assays. We observed that ARNT expression was efficiently blocked by siARNT (**Figure 5A**), and ARNT knockdown abrogated FTIN induction of *Cyp1a1* promoter activity in siARNT-transfected SNU-C4 cells (**Figure 5A**). Further, FTIN treatment did not promote the proliferation of siARNT-transfected SNU-C4 cells, compared with the siControl-transfected SNU-C4 cells (**Figure 5B**). In addition to the AHR, ARNT (also known as hypoxia-inducible factor 1β; HIF-1β) forms a heterodimer with HIF-1α to become an active transcription regulator, HIF-1, in the nucleus.^85,86^ However, we determined using a luciferase-based HIF-1 reporter assay that FTIN does not induce HIF-1 transcriptional activities in SNU- C4 cells, while the positive control CoCl_2_ activated HIF-1 (**Figure S5D**), ruling out the possibility that the abrogation of the FTIN effect by ARNT knockdown is due to HIF-1 inactivation. We further validated the requirement of the genomic action of the AHR for the FTIN effect using a human AHR variant defective in DNA binding. This AHR variant (named AHR-GS) was previously characterized to be intact in ligand binding and nuclear translocation, but unable to bind to the AHR DNA-binding motif.^87^ SNU-C4^ΔAHR^ cells transfected with plasmid expressing AHR-GS or wild-type AHR (AHR-WT) expressed similar levels of respective proteins (**Figure 5C**), and the AHR reporter assays showed that FTIN induces *Cyp1a1* promoter activities in SNU-C4^ΔAHR^/AHR-WT cells, but not in SNU-C4^ΔAHR^/AHR-GS cells (**Figure 5C**). Moreover, FTIN treatment promoted proliferation of SNU-C4^ΔAHR^/AHR- WT cells but not SNU-C4^ΔAHR^/AHR-GS cells (**Figure 5D**), indicating that the DNA binding activity of the AHR is critical for the FTIN-promoted CRC cell proliferation. Together, these results suggest that FTIN-activated AHR promotes SNU-C4 cell proliferation via transcriptional regulation of AHR target gene(s).

**Figure 5.**
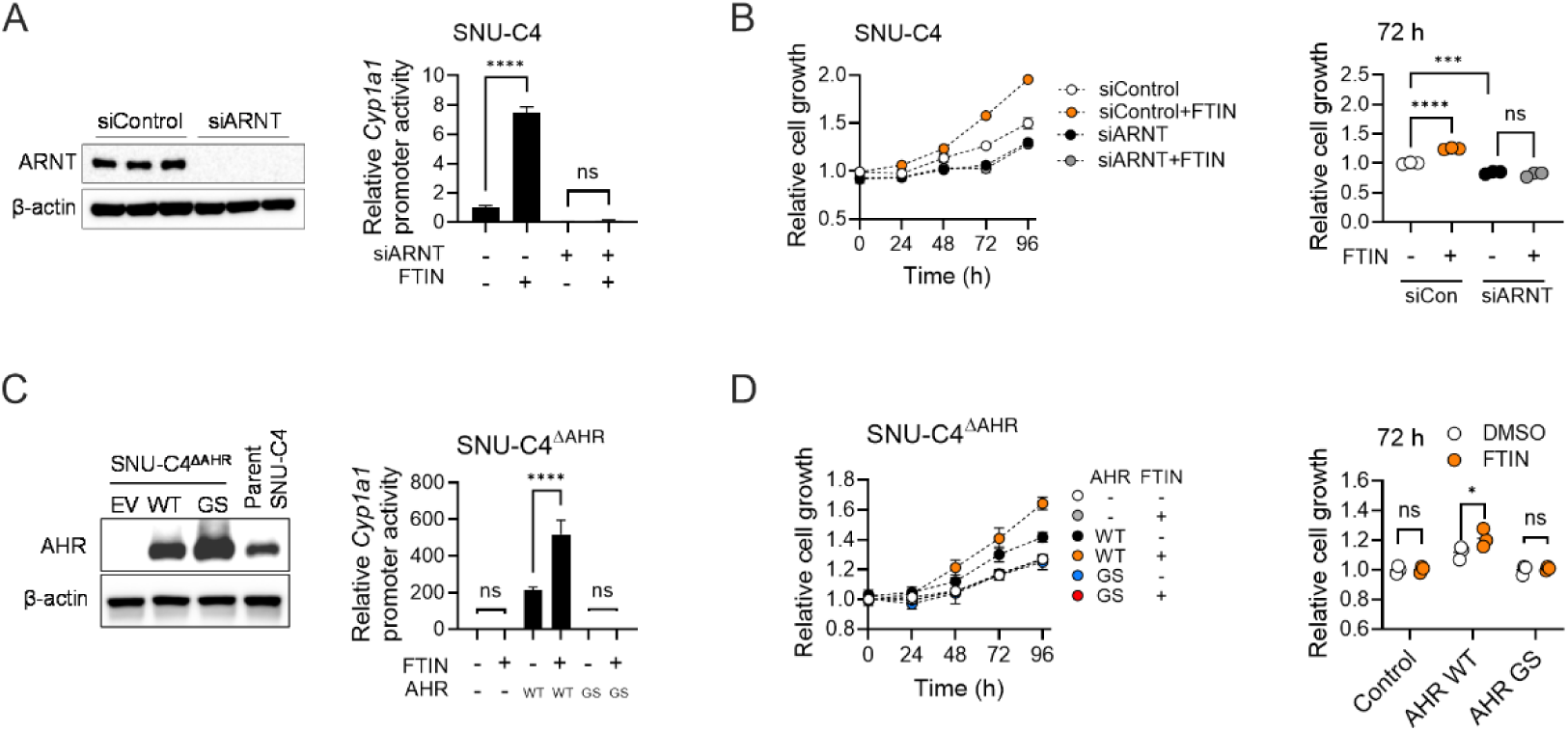
FTIN-promoted CRC cell proliferation requires the genomic action of AHR (A) *ARNT* knockdown abrogates FTIN-induction of *Cyp1a1* promoter activity (*n* = 3). SNU-C4 CRC cells transfected with siRNA (20 nM) targeting ARNT (siARNT) or control (siControl) for 24 h were transfected with the AHR reporter system for another 24 h and cultured in serum-free media containing FTIN (100 nM) or vehicle (DMSO) for 4 h. The AHR reporter assay was performed as described in Experimental Method Details, and ARNT protein levels were assessed at 72 h post-transfection of siARNT or siControl by western blot, with β-actin as a loading control (left). (B) *ARNT* knockdown abrogates FTIN-enhanced proliferation of the SNU-C4 CRC cell line (*n* = 3). SNU-C4 CRC cells transfected with siRNA (20 nM) targeting *ARNT* or control for 24 h were cultured in serum-free media containing FTIN (100 nM) or vehicle (DMSO) for 96 h. The media containing FTIN or DMSO were renewed, and cell growth was determined by the SRB assay every 24 h. (C) FTIN is unable to induce *Cyp1a1* promoter activity in the SNU-C4^ΔAHR^ cell line overexpressing DNA-binding defective AHR protein (*n* = 3). SNU-C4^ΔAHR^ cells transfected with empty vector (EV), plasmid expressing wild-type AHR (AHR-WT), or DNA-binding defective AHR (AHR-GS) were treated with FTIN (100 nM) or vehicle control (DMSO) for 4 h. The AHR reporter assay was performed as described in Experimental Method Details, and AHR-WT and AHR-GS protein levels were assessed at 28 h post-transfection of the AHR-expressing plasmid by western blot, with β-actin as a loading control (left). (D) FTIN is unable to promote the proliferation of the SNU-C4^ΔAHR^ cell line overexpressing DNA-binding defective AHR protein (*n* = 3). Respective SNU-C4^ΔAHR^ cell lines (SNU- C4^ΔAHR^/EV, SNU-C4^ΔAHR^/AHR-WT, and SNU-C4^ΔAHR^/AHR-GS) were cultured in serum- free media containing FTIN (100 nM) or vehicle control (DMSO) for 96 h. The media containing FTIN or DMSO were renewed, and cell growth was determined by the SRB assay every 24 h. Data are represented as mean ± SD, where appropriate. Statistical significance was assessed by ordinary one-way ANOVA with Tukey’s multiple comparisons test (Figures 4A-4D): *, *p* < 0.05; ***, *p* < 0.001; ****, *p* < 0.0001; ns, not significant.

### FTIN-mediated AHR activation promotes CRC cell proliferation via TERT

Given our finding that FTIN promotes proliferation of SNU-C4 and H508 CRC cell lines, but not that of HCT116 and HT29 CRC cell lines, via the genomic action of AHR, we hypothesized that AHR signaling in two different groups of CRC cells regulates the expression of distinct gene(s) that account for the phenotypic difference in response to FTIN. To identify AHR-regulated genes in SNU-C4/H508 *vs*. HCT116/HT29 CRC cell lines, we obtained gene expression profiles in FTIN-treated CRC cells using RNA-sequencing (RNA-seq). Respective CRC cell lines (SNU-C4, H508, HCT116, and HT29) were treated with FTIN (100 nM) or vehicle control for 4 h (chosen for identifying early FTIN-responsive genes) and harvested for isolation and processing of RNA for Illumina sequencing (**Table S1** for complete RNA-seq data). RNA-seq data revealed that 13 genes were differentially expressed (adjusted *p*- value<0.05 and |log_2_(fold difference)|>0.3; **Table S2**) in FTIN-treated SNU-C4 cells *vs*. vehicle-treated SNU-C4 cells (**Figure 6A**). These 13 genes included seven genes (*CYP1A1*, *CYP1B1*, *ALDH1A3*, *CEMIP*, *KSR2*, *RALGAPA2*, and *ADGRF1*) that were also upregulated in either FTIN-treated HCT116 or HT29 CRC cell lines (**Figure 6B**), implying that these seven genes are unlikely to be responsible for FTIN promotion of SNU-C4 cell proliferation. Three genes (*STMND1*, *AQP8*, and *DLEC1*) were upregulated in FTIN-treated SNU-C4 cell line but not in H508 cell line. In contrast, the remaining three genes (*TERT*, *CABLES1*, and *CAMK1D*) were upregulated in both FTIN-treated SNU-C4 and H508 cell lines, but not in FTIN-treated HCT116 and HT29 cell lines. Using qPCR, we validated the RNA-seq results of the three genes (*TERT*, *CABLES1*, and *CAMK1D*), as well as *CYP1A1* as a control, in SNU-C4 and HCT116 CRC cell lines, respectively (**Figures S6A** and **S6B**).

**Figure 6.**
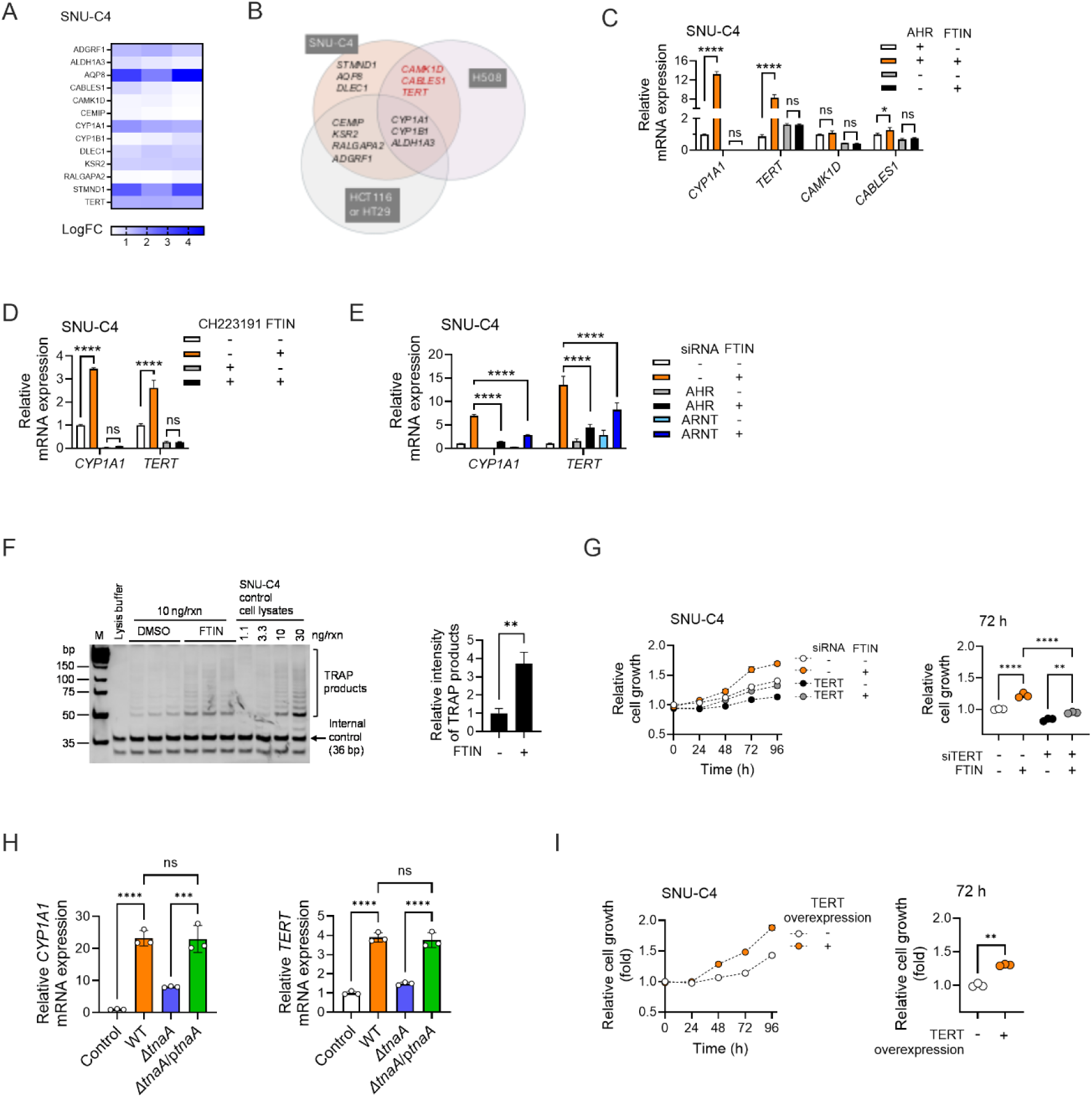
*Fn*-activated AHR promotes CRC cell proliferation by upregulating TERT expression (A) 13 FTIN-responsive genes in the SNU-C4 CRC cell line identified by RNA-seq (FTIN *vs*. DMSO, 4 h treatment) (B) Venn diagram showing FTIN-responsive genes in four different CRC cell lines (SNU-C4, H508, HCT116, and HT-29) identified by RNA-seq (FTIN *vs*. DMSO, 4 h treatment). (C) qRT-PCR validation of four FTIN-responsive genes identified by RNA-seq (*n* = 3). The parent SNU-C4/gAHR and SNU-C4^ΔAHR^ cell lines were cultured in serum-free medium containing FTIN (100 nM) for 4 h, and mRNA levels of *CYP1A1*, *CABLES1*, *CAMK1D*, and *TERT* were determined by qRT-PCR. (D) Inhibition of AHR activity by CH223191 abrogates FTIN-induction of *CYP1A1* and *TERT* mRNA expression in SNU-C4 CRC cells (*n* = 3). SNU-C4 cells were pretreated with CH223191 (10 µM) for 1 h, followed by FTIN (100 nM) treatment for 4 h. mRNA levels were determined by qRT-PCR. (E) Either *AHR* or *ARNT* knockdown partially or completely abolishes FTIN-induction of *CYP1A1* and *TERT* mRNA expression in SNU-C4 CRC cells (*n* = 3). SNU-C4 cells transfected with siRNAs (20 nM) for AHR, ARNT, or control for 72 h were treated with FTIN (100 nM) for 4 h. mRNA levels were determined by qRT-PCR. (F) TRAP assays showing FTIN-induced telomerase activity in SNU-C4 CRC cells (*n* = 3). Cells were cultured in serum-free medium containing FTIN (100 nM) for 52 h. The media were renewed every 24 h. Celly lysates prepared from SNU-C4 cells maintained in DMEM medium supplemented with 10% FBS were used as controls. (G) *TERT* knockdown abolishes FTIN-promoted proliferation of SNU-C4 CRC cells (*n* = 3). SNU-C4 CRC cells transfected with siRNAs (20 nM) for *TERT* or control for 24 h were cultured in serum-free medium containing FTIN (100 nM) or DMSO for 96 h. The media were renewed, and cell growth was determined by the SRB assay every 24 h. (H) The Δ*tnaA* mutant is defective in inducing *TERT* mRNA expression in SNU-C4 CRC cells (*n* = 3). SNU-C4 CRC cells were co-incubated with respective *Fn* strains (at MOI 300) in serum-free medium for 2 h. mRNA levels were determined by qRT-PCR. Control: media. (I) Overexpression of *TERT* is sufficient to promote the proliferation of the SNU-C4 cell line (*n* = 3). SNU-C4 CRC cells transfected with either empty vector (EV) or plasmid expressing the 3×HA-tagged TERT protein were cultured in serum-free medium for 96 h. Media were renewed, and cell growth was determined by the SRB assay every 24 h. Data are represented as mean ± SD, where appropriate. Statistical significance was assessed by ordinary one-way ANOVA with Tukey’s multiple comparisons test (Figures 6C-6E, 6G, 6H) and a unpaired *t*-test (Figures 6F and 6J): *, *p* < 0.05; **, *p* < 0.01; ***, *p* < 0.001; ****, *p* < 0.0001; ns, not significant.

Next, we determined whether the upregulation of the three FTIN-responsive genes (*TERT*, *CABLES1*, and *CAMK1D*) requires AHR by comparing their mRNA expression in the parent SNU-C4/gAHR *vs*. SNU-4^ΔAHR^ cells. Again, we observed that FTIN treatment results in upregulation of *TERT* and *CABLES1* mRNA expression in the parent SNU-C4/gAHR cells, although it did not affect *CAMK1D* expression (**Figure 6C**). In contrast, FTIN-induced *TERT* and *CABLES1* mRNA upregulation was completely abolished in SNU-C4^ΔAHR^ cells, indicating the requirement of AHR for the upregulation of these two FTIN-responsive genes. Relative to *TERT* mRNA induction by FTIN (≥ 3 fold in both RNA-seq and qPCR; **Table S1**; **Figures 6C- E** and **Figure S6A**), the FTIN induction of *CAMK1D* (1.5 fold in RNA-seq and no to 1.3 fold in qPCR) and *CABLES1* (1.3 fold in both RNA-seq and qPCR) mRNAs were small. We excluded *CAMK1D* and *CABLES1* from our follow-up investigation for an additional reason. The FTIN induction of *CAMK1D* was inconsistent in the qPCR validation of the RNA-seq result (**Figure 6C** *vs*. **Figure S6A**), and *CABLES1* has been proposed as a tumor suppressor, since the loss of *Cables1* in mice increases tumor burden in a mouse CRC model.^88^ Therefore, we focused our characterization on *TERT* and further validated the requirement of AHR for FTIN-induced upregulation of *TERT*, with *CYP1A1* as a control. In SNU-C4 cells treated with the AHR antagonist CH223191 or transfected with either siAHR or siARNT, FTIN-induced *TERT* upregulation was completely or partially abolished (**Figures 6D** and **6E**). We ruled out the possibility that FTIN-induced *TERT* upregulation is due to mRNA stabilization, as the half-life of *TERT* mRNA was similar between FTIN- and vehicle-treated SNU-C4 cells after actinomycin D was co-treated (**Figure S6C**). These results demonstrate that *TERT* expression is upregulated by FTIN-activated AHR signaling.

The human *TERT* gene encodes reverse transcriptase, a catalytic subunit that, together with another subunit, the noncoding RNA TERC, forms an active ribonucleoprotein enzyme termed telomerase that maintains telomere length in immortal (cancer) cells.^89,90^ Reactivation of otherwise silent telomerase activity is a hallmark of most, if not all, cancer types, including CRC.^91^ We sought to determine whether FTIN-induced *TERT* mRNA upregulation results in an increase in TERT protein expression using a western blot. However, three anti-TERT antibodies commercially available from different vendors were unable to detect the endogenous TERT protein in SNU-C4 cells, regardless of FTIN treatment (**Figures S6D** and **S6E**). This result was somewhat consistent with a previous report that commercial anti-TERT antibodies are nonspecific and only detect highly overexpressed, but not low levels of endogenous TERT protein,^92,93^ as was also shown in our western blot analysis of TERT protein overexpression (**Figures S6D** and **S6E**). Due to the lack of suitable anti-TERT antibodies, we employed a telomerase enzyme assay called the telomerase repeated amplification protocol (TRAP),^94^ which measures telomerase activity as a surrogate for TERT protein expression. Since telomerase activity in the SNU-C4 cell line has not been reported, and a cancer cell can maintain telomere length through a telomerase-independent mechanism,^95^ we first confirmed, using TRAP assays, that SNU-C4 cells harbor active telomerase, with telomerase-positive cell lines (HCT116^96^ and Namalwa^97^) and the telomerase-negative U2OS^98^ cell line as controls (**Figure S6F**). Next, we compared telomerase activity between vehicle-treated and FTIN- treated SNU-C4 cells, and observed that cell lysates prepared with FTIN-treated SNU-C4 cells exhibited significantly higher telomerase activity than those with vehicle-treated SNU-C4 cells (**Figure 6F**). These results demonstrate that FTIN-induced *TERT* transcription results in increased telomerase activity (likely through elevated TERT protein expression) in SNU-C4 cells.

Next, we examined the impact of blocking TERT expression (using *TERT*-targeting siRNA: siTERT) on the FTIN-promoted proliferation of SNU-C4 cells. siTERT transfection significantly reduced *TERT* mRNA levels in SNU-C4 cells (**Figure S6G**), and FTIN-promoted cell proliferation was completely abolished in SNU-C4/siTERT cells (**Figure 6G**). These results indicate that the promoting effect of FTIN on CRC cell proliferation requires TERT upregulation.

### *Fn*-mediated promotion of CRC cell proliferation requires TERT

To determine the contribution of *Fn*-mediated AHR activation to *TERT* upregulation, we compared *TERT* mRNA levels between SNU-C4 cells co-incubated with wild-type *Fn*, Δ*tnaA*, or Δ*tnaA*/p*tnaA* strain. Both wild-type *Fn* and the complemented Δ*tnaA*/p*tnaA* strain, but not the Δ*tnaA* mutant, significantly upregulated *TERT* mRNA levels to a similar extent, compared with the media control (**Figure 6H**). We also observed a similar pattern of mRNA expression of *CYP1A1*, used as a control. Next, we tested whether increased TERT expression enhances SNU-C4 cell proliferation. TERT overexpression in SNU-C4 cells resulted in increased telomerase activity (**Figure S6H**) and was sufficient to promote the proliferation of SNU-C4 cells (**Figure 6I**). These results suggest that *Fn*-induced TERT upregulation is necessary and sufficient for the *Fn*-mediated promotion of SNU-C4 cell proliferation.

### FTIN is produced by both *Fusobacterium* and other CRC-associated bacteria and is enriched in human CRC tissues

To gain insights into the clinical relevance of *Fn*-produced AHR-activating metabolites (FTIN, STIN, and TIN), we attempted to detect the *Fn* metabolites in human CRC tissues using LC-MS/MS. To this end, we obtained a total of 30 pairs of human CRC tissues and their matched normal colon tissues originating from 30 CRC patients (**Table S3**). Two independent pieces of cuts from the respective tissue samples were homogenized in 50% methanol, clarified by centrifugation, and subjected to LC-MS/MS analysis. With isotope-labeled FTIN-^13^C_6_ as an internal control, a limit of quantification for FTIN was 0.02 nmol/g tissue, and for both STIN and TIN, 0.08 nmol/g tissue under our analytical conditions. For follow-up analysis of LC-MS/MS data, we used the average values of two measurements. Neither STIN nor TIN was detectable in all tissue samples; in contrast, we were able to detect FTIN at quantifiable levels in most CRC (26/30) and one-third of the normal colon (10/30) tissue samples (**Figure 7A**; **Table S4**). The paired comparison of tissues revealed that FTIN levels are significantly higher in CRC tissues at all clinical stages compared to matched normal colon tissues (**Figure 7B**). In 26 CRC tissues where FTIN could be measured, FTIN levels ranged from 0.02 to 0.96 nmol/g tissue, and the median FTIN level in all 30 CRC tissues was 0.14 nmol/g tissue. These FTIN values in CRC tissues were in sharp contrast with those in normal colon tissues: FTIN levels in 10 normal colon tissues where FTIN could be measured ranged from 0.02 nmol/g tissue to 0.14 nmol/g tissue, and the median FTIN level in all 30 normal colon tissues was zero. These results indicate that FTIN is more prevalent and abundant in clinical CRC tissues than in adjacent normal colon tissues.

**Figure 7.**
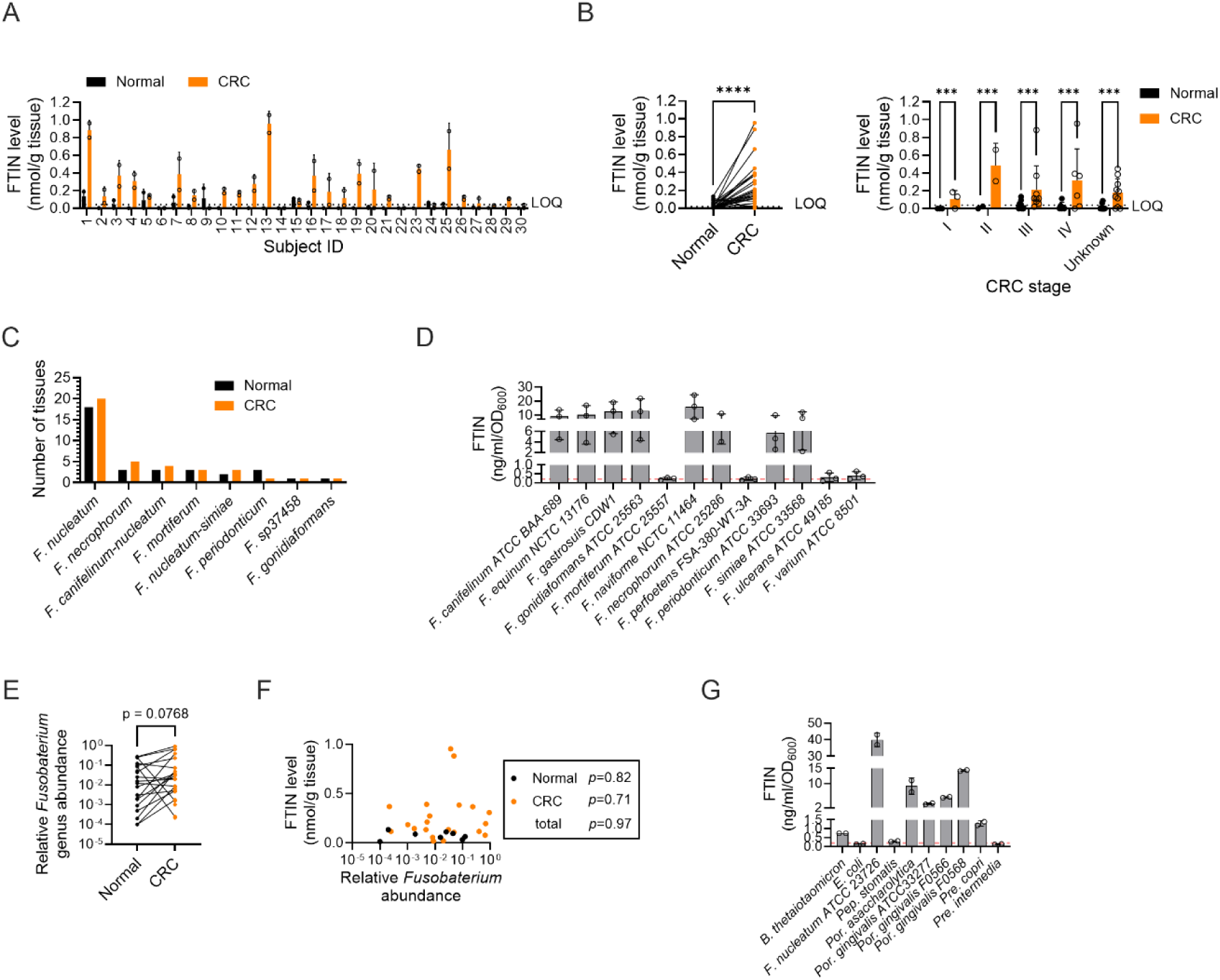
FTIN is enriched in human CRC tissues and is produced by both ***Fusobacterium* and non-*Fusobacterium* species of CRC-associated bacteria** (A) FTIN levels in 30 pairs of human CRC tissues and matching normal colon tissues from 30 subjects. Two pieces of cuts from each tissue were processed for FTIN measurement, and the average value of the two measurements was used for each tissue. The dotted line in (A and B) indicates LOQ for FTIN (0.02 nmol/g tissue). (B) FTIN levels shown in (A) are significantly higher in CRC tissues at all clinical stages (stage I, *n* = 3 pairs; stage II, *n* = 2 pairs; stage III, *n* = 9 pairs; stage IV, *n* = 6 pairs; stage unknown, *n* = 10 pairs). *p*-value was determined by paired t-test (left figure). Statistical significance was assessed using a paired *t*-test (left) and a two-way ANOVA with Šídák’s multiple comparisons test (right): ***, *p* < 0.001; ****, *p* < 0.0001. (C) The number of tissues harboring each *Fusobacterium* species with a relative abundance of ≥ 0.01%. (D) FTIN production by non-*Fn Fusobacteria* (*n* = 3). Respective bacteria were grown anaerobically in BHIc for 24 h, and culture supernatants were collected for FTIN measurement. Data are represented as mean ± SD. The dotted line indicates the LOQ (0.2 ng/ml). (E) Relative abundance of the *Fusobacterium* genus was significantly higher in the CRC tissues than that in the matching normal colon tissues (*n* = 27 pairs for which FTIN levels were above LOQ=0.02 nmol/g tissue). Statistical significance was assessed using a paired *t*- test. (F) No correlation between FTIN levels and the *Fusobacterium* relative abundance in CRC, normal colon, or total (CRC and normal colon) tissues used in (E). The correlation coefficient and associated *p*-value were obtained by Pearson correlation analysis. (G) FTIN production by other non-*Fusobacterium* CRC-associated bacteria (*n* = 2). Respective bacterial species were grown anaerobically in BHIS for 24 h, and their culture supernatants were collected for FTIN measurement. *F. nucleatum* ATCC 23726 was included for comparison. Data are represented as mean ± SD. The dotted line indicates the LOQ (0.2 ng/ml).

Our finding that *Fn*-produced FTIN was significantly more abundant in human CRC tissues suggested an association between *Fn* abundance and FTIN levels. To test this possibility, we profiled bacterial composition in the 30 pairs of CRC-normal colon tissues using 16S rRNA gene amplicon sequencing (16S-seq) (**Table S5**). The total bacterial abundance was more or less similar between CRC and normal colon tissues (**Figure S7A**), and bacterial communities between most CRC and their matched normal colon tissues from the same patients exhibited similar compositions (**Figure S7B**), in line with the previous reports.^22,99^ *Fusobacterium nucleatum* has been the most prevalently detected *Fusobacterium* species in CRC tissues.^15^ Consistently, our 16S-seq data confirmed *F. nucleatum* as being most prevalent in our CRC (20/30) and normal colon tissues (18/30), and found other *Fusobacterium* species in a much smaller number of tissues (1∼5 CRC or normal colon tissues/species) (**Figure 7C**). The minor *Fusobacteria* detected were six known species (*F. canifelinum*, *F. gonidiaformans*, *F. mortiferum*, *F. necrophorum*, *F. peridonticum*, *F. simiae*) and one unknown. We wondered whether these other *Fusobacteria* also produce FTIN. We tested 12 other *Fusobacterium* species and found that eight of them produce significant amounts of FTIN when grown overnight in BHIc media (**Figure 7D**). The *Fusobacteria* newly identified as being FTIN producers included five species (*F. canifelinum*, *F. gonidiaformans*, *F. necrophorum*, *F. peridonticum*, and *F. simiae*) detected in our CRC and normal colon tissues, as well as *F. equinum*^100^ and *F. gastrosuis*^101^ that are known to be animal isolates. *F. mortiferum* was the only non-FTIN producer detected in our tissue samples. The finding of additional FTIN- producing *Fusobacterium* species suggested that the FTIN amounts measured in our CRC and normal colon tissues are likely the summation resulting from the additive action of FTIN- producing *Fusobacteria*. Given this finding, we analyzed our 16S-seq data at the *Fusobacterium* genus level.

*Fusobacterium* was detected (with a relative abundance of 0.01% as a cutoff) in both normal colon (21/30) and CRC (23/30) tissues (**Figure S7C**), consistent with a previous report on the prevalence of *Fusobacterium* in both cancer and normal colon tissues from CRC patients.^102^ In 27 tissue pairs, in which FTIN levels were above the quantification limit (i.e., 0.02 nmol/g tissue) in either 26 CRC or 10 normal colon tissues, the relative abundance of *Fusobacterium* (**Figure 7E**), but not other genera (**Figure S7D**), was significantly higher in CRC *vs*. normal colon tissues. Somewhat unexpectedly, however, when we analyzed the correlation between relative *Fusobacterium* abundance and FTIN levels in 36 tissues (i.e., 26 CRC and 10 normal colon), we observed no correlation, either in CRC or normal colon tissues alone or combined (**Figure 7F**). We reasoned that such a lack of correlation may be due to other FTIN-producing CRC-associated bacteria, in addition to those in the *Fusobacterium* genus. In line with this notion were relatively high levels of FTIN measured in four CRC tissue samples (i.e., subjects 11, 23, 25, and 26) where *Fusobacterium* was not detected: in these tissue four CRC samples, the FTIN levels ranged from 0.12 to 0.66 nmol/g tissue (**Table S4**), with the median FTIN level of 0.31 nmol/g tissue (note that the median FTIN level in all 30 CRC tissues was 0.14 nmol/g tissue). Due to the small sample size, we could not perform statistical analysis to identify bacteria enriched in these five CRC tissues (*vs*. matched normal colon tissues). As an alternative approach, we compiled 30 previously reported CRC-enriched bacteria (22 species from 16 genera)^16,24,25,27,31,32,99,103^ and tested them for FTIN production, with *Fn* as a control (**Table S6**). Remarkably, we found that nine bacteria (in seven species) from five genera (*Bacteroides*, *Escherichia*, *Peptostreptococcus*, *Porphyromonas*, *Prevotella*) produce FTIN at measurable levels ranging from 0.12 to 14.1 ng/ml/OD_600_ (**Figure 7G**). Under our culture conditions, those belonging to the *Porphyromonas* genus (*Por*. *asaccharolytica* and three strains of *Por. gingivalis*) robustly produced FTIN relatively higher than others (*B. thetaiotaomicro*n, *E. coli*, *Pep. anaerobius*, *Pre. copri*, and *Pre. intermedia*) (**Figure 7G**). Although the capacity of FTIN production by these CRC-associated bacteria newly identified as FTIN producers, e.g., under different culture conditions, remains to be thoroughly investigated, these results support the notion that, in addition to *Fusobacterium*, other CRC-associated bacteria may contribute to the FTIN levels detected in CRC tissues.

## DISCUSSION

Our study reveals that the anaerobic bacterium *Fn* produces previously unknown AHR-activating metabolites (FTIN, STIN, and TIN). Before this study, each of the three metabolites was primarily isolated from facultative anaerobic or syntrophic bacteria grown under aerobic conditions. While *Enterococcus faecium* (previously named *Streptococcus faecium*) is the only known bacterium producing STIN,^60,104^ multiple bacteria have been shown to produce either FTIN,^57,58,105–107^ TIN,^108^ or both.^59,109,110^ During the preparation of our manuscript, a study reported the production of FTIN and TIN by *Campylobacter concisus*, and it showed both FTIN and TIN as AHR activators using a sequencing-based reporter assay.^111^ Our study, using the AHR reporter assay, mRNA induction, cell-based AHR binding assay, *in silico* modeling, chemical inhibition of AHR, and genetic perturbations (AHR knockdown and knockout), clearly establishes that all three metabolites are bona fide AHR agonists. This establishment provides a solid basis for investigating the (patho)physiological roles of these metabolites in inter-kingdom interactions between host-commensal (or pathogenic) bacteria, such as FTIN- producers newly identified in this study.

Our study identifies FTIN as the dominant functional entity: it is produced at the highest levels under our culture conditions, binds AHR with the strongest affinity, and is the most potent inducer of canonical AHR signaling (CYP1A1 transcription and activity). Using genetic and pharmacologic perturbations, we have demonstrated that FTIN-induced phenotypes require AHR and proceed via its genomic (ARNT-dependent, DNA-binding) action rather than non-genomic signaling. Functionally, FTIN promotes proliferation, migration, and invasion in a subset of CRC cell lines (those responsive to EGFR blockade, such as SNU-C4), whereas it has no such effects in EGFR blockade-resistant CRC cell lines such as HCT116. Whether FTIN-activated AHR plays any role in the progression of HCT116-like CRC requires further investigation.

A recent study has shown differences in the pathogenic potential of *Fna* clades in CRC; the clade 2 of *Fna* (i.e., *Fna* C2) is more invasive than *Fna* C1 into CRC cells, and *Fna* C2 is predominant over *Fna* C1 and other *Fn* subspecies in CRC tissues compared to adjacent normal colon tissues.^15^ While the existence of the *Fna* C2 clade is still under debate,^112^ our results showed that all tested *Fn* subspecies (i.e., *Fna*, *Fnn*, *Fnp*, and *Fnv*), including *Fna* C2 strains CTI-01, CTI-05, ATCC 51191, and EVAG_002 (also known as 7_1), are robust FTIN producers, suggesting that our findings with the model strain, *Fnn* ATCC 23726, are likely generalizable to other FTIN-producing *Fn* subspecies as part of virulence traits.

Our results demonstrating that *Fn* enhances CRC cell proliferation via FTIN-activated AHR *in vitro* and *in vivo* are in line with two recent studies reporting that *Fn*-mediated AHR activation stimulates the proliferation of esophageal squamous cell carcinoma^113^ and that *Fn* activates AHR in macrophages, which indirectly promotes CRC cell proliferation.^47^ In the former, however, the entity for *Fn*-mediated AHR activation was unknown. In the latter, IPA was proposed to be the *Fn*-produced A HR-activating metabolite. However, we were unable to detect IPA at measurable levels in *Fn* culture, at least under our culture conditions. Our failure to detect IPA in *Fn* cultures is consistent with a previous *in silico* prediction that the genome of *Fn* ATCC 25586 lacks an IPA biosynthetic pathway from L-tryptophan,^54,114^ although we cannot rule out the possibility that *Fn* may produce IPA from other intermediate(s) unavailable in our culture media. One additional study reported that *Fn*-derived formate enhances AHR signaling, promoting Transwell invasion of HCT116 CRC cells, increasing CRC stemness, and driving CRC metastasis,^29^ although the mechanism by which formate enhances the AHR signaling is undefined. By contrast, FTIN, the most potent *Fn*-produced AHR agonist found in this study, did not promote Transwell migration of HCT116 cells under our test conditions, perhaps suggesting that FTIN-activated and formate-enhanced AHR signaling may elicit distinct cellular programs in CRC cells. The combined and context-dependent contributions of FTIN and formate to the *Fn*-mediated promotion of CRC progression remain to be determined. We find indole as the entry point of FTIN/STIN/TIN biosynthesis. Deleting *tnaA* eliminates indole and abolishes production of all three metabolites, while genetic and chemical complementation restores it. Consistently, the *tnaA* mutant fails to activate AHR, is unable to promote proliferation, migration, and invasion of the SNU-C4 cell line *in vitro*, and loses the ability to enhance SNU-C4 tumor growth *in vivo*, despite displaying wild-type growth and attachment to and invasion into CRC cells. These results provide a reasonable rationale to explore small-molecule TnaA inhibitors as adjuvants, especially in EGFR blockade-responsive CRC.

Our study, using gene expression profiling and targeted validation, uncovers TERT as a key downstream effector of FTIN-activated AHR signaling, which explains the cell line-dependent proliferative response to FTIN. FTIN upregulates *TERT* mRNA via the AHR/ARNT pathway and increases telomerase activity. Conversely, TERT knockdown eliminates the FTIN-induced promotion of CRC cell proliferation, while TERT overexpression is sufficient to boost CRC cell proliferation. These findings support a model in which FTIN-activated AHR drives a TERT-dependent program that fuels growth in subtypes of CRC cell lines such as SNU-C4. On the other hand, the absence of FTIN effects in HCT116 and HT-29 CRC cell lines, despite intact AHR reporter responses, suggests that the transcriptional output of AHR signaling is dependent on cell state (likely defined by distinct sets of oncogenic mutations). The genetic contexts of CRC cell lines and the regulatory molecular mechanism that couples AHR activation to TERT induction, promoting cell proliferation, remain to be determined.

TERT has been implicated in several features of cancer hallmarks, including enhanced proliferation and invasion/metastasis, as well as resistance to apoptosis.^115–118^ It has also been known that TERT promotes cancer progression via mechanisms that are both dependent and independent of its telomerase function.^115–118^ It remains to be determined whether TERT’s telomerase-dependent and/or telomerase-independent functions mediate FTIN-driven CRC cell proliferation.

We demonstrate that FTIN production is not only conserved across multiple *Fn* subspecies and strains, as well as other *Fusobacterium* species, but also detectable in additional CRC-associated genera under our test conditions. Moreover, in patient samples, FTIN is significantly enriched in CRC tissues relative to matched normal colon tissues, supporting its *in vivo* relevance. Notably, FTIN levels do not correlate with *Fusobacterium* relative abundance at the genus level across CRC and normal colon tissues. Two non-exclusive scenarios may explain these results. First, FTIN production by a community of CRC-associated bacteria: multiple intratumoral bacteria contribute to the FTIN pool in CRC tissues. Second, the regulation of FTIN production in intratumoral bacteria under *in vivo* conditions. In the latter scenario, several physical and/or chemical factors may influence FTIN production/pool, including intratumoral bacterial metabolism under *in vivo* conditions (e.g., oxygen levels and nutrient availability), the spatial localization of intratumoral bacteria (e.g., intracellular or extracellular), metabolic cross-feeding between intratumoral bacteria and/or between a host cell type (e.g., immune or epithelial) and intratumoral bacteria, and host clearance. Addressing these physical and chemical factors in the CRC microenvironment will further clarify the contribution of FTIN- activated AHR signaling to CRC progression and provide a better gauge for the potential utilization of FTIN as a biomarker in CRC.

### Limitations of the study

While we utilized the *tnaA* mutant as evidence that FTIN-induced AHR activation is critical for *Fn*-mediated promotion of CRC cell proliferation *in vitro* and *in vivo*, as well as cell migration and invasion *in vitro*, the CRC phenotypes observed could reflect broader perturbations in the *tnaA* mutant. The TnaA enzyme converts L-tryptophan to indole, as well as ammonia and pyruvate. Deleting *tnaA* eliminates not only FTIN/STIN/TIN production but also the three immediate products of L-tryptophan. Therefore, our results obtained with the *tnaA* mutant do not distinguish the (potential) role of indole, ammonia, and/or pyruvate from that of FTIN in *Fn* interaction with CRC cells. To clarify this aspect, it will be necessary to elucidate the FTIN/STIN/TIN biosynthetic pathway and construct and test a mutant that is specifically defective in FTIN, STIN, and/or TIN production.

## RESOURCE AVAILABILITY

### Lead contact

Further information and requests for resources and reagents should be directed to the lead contact, Hyunyoung Jeong

## Materials availability

The plasmids, bacterial strains, and cell lines generated in this study will be made available upon request.

## Data and code availability

- 16S rRNA gene amplicon sequencing and RNA sequencing data are publicly available online at the NCBI database (https://www.ncbi.nlm.nih.gov/bioproject) under BioProject accession numbers SRA: PRJNAxxxxxxx (RNA-seq data) and PRJNAxxxxxxx (16-seq data).
- This paper does not report original code.
- Additional information required to reanalyze the data reported in this paper is available from the lead contact upon request.

## Supporting information

Tabls S3

Tabls S5

## ACKNOWLEDGMENTS

This work was partially funded by NIH/NCCIH R21AT011391 (HJ and HL) and the Purdue Institute for Cancer Research (HJ). L.J.D. was supported by the T32 AT007533 training grant. We thank Cynthia L. Sears at Johns Hopkins Bloomberg School of Public Health and E. Allen-Vercoe at the University of Calgary for providing *Fusobacterium nucleatum* strains. We are grateful for access to the Bruker Avance 600 MHz NMR at the UIC Center for Structural Biology. We thank Dr. J. Bisson for assistance with Bruker ImpactII Qq-TOF experiments at the Institute for Tuberculosis Research at UIC.

## AUTHOR CONTRIBUTIONS

Conceptualization, K-J.W., H.L., and H.J.; investigation, K-J.W., P-R.J., L.D., J.O., A.T.A, W-H.W, H. K., S.M.U, D.B., B-S. J., I.A.M., C.X., J.L., and G.H.P.; resources, C.W., B-S.J. and G.H.P.; writing – original draft, K-J.W., P-R.J., L.D., J.O., B-S.J., G.H.P., H.L., and H.J.; writing – review & editing, H.L. and H.J.; supervision, H.L. and H.J.

## EXPERIMENTAL MODEL AND SUBJECT DETAILS

### Bacterial strains and growth conditions

All strains used and their source of origin are listed in the KEY RESOURCES TABLE. All anaerobic bacterial strains used were maintained as frozen glycerol stocks at -80°C, and media (agar and broth) were pre-reduced overnight in an anaerobic chamber (with a gas mixture of 5% H_2_, 5% CO_2_, and 90% N_2_; Anaerobe Systems, Morgan Hill, CA) before use. Bacterial strains were grown in pre-reduced BHIc (brain heart infusion medium containing 0.03% L-cysteine); BHIS (BHIc supplemented with 5 mg/mL of hemin and 0.5 mg/mL of vitamin K_3_); or M9CT (M9 medium: M9 salts, 0.4% glucose, 2 mM MgSO_4_, and 0.1 mM CaCl_2_; supplemented with 0.03% L-cysteine and 1% tryptone). For preparation of bacterial cultures in an experiment, a bacterial strain from frozen stock was streaked on BHIc agar, and a freshly grown colony was inoculated into liquid media and grown overnight at 37°C under anaerobic conditions. Typically, fresh bacterial cultures were prepared by diluting overnight cultures 50 to 100-fold in appropriate fresh broth and growing overnight or for 2-3 days at 37°C under anaerobic conditions and used for experiments. For cloning, *E. coli* DH5α and TOP10 strains were used, and they were grown in Luria-Bertani (LB) media.

### Cell lines

Human hepatoma HepG2 and human colorectal cancer (CRC) cell lines (LIM1215, HCT-8, H508, SNU-C4, HCT116, and HT-29) were cultured in Dulbecco’s modified Eagle’s medium (DMEM). Murine hepatoma Hepa1c1c7 cells were cultured in Minimum Essential Medium α (MEMα). For maintenance of cell lines, all media were supplemented with 10% (v/v) fetal bovine serum (FBS, GeminiBio), penicillin (100 units/ml), and streptomycin (100 µg/ml), and all cell lines were cultured in a humidified incubator (Nuaire, ThermoFischer Scientific) at 37°C with 95% air and 5% CO_2_.

### Animal experiments

Athymic nude mice of 6 weeks old (NU/J, Stock number: 002019) were purchased from Jackson Laboratory. Mice were housed with gamma-irradiated diet (Teklad 2918), autoclaved water, and corncob bedding under specific pathogen-free conditions, maintained in 12 h light/12 h dark cycle, and used for the experiment after one week of acclimation. All mouse experiments were performed using protocols (#2011002089) approved by Purdue University Animal Care and Use Committee.

### Human colorectal cancer and normal colon tissues

A total of 30 pairs of fresh-frozen colorectal cancer and matched normal colon tissues were purchased from the Indiana University Simon Cancer Center Komen Tissue Bank, with informed consent approved by the Komen Tissue Bank Institutional Review Board. The study cohort consisted of patients with histologically confirmed primary colorectal adenocarcinoma who underwent surgical resection in Indiana University Health Hospitals. Pathological assessment of all tissue samples was performed by a resident pathologist for the confirmation of cancerous and normal states of tissues. Clinical data, including age, sex, tumor location, stage, and histology, were provided by Komen Tissue Bank. Upon procurement, frozen tissue samples were stored in liquid nitrogen vapor phase until use.

## METHOD DETAILS

### Preparation of bacterial culture extracts

Bacterial strains tested for induction of *Cyp1a1* promoter activity or for measurement of FTIN production were grown in BHIc (Figure 1A and 1D) or in BHIS (Figure 7D and 7G) at 37°C for 1-3 days under anaerobic conditions. For other experiments involving strain *F. nucleatum* ATCC 23726 (*Fn*), extracts were prepared with cultures grown overnight in M9CT at 37°C under anaerobic conditions unless otherwise mentioned. To prepare the organic extracts of supernatants, cultures were centrifuged (4,000*g* × 15 min) to remove cells, and culture supernatants were collected and extracted with an equal volume of ethyl acetate. The organic layer was transferred and dried *in vacuo* using a Vacufuge (Eppendorf). Dried extracts were dissolved in dimethyl sulfoxide (DMSO) at 100× concentration. To prepare the extracts of *Fn* cells, cell pellets were extracted with 100% MeOH for 16 h. After clarification by centrifugation (4800 *g* for 10 min), MeOH extracts were collected and dried *in vacuo* using Vacufuge and reconstituted in DMSO at 100× concentration, relative to the original culture volume. Final extracts in DMSO were tested at 1:1,000 dilution, equivalent to 10% of the total culture volume.

### Preparation of chemically competent and electrocompetent bacterial cells

Chemically competent cells of *E. coli* were prepared as described previously (Hanahan, 1983; PMID 6345791). Electrocompetent cells of *Fusobacterium nucleatum* ATCC 23726 (*Fn*) were prepared as described previously (Peluso *et al*., 2020; PMID 32539234), with a slight modification. *Fn* cells were grown to mid-log phase (OD_600_ ∼0.6) in 10 mL of BHIc media at 37°C under anaerobic conditions. After the culture was kept on ice for 20 min, cells were harvested by centrifugation (4800*g* × 10 min at 10°C), washed twice with ice-cold 10% glycerol, and resuspended in 1 mL of ice-cold 10% glycerol. Aliquots (100 µL) of the electrocompetent cells were snap-frozen in liquid nitrogen and stored at -80°C before use.

### Cytotoxicity (MTT) assay

Cytotoxicity of bacterial extracts against HepG2 cells was determined by the 3-(4,5- dimethylthiazol-2-yl)-2,5-diphenyltetrazolium bromide (MTT) assay (van Meerloo *et al*., 2011; PMID 21516412). HepG2 cells were seeded in 96-well plates (5×10^3^ cells/well) in DMEM medium supplemented with 10% FBS. After 24 h incubation, cells were treated with organic extracts of bacterial cultures in serum-free DMEM medium. After 24 h treatment, MTT reagent (5 mg/mL in PBS) was added to each well (15 µL) and the reaction was incubated at 37°C for 3 h. After the medium was aspirated, DMSO (50 µL/well) was added to dissolve formazan, and the absorbance was measured at 570 nm.

### Purification of *Fn* metabolites and structure determination

Complete information about the metabolite purification and structure validation was described in Supplemental Texts that include Supplemental Results, Supplemental Method Details, Supplemental References, Supporting Table 1, and Supporting Figures (S1-S16).

### Syntheses of FTIN, isotope-labeled FTIN, STIN, and TIN

FTIN (>96% purity) and isotope labelled FTIN (FTIN-^13^C_6_, >99% purity) were custom synthesized by WUXI, and STIN (>95% purity) by KareBay Bio. FTIN and TIN were also in-house synthesized. For in-house synthesis of FTIN and TIN, we searched in CAS SciFinderⁿ, a chemical compound database, and retrieved previous reports on the chemical synthesis of FTIN and TIN (274 and 35 papers, respectively). We selected synthetic methods for which precursor chemicals are readily available. In brief, to prepare FTIN, we employed a strategy for C2-quaternary indolin-3-ones, the nitrogen dioxide-mediated oxidative trimerization of indoles (Xue *et al*., 2014). FTIN was obtained from a reaction of indole with methanesulfonic acid and sodium nitrite in pyridine in an air atmosphere. To prepare TIN, we employed a reaction of isatin with two equivalents of indole in the presence of *p*-toluenesulfonic acid in dichloromethane at room temperature (Yu *et al*., 2014). Both FTIN and TIN were obtained with >95% purity. ^1^H NMR spectra of custom and in-house synthesized FTIN, isotope-labeled FTIN, STIN, and TIN were shown as Supporting Figures (S2-S6) in Supplemental Texts.

### Quantification of FTIN, indole, IPA, STIN, and TIN

To determine the concentrations of FTIN, TIN, STIN, and indole in the culture of wild-type *Fn*, Δ*tnaA*, or Δ*tnaA*/p*tnaA* strains, each strain was grown in M9CT broth at 37°C under anaerobic conditions. Cultures were sampled at the indicated time points, and culture supernatant was collected by centrifugation (4800*g* × 10 min). After adding FTIN-^13^C_6_ as the internal standard (at 10 ng/mL) to samples for FTIN, TIN, or STIN measurement, the culture supernatant was mixed and extracted with an equal volume of ethyl acetate. The organic phase was separated by centrifugation (4800*g* × 1 min), and the upper layer was transferred to a fresh tube and dried *in vacuo* using Vacufuge (Eppendorf). Dried extracts were then reconstituted with methanol and subjected to HPLC-UV analysis for indole and LC-MS/MS analysis for FTIN, IPA, STIN, and TIN. Samples for other *Fusobacterium* species and non-*Fusobacterium* CRC-associated bacteria were prepared similarly and analyzed for FTIN.

Indole measurement: Organic extracts were diluted in 50% methanol (v/v) in water and filtered (Cytiva, 4 mm nylon syringe filter, 0.2 µm pore size). Analysis of the filtrates (40 µL) was conducted using an Agilent 1100 series HPLC system (Agilent, Santa Clara, CA) coupled with a G1314A UV detector. Chromatographic separation was achieved using a C_18_ column (Waters, XTerra MS, 5 µm, 4.6 × 250 mm). The mobile phase consisted of 10 mM ammonium acetate in water (solvent A) and acetonitrile (solvent B) and was run as follows: 0-25 min (15- 90% B), 25-34 min (90% B), 34-35 min (90-15% B) and 35-45 min (15% B) at a flow rate of 1 mL/min. Eluates were monitored at 274 nm wavelength. Indole standard solutions (ranging from 182.75 ng/mL to 11.72 µg/mL) were prepared in 50% methanol (v/v) in water and used to obtain a standard curve run for each batch of samples, and indole concentrations in respective samples were calculated from the standard curve.

FTIN measurement: Organic extracts were diluted in 50% acetonitrile (v/v) in water containing 0.1% formic acid and filtered (Cytiva, 4 mm nylon syringe filter, 0.2 µm pore size). Analysis of the filtrates (5 μL) was conducted using a Sciex 4500 QTRAP mass spectrometer equipped with an electrospray ionization source operating in positive ion mode, connected to a Prominence UFLC HPLC system (Shimadzu?). Chromatographic separation was achieved using a C_18_ column (Waters, Atlantis T3, 3 µm, 3 × 100 mm) operating at a flow rate of 0.25 mL/min. The mobile phase consisted of 0.1% (v/v) formic acid in water (solvent A) and acetonitrile (solvent B): 0-3.5 min (30-90% B), 3.5-7.5 min (90% B), 7.5-12 min (30% B). FTIN-^13^C_6_ was used as the internal standard to obtain the calibration curve for the quantification of FTIN. The detection and quantification of analytes were accomplished by multiple reaction monitoring (MRM) of *m/z* 364.200/219.100 for FTIN; and 370.300/222.000 for FTIN-^13^C_6_. FTIN standard solutions (ranging from 0.01 to 100 ng/mL) were prepared in 50% acetonitrile (v/v) in water containing 0.1% formic acid and internal standard (FTIN-^13^C_6_) and used to obtain a standard curve run for each batch of samples. FTIN concentrations in respective samples were calculated from the standard curve.

TIN measurement: Samples were prepared as described for FTIN and analyzed by the LC-MS/MS system operating in negative ion mode and chromatographic separation method as described for FTIN measurement. The detection and quantification of analytes was accomplished by MRM of *m/z* 362.000/244.900 for TIN. TIN standard solutions (ranging from 0.78 to 100 ng/mL) were prepared in 50% acetonitrile (v/v) in water containing 0.1% formic acid and internal standard (FTIN-^13^C_6_) and used to obtain a standard curve run for each batch of samples. TIN concentrations in respective samples were calculated from the standard curve.

STIN measurement: Organic extracts were diluted in 50% acetonitrile (v/v) in water containing 10 mM ammonium acetate and filtered. The filtrates (5 μL) were analyzed by the LC-MS/MS system as described for FTIN measurement. Chromatographic separation was achieved using a C_18_ column (Waters, Atlantis T3, 3 µm, 3 × 100 mm). The mobile phase consisted of 10 mM ammonium acetate in water (solvent A) and acetonitrile (solvent B) and was run with the chromatographic program used in FTIN measurement. The detection and quantification of analytes was accomplished by MRM of *m/z* 336.400/160.200 for STIN. STIN standard solutions (ranging from 0.78 ng/mL to 100 ng/mL) were prepared in 50% acetonitrile (v/v) in water containing 10 mM ammonium acetate and internal standard (FTIN-^13^C_6_) and used to obtain a standard curve run for each batch of samples. TIN concentrations in respective samples were calculated from the standard curve.

IPA measurement: Culture extracts were prepared as described above with indolepropionic acid-*d*_2_ (IPA-*d*_2_ at 10 ng/ml) spiked as the internal standard. Organic extracts were diluted in 50% acetonitrile (v/v) in water containing 0.1% formic acid and filtered (Cytiva, 4 mm nylon syringe filter, 0.2 µm pore size). Analysis of the filtrates (5 μL) was conducted using the LC-MS/MS used for FTIN measurement. Chromatographic separation was performed on a C_18_ column (Waters, Xterra MS, 3.5 µm, 2.1 × 50 mm) at a flow rate of 0.3 mL/min. Mobile phase consisted of 0.1% (v/v) formic acid in water (solvent A) and acetonitrile (solvent B): 0-0.9 min (10% B), 0.9-3.6 min (10-35% B), 3.6-5 min (35% B), 5-6.5 min (35-90% B), 6.5-8 min (90% B), 8-8.1min (90-10% B), 8.1-11 min (10% B). IPA-*d*_2_ was used as an internal standard to obtain the calibration curve for the quantification of IPA. The detection and quantification of analytes were accomplished by MRM of *m/z* 190.100/130.200 for IPA; and 192.000/130.200 for IPA-*d*_2_. IPA standard solutions (ranging from 0.39 ng/mL to 100 ng/mL) were prepared in 50% acetonitrile (v/v) in water containing 0.1% formic acid and internal standard (IPA-*d*_2_) and used to obtain a standard curve run for each batch of samples. IPA concentrations in respective samples were calculated from the standard curve.

### Luciferase-based AHR reporter assay

Cells were seeded in 48-well plates at a density of 5×10^4^ cells/well. After 24 h of incubation in DMEM medium supplemented with 10% FBS, cells were transfected with both pGL2-Cyp1a1 (Firefly) and pRL-TK (Renilla) plasmids using FuGENE® HD transfection reagent for 24 h and treated with vehicle control, a compound, bacterial culture extracts, or a bacterial strain at a multiplicity of infection (MOI) of 300 in serum-free medium for 4 h or 24 h. For the inhibition of AHR activity with an AHR antagonist, cells were pretreated with CH223191 (10 µM) or SR1 (1 µM) in serum-free medium for 1 h before treatment. Luciferase activities were measured with the Dual-Luciferase® Reporter Assay System using a microplate reader (BioTek). *Cyp1a1* promoter activity was calculated by normalizing Firefly luciferase activity to Renilla luciferase activity.

### Luciferase-based β-catenin reporter assay

Cells were seeded in 48-well plates at 5×10^4^ cells/well. After 24 h of incubation in DMEM medium supplemented with 10% FBS, cells were transfected with M50 Super 8× TOPFlash (Firefly) or M51 Super 8× FOPFlash (Firefly) together with pRL-TK (Renilla) plasmids using FuGENE® HD transfection reagent for 24 h and treated with vehicle control or a compound in serum-free medium for 24 h. Luciferase activities were measured with the Dual-Luciferase® Reporter Assay System using a microplate reader (BioTek). Firefly luciferase activity is normalized to Renilla luciferase activity.

### Luciferase-based HIF-1 reporter assay

Cells were seeded in 48-well plates at 5×10^4^ cells/well. After 24 h of incubation in DMEM medium supplemented with 10% FBS, cells were transfected with both HRE-luciferase (Firefly) and pRL-TK (Renilla) plasmids using FuGENE® HD transfection reagent for 24 h and treated with vehicle control, compounds in serum-free medium for 4 h or 24 h. Luciferase activities were measured with the Dual-Luciferase® Reporter Assay System using a microplate reader (BioTek). Firefly luciferase activity is normalized to Renilla luciferase activity.

### Determination of *TERT* mRNA stability

Cells were seeded in 6-well plates at 5×10^4^ cells/well and cultured in DMEM medium supplemented with 10% FBS for 24 h. The media were aspirated, and cells were treated with FTIN (100 nM) or vehicle control in serum-free medium. After 24 h of incubation, cells were cultured in serum-free media containing actinomycin D (5 µg/ml), with or without FTIN (100 nM), and harvested for RNA isolation at time zero and at intervals up to 9 h. Relative mRNA levels were determined by qRT-PCR as described in the qRT-PCR section.

### CYP1A enzyme assay

The assay for the measurement of ethoxyresorufin-*O*-deethylase (EROD) activity of CYP1A enzymes was performed as described previously (Schiwy *et al*., 2015; PMID 26448361). In brief, HepG2 cells were seeded in 96-well plates (100 µL/well) at a density of 3×10^5^ cells/mL. After 16 h of incubation, cells were treated with FTIN (1 µM), STIN (1 µM), TIN (1 µM), TCDD (3 nM), or 3MC (1 µM). At the indicated time point posttreatment, the medium was replaced with fresh medium containing 7-ethoxyresorufin (1 µM). After incubation at 37°C for 30 min, the reaction was stopped by adding 75 µL of ice-cold methanol. An aliquot (50 µL) of the sample was then transferred to a black 96-well plate, and the fluorescence intensity of resorufin (excitation at 558 nm, emission at 593 nm) was measured with a plate reader (BioTek). Protein concentration of samples was determined using the BCA protein assay kit (Pierce) and used for normalization.

### Cell-based AHR photoaffinity ligand (PAL) competition assay

Human HN30 (pharyngeal squamous cell carcinoma) cells were seeded in 12-well plates (1 mL/well) at a density of 5×10^4^ cells/mL, and cultured for 48 h in DMEM/F12 basal media (Sigma) supplemented with 10% FBS, 25 mM HEPES, pH 7.4, 100 U/mL penicillin and 100 µg/mL streptomycin. Culture medium was aspirated, and cells were washed with 1 mL Dulbecco’s phosphate-buffered saline (PBS; Sigma) followed by the addition of 0.5 mL of 1× Hank’s balanced salt solution (HBSS) supplemented with 5 mg/mL bovine serum albumin (fraction V). Cells were allowed to equilibrate for 30 min at 37°C before 15 min incubation with the indicated test competitors (fusotrisindoline, streptindole, and trisindoline) at a final concentration of 10 µM. DMSO, 10 µM estradiol, and 100 nM indirubin were included as vehicle, negative, and positive controls, respectively. Following 15 min preincubation, cells were treated with 2 pmol (4 nM) 2-azido-3-[^125^I]-iodo-7, 8-dibromobenzo-*p*-dioxin AHR photoaffinity ligand (PAL) for an additional 30 min at 37°C. Binding media were aspirated, 0.5 mL PBS was added to each well, and the cells were exposed to two 15 W UV lamps (λ > 302 nm) for 4 min, 8 cm from the UV source to facilitate covalent PAL-AHR cross-linkage. Following UV irradiation, PBS was removed, and the cells incubated on ice for 10 min with 0.1 mL/well lysis buffer comprising 25 mM MOPS, 2 mM EDTA, 0.02% (v/v) sodium azide, 10% (v/v) glycerol, 20 mM sodium molybdate, pH 7.28 supplemented with 1% (v/v) Igepal CA630, and Roche complete Mini protease inhibitor cocktail. Lysates were transferred to 1.5 mL tubes, centrifuged at 18,000*g*, 4°C for 20 min. The supernatants were resolved on 8% polyacrylamide-tricine-SDS gels and transferred to the PVDF membrane. Transfer efficiency was determined through Ponceau Red staining. Specific PAL labeled-AHR bands were visualized through autoradiography and subsequently quantified by phospho-imaging densitometry. AHR levels in each lane were determined by probing with sc-5579, rabbit anti-AHR (Santa Cruz) at 1:1,000 dilution, followed by HRP-conjugated goat anti-rabbit IgG (Jackson ImmunoResearch) at 1:2,000 dilution. Specific antibody detection was achieved through HRP-dependent enhanced chemiluminescence (Cytiva-Amersham) using a 1-min incubation time and 15-sec exposures to Biomax MS film (Carestream). Specific AHR bands were subsequently quantified by densitometry (ImageJ).

### *In silico* Modeling of *Fn* metabolites (FTIN, STIN, and TIN) with human AHR

The human AHR model was built from the cryo-EM structure of the indirubin-bound Hsp90- XAP2-AHR complex (PDB ID: 7ZUB). Molecular docking was performed using the Glide software (2024-4 release, Schrödinger, Inc.) (PMID 17034125; 15027866). FTIN, STIN, and TIN were docked to the indirubin-binding pocket, and the Glide XP scores were calculated to compare their binding affinity.

### Sulforhodamine B (SRB) assay for CRC cell proliferation

Cell proliferation was determined with the sulforhodamine B (SRB) assay as previously described (Vichai et al., 2006; PMID 17406391). CRC cells were seeded into 48-well plates at 2×10^4^ cells/well in DMEM medium supplemented with 10% FBS. After 24 h incubation, cells were treated with FTIN (100 nM), TCDD (1 nM), or *Fn* bacterial cells (at an MOI of 300) in serum-free medium. Medium containing FTIN or *Fn* was renewed every 24 h. For the inhibition of AHR activity using an AHR antagonist, cells were pretreated with CH223191 (10 µM) or SR1 (1 µM) in serum-free medium for 1 h before compound or *Fn* treatment. In the case of small interfering RNA (siRNA) for gene silencing, cells were transfected with an siRNA for 24 h before treatment. At the indicated time point after treatment, cells were fixed with 5% trichloroacetic acid solution at 4°C for 1 h, washed with water three times, and air-dried. Cells were then stained with 0.057% SRB solution for 30 min and washed with 1% acetic acid solution three times to remove unbound dye. Protein-bound SRB was eluted with 10 mM Tris base (pH 10.5), and absorbance was measured at 530 nm.

### Transwell assay for CRC cell migration and invasion

Cells were grown to 50-80% confluency in DMEM supplemented with 10% FBS. For the migration assay, cells were trypsinized and resuspended in serum-free medium supplemented with 1 mg/ml BSA, followed by the addition of FTIN (100 nM) or *Fn* cells (MOI 300). Cells (200-400 µL) were then seeded at a density of 0.5-1×10^5^ cells/well into the upper chamber (8.0 µm pore size transwell insert, Corning). The lower chamber was filled with medium containing 10% FBS to serve as a chemoattractant. After 24 h incubation, cells in the upper chamber were fixed with 4% formaldehyde for 15 min at room temperature. Non-migrated cells were gently removed using cotton swabs. Migrated cells were stained with 0.1% crystal violet in PBS for 10 min, followed by washing with PBS and air-drying. Bound crystal violet was eluted using 33% acetic acid, and absorbance was measured at 590 nm using a plate reader (Biotek). For the invasion assay, Matrigel-coated transwell inserts (Corning) were used, and all procedures were performed as described for the migration assay.

### Quantitative real-time (qRT)-PCR

Total RNA was isolated from CRC cells using Trizol (Invitrogen) and was reverse transcribed into cDNA using High-Capacity cDNA Reverse Transcription Kit (Applied Biosystems). Reactions were carried out in 96-well PCR plates using FastStart Universal Probe Master Mix (Roche). Each reaction was performed in duplicate in a final volume of 10 µL with PrimeTime probes (Integrated DNA Technologies). Amplifications were performed with the following thermal cycling conditions: 1 cycle at 50°C for 2 min, 1 cycle at 95°C for 10 min, followed by 40 cycles of 95°C for 15 sec, 60°C for 1 min. The qPCR probes are listed in the KEY RESOURCES TABLE. Target mRNA levels were expressed as values normalized to those of *ACTB* or *Actb* using the 2^-ΔΔct^ method (Livak and Schmittgen, 2001; PMID 11846609).

### Western blot

Cells were washed with PBS and lysed in RIPA buffer (50 mM Tris, pH 7.4, 150 mM NaCl, 0.25% deoxycholic acid, 1% NP-40, and 1 mM EDTA) containing protease and phosphatase inhibitor cocktails (Roche). Protein concentration was determined using the BCA protein assay kit (Pierce). Cell lysates with sample buffers (50 mM Tris-HCl, pH 6.8, 10% glycerol, 2% SDS, 5% 2-mercaptoethanol, 0.04% bromophenol blue) were boiled for 5 min. An aliquot of cell lysates (10-30 µg) was separated by sodium dodecyl sulfate-polyacrylamide gel electrophoresis (SDS-PAGE) and transferred to polyvinylidene difluoride (PVDF) membrane. Membranes were incubated with 5% skim milk in Tris-buffered saline with 0.05% Tween 20 (TBST) for 1 h and incubated with primary antibody at 4°C overnight. Membranes were washed with TBST and incubated with a secondary antibody conjugated to Horseradish peroxidase (HRP) for 1 h. After washing with TBST, proteins on membranes were visualized with chemiluminescent substrate (luminol) using the Azure imaging system. All primary and secondary antibodies used are listed in the KEY RESOURCES TABLE.

### Determination of L-tryptophan auxotrophy of *Fusobacterium nucleatum*

Bacterial strains were grown overnight in BHIc medium. A portion of the overnight culture was streaked on M9 agar containing EZ solution (Teknova) with complete amino acids or EZ solution without L-tryptophan for 48 h. The image was captured with the Azure imaging system (Azure Biosystems).

### Construction of a Δ*fap2* mutant

Primers used for cloning are listed in Table S7. A Δ*fap2* mutant in wild-type *Fn* ATCC 23726 was constructed using a HicA toxin-based mutagenesis (Gc *et al*., 2023; PMID 37039662). Upstream and downstream regions (∼1.2 kb each) of the *fap2* gene were PCR-amplified with genomic DNA as template and primer pairs (UP_Fn-fap2_F/UP_Fn-fap2_R and DN_Fn- fap2_F/DN_Fn-fap2_R). A linearized backbone plasmid was prepared by PCR amplification with pBCG02 as template with a pair of primers (pCWU6-F/pCWU6-R). The two PCR-amplified DNA fragments and linearized backbone plasmid were ligated using In-Fusion Snap Assembly Kit (Takara), and the ligates were transformed into chemically competent *E.coli* DH5α cells. Transformants were selected overnight on Luria-Bertani agar (Sigma) containing chloramphenicol (25 µg/mL). Plasmid pBCG02-Δ*fap2* was isolated from two independent transformants, and its nucleotide sequence was confirmed by whole plasmid sequencing (Genewiz). The sequence-verified pBCG02-Δ*fap2* plasmid (∼36 µg) was introduced to *Fn* ATCC 23726 electrocompetent cells by electroporation (2.5 kV, 200 Ω, 25 µF). Integration of the plasmid into the chromosome was selected on BHIc agar containing thiamphenicol (5 µg/mL). A colony of a transformant was then cultured in BHIc broth for 16 h for the second recombination, and an aliquot of the overnight culture was spread on BHIc agar containing 2 mM theophylline for up to 3 days. Theophylline-resistant clones were confirmed for the deletion of the *fap2* gene by colony PCR with a pair of primers (fap2-CK-F/fap2-CK-R). A Δ*fap2* deletion mutant obtained was used as a control impaired for *Fn* attachment to and invasion into CRC cells.

### Construction of a Δ*tnaA* mutant, a complementation plasmid (p*tnaA*), and a Δ*tnaA*/p*tnaA* strain

Primers used for cloning are listed in Table S7. The construction of a Δ*tnaA* mutant in wild-type *Fn* ATCC 23726 (*Fn*) was reported previously (Gc *et al*., 2023; PMID 37039662). To create a complementation plasmid (p*tnaA*) expressing the *tnaA* gene of *F. nucleatum* ATCC 23726, a DNA fragment containing the *tnaA* gene with its own promoter (2,038 bp) was PCR-amplified with the genomic DNA of *F. nucleatum* ATCC 23726 as the template and a pair of primers (Fn-tnaAB-pCWU6-F/Fn-tnaAB-pCWU6-R). To prepare a backbone plasmid, the pCWU6 plasmid was double-digested with *Sal*I and *Bam*HI and gel-purfied. The *tnaA*- containing DNA fragment and backbone plasmid were ligated using the In-Fusion Snap Assembly Kit (Takara Bio). The ligates were then transformed into chemically competent cells of *E. coli* TOP10 strain, and transformants were selected overnight on LB agar (Sigma) containing chloramphenicol (25 µg/ml). Plasmid was isolated from two independent transformants, and its nucleotide sequence was confirmed by whole plasmid sequencing (Eurofins).

To obtain a complemented Δ*tnaA*/p*tnaA* strain, a sequence-verified p*tnaA* plasmid was transformed into the Δ*tnaA* strain by electroporation (2.5 kV, 25 µF, 200 Ω), and transformants were selected overnight on BHIc agar (BD) containing thiamphenicol (5 µg/ml).

### Determination of FTIN production in the Δ*tnaA* mutant supplemented with indole

The Δ*tnaA* strain was grown overnight in BHIc at 37°C under anaerobic conditions. Half of the overnight culture was heat-treated at 95°C for 10 min. After cooling at room temperature, indole at a final concentration of 500 µM or vehicle (DMSO) was added to heat-inactivated and intact overnight cultures, respectively. After 4 h of incubation at 37°C under anaerobic conditions, culture supernatants were collected by centrifugation (4800*g* × 10 min) and processed as described in the FTIN measurement.

### *Fn* attachment and invasion assays

*Fn* attachment to and invasion into CRC cells were performed as previously described (Rubinstein et al., 2013; PMID 23954158) with a slight modification. SNU-C4 CRC cells were seeded in 24-well plates (500 µL/well) at a density of 6×10^5^ cells/mL in DMEM supplemented with 10% FBS. After 24 h of incubation, cells (at ∼100% confluency) were treated with *Fn* bacterial cells (at an MOI of 5) in serum-free DMEM medium. For the attachment assay, SNU- C4 CRC cells were incubated with bacterial cells at 37°C for 1 h, washed three times with PBS, and lysed with sterilized water for 15 min. Serial dilutions of the cell lysates were plated on BHIc agar and incubated anaerobically at 37°C for 48 h for enumeration of bacterial cells. For the invasion assay, SNU-C4 cells were incubated with bacterial cells for 4 h and washed three times with PBS. SNU-C4 cells were then incubated with fresh serum-free DMEM medium containing gentamicin (500 µg/mL) for 1 h, washed three times with PBS, and lysed with sterilized water for 15 min. Serial dilutions of the cell lysates were plated on BHIc agar and incubated anaerobically at 37°C for 48 h for enumeration of bacterial cells.

### Gene knockdown using siRNA

siRNA transfection was performed using Lipofectamin RNAiMAX transfection reagent, according to the manufacturer’s instructions (ThermoFisher Scientific). siRNAs used are listed in the KEY RESOURCES TABLE.

### Construction of AHR knockout in SNU-C4 and HCT116 CRC cell lines

AHR knockout (ΔAHR) in SNU-C4 and HCT116 CRC cell lines, respectively, was established by introducing indels targeting exon 2 of the AHR gene using the CRISPR/Cas9 method. Cas9- 2NLS ribonucleoprotein (10 µmol, Synthego) and guide RNA (100 pmol, Synthego) were electroporated into ∼1.2×10^5^ cells, using Neon Transfection System (ThermoFisher Scientific) at 1200 V for 20 msec and four pulses. After 48 h post-electroporation, the genomic DNA of electro-transfected cells was isolated, and the incorporation of indels was determined using a polyacrylamide gel electrophoresis-based (PAGE) genotyping (Zhu *et al*., 2014; PMID 25236476), with a pair of primers (AHR-PAGE-F/AHR-PAGE-R). Validated pools of cells were then subjected to clonal selection. The genomic DNA isolated from individual clones was analyzed by PCR with a pair of primers (AHR-KO-F/AHR-KO-R), and the PCR products of candidate clones were purified and sequenced by Sanger DNA sequencing (Genewiz) to analyze the edited sequences using online ICE (Inference of CRISPR Edits) Analysis Tool (Synthego). For the HCT116 CRC cell line, we obtained a homozygous HCT116^ΔAHR^ clone. For the SNU-C4 cell line, to clone edited sequences of each allele, PCR fragments generated from the genomic DNA of a heterozygous SNU-C4 ^ΔAHR^ clone were subjected to TA cloning, using the pGEM-T Easy Vector System (Promega) following the manufacturer’s instructions. Plasmids of *E. coli* DH5α transformants were individually isolated and submitted for Sanger sequencing (Genewiz). Primers used for the PAGE genotyping and for the PCR amplification of AHR alleles are listed in Table S7. The final AHR knockout clones were confirmed for the absence of AHR protein expression by western blot.

### Xenograft mouse experiments

Cells of wild-type SNU-C4, parent SNU-C4/gAHR, or SNU-C4^ΔAHR^ CRC cell line were grown to 50-80% confluency in DMEM medium supplemented with 10% FBS, washed with PBS, and harvested by trypsinization. Cell suspensions were then mixed with Matrigel at a 1:1 ratio and injected subcutaneously in the flank of athymic nude mice (5×10^5^ CRC cells per mouse). To prepare a bacterial suspension for intratumoral injection, the wild-type *Fn*, Δ*tnaA*, or Δ*tnaA*/p*tnaA* strain was grown overnight in BHIc at 37°C under anaerobic conditions. The next day, bacterial cells were harvested by centrifugation (6,000*g* × 15 min), washed with pre-reduced PBS (rPBS) twice, and resuspended in rPBS at a density of ∼5×10^8^ colony-forming units (CFU)/mL. After tumor formation, 100 µL of a bacterial suspension or the vehicle control (rPBS) was injected into tumors twice weekly. Tumor volumes were measured three times per week and calculated with the LWW formula (volume = 0.5 × length × width × width) (Tomayko and Reynolds, 1989; PMID 2544306).

### TRAP assay

The telomeric repeat amplification protocol (TRAP) assay was conducted with the TRAPeze Telomerase Detection Kit (Millipore Sigma) following the manufacturer’s instructions. The CRC cells were washed with PBS and lysed in cold CHAPS lysis buffer supplemented with RNase inhibitor (100 units/ml). Supernatant was collected by centrifugation at 12000 *g* for 20 min, and protein concentration was determined with the BCA protein assay kit (Pierce). Equal amounts of protein were examined for telomerase activity under the following thermal cycling conditions: 1 cycle at 30°C for 30 min, 1 cycle at 95°C for 2 min, followed by 30-32 cycles of 94°C for 15 sec, 59°C for 30 sec, 72°C for 1 min. Samples were run on a 10% PAGE in 0.5×Tris borate EDTA (TBE) buffer. The gel was then stained with SYBR green at a dilution of 1:1000 in 0.5×TBE for 30 min and visualized with the Azure imaging system (Azure Biosystems). The band intensity was determined using ImageJ (ver. 1.54). The values from TRAP products were normalized to those of the internal control.

### RNA sequencing

Respective CRC cell lines were seeded in 6-well plates (5×10^5^ cells/well) and cultured to 50- 70% confluency in DMEM medium with 10% FBS for 24 h. Media were aspirated, and cells were then incubated in serum-free DMEM containing FTIN (at 100 nM). After 4 h incubation, cells were harvested for RNA isolation using Trizol (Invitrogen) according to the manufacturer’s instructions. Isolated RNAs were treated with DNase I (Qiagen) and purified using the RNeasy MinElute Cleanup Kit (Qiagen). RNA integrity was assessed by agarose gel electrophoresis before sending it to Azenta for further quality check and processing for RNA sequencing (RNA-seq).

Library preparation, sequencing, data processing, and analysis were conducted at Azenta Life Sciences (South Plainfield, NJ, USA) as follows. RNA samples were quantified using Qubit 2.0 Fluorometer (ThermoFisher Scientific), and RNA integrity was checked using TapeStation (Agilent Technologies). RNA sequencing libraries were prepared using the NEBNext Ultra II RNA Library Prep for Illumina using the manufacturer’s instructions (New England Biolabs). Briefly, mRNAs were enriched with oligo-d(T) beads, fragmented for 15 min at 94°C, and used to synthesize first and second-strand cDNA. cDNA fragments were end- repaired and adenylated at 3′ends, and universal adapters were ligated to cDNA fragments, followed by index addition and library enrichment by PCR with a limited number of cycles. The sequencing libraries were validated on the Agilent TapeStation (Agilent Technologies) and quantified by using Qubit 2.0 Fluorometer (ThermoFisher Scientific) as well as by quantitative PCR (KAPA Biosystems). The sequencing libraries were then multiplexed and clustered onto a flowcell on the Illumina NovaSeq X Plus instrument according to the manufacturer’s instructions. The samples were sequenced using a 2×150 bp paired-end configuration. Image analysis and base calling were conducted by the NovaSeq Control Software (NCS). Raw sequence data (.bcl files) generated from Illumina NovaSeq X Plus were converted into fastq files and de-multiplexed using Illumina bcl2fastq 2.20 software. One mismatch was allowed for index sequence identification.

Sequence reads were trimmed to remove possible adapter sequences and nucleotides with poor quality using Trimmomatic v.0.36. The trimmed reads were mapped to the Homo sapiens GRCh38 reference genome available on ENSEMBL using the STAR aligner v.2.5.2b. The STAR aligner is a splice aligner that detects splice junctions and incorporates them to help align the entire read sequences. BAM files were generated as a result of this step. Unique gene hit counts were calculated using featureCounts from the Subread package v.1.5.2. Unique reads (with no mismatch) that fell within exon regions were counted. After the extraction of gene hit counts, the gene hit counts table was used for downstream differential expression analysis. Using DESeq2, a comparison of gene expression between FTIN-treated and vehicle (DMSO)- treated groups was performed. The Wald test was used to generate *p*-values and log_2_ fold changes. Genes with an adjusted *p*-value < 0.05 and absolute log_2_ fold change >0.5 were called as differentially expressed for each comparison. RNA-seq data were deposited to NCBI with accession number (BioProject ID: PRJNAxxxxxxx)

### 16S rRNA gene amplicon sequencing

Genomic DNA was isolated from ∼10 mg of the respective human CRC and normal colon tissues (a total of 60 samples) using ZymoBIOMICS^TM^ DNA Miniprep Kit (Zymo Research) and sent to Zymo Research for bacterial 16S rRNA gene targeted sequencing service. An absolute abundance of bacterial genomes in respective DNA samples was determined by qRT- PCR. The standard curve was made with plasmid DNA containing one copy of the 16S rRNA gene prepared in 10-fold serial dilutions. The primers used were the same as those used in Bacterial 16S Targeted Library Preparation. The equation generated by the plasmid DNA standard curve was used to calculate the number of 16S rRNA gene copies in the reaction for each sample. The number of genome copies per microliter DNA sample (genome_copies) was calculated by dividing the obtained 16S rRNA gene copy number by an assumed number (4) of 16S rRNA gene copies per genome. Bacterial 16S rRNA gene targeted sequencing was performed using the Quick-16S™ NGS Library Prep Kit (Zymo Research). The sequencing library was prepared by PCR with pre-designed primers amplifying the V3-V4 region of 16S rRNA. The final PCR products were quantified with qPCR, cleaned with the Select-a-Size DNA Clean & Concentrator^™^ (Zymo Research), then quantified with TapeStation^®^ (Agilent Technologies) and Qubit^®^ (Thermo Fisher Scientific). The final library was sequenced on Illumina^®^ Nextseq^™^ with a P1 reagent kit. Unique amplicon sequence variants were inferred from raw reads using the DADA2 pipeline. Potential sequencing errors and chimeric sequences were also removed with the DADA2 pipeline. Taxonomy assignment was performed using Uclust from Qiime v.1.9.1 with the Zymo Research Database, a 16S database that is internally designed and curated, as a reference. Raw data are deposited to NCBI with accession number (BioProject ID: PRJNAxxxxxxx)

### FTIN measurement in human CRC and normal colon tissues

Two pieces (each ∼10 mg) of cuts from 60 respective human CRC and normal colon tissue samples were homogenized in 1 ml of 50% methanol (v/v in water) containing the internal standard (FTIN-^13^C_6_, 0.5 ng/mL) using TissueLyser II (Qiagen) with 5 mm stainless steel beads at a frequency of 30/s for 30 sec. Homogenization was repeated twice. After keeping homogenates on ice for 20 min, tissue lysates were clarified by centrifugation (16,000*g* at 4°C × 10 min) for the collection of supernatant. The supernatant was then dried *in vacuo* using a Vacufuge (Eppendorf), reconstituted with 100 µl of 50% acetonitrile containing 0.1% formic acid, and subjected to LC-MS/MS analysis as described for FTIN measurement above.

### Statistical analysis

Statistical analysis was performed using GraphPad Prism (ver. 10.6.0; GraphPad Inc., La Jolla, CA, USA). A *p*-value less than 0.05 was considered significant.

## KEY RESOURCES TABLE

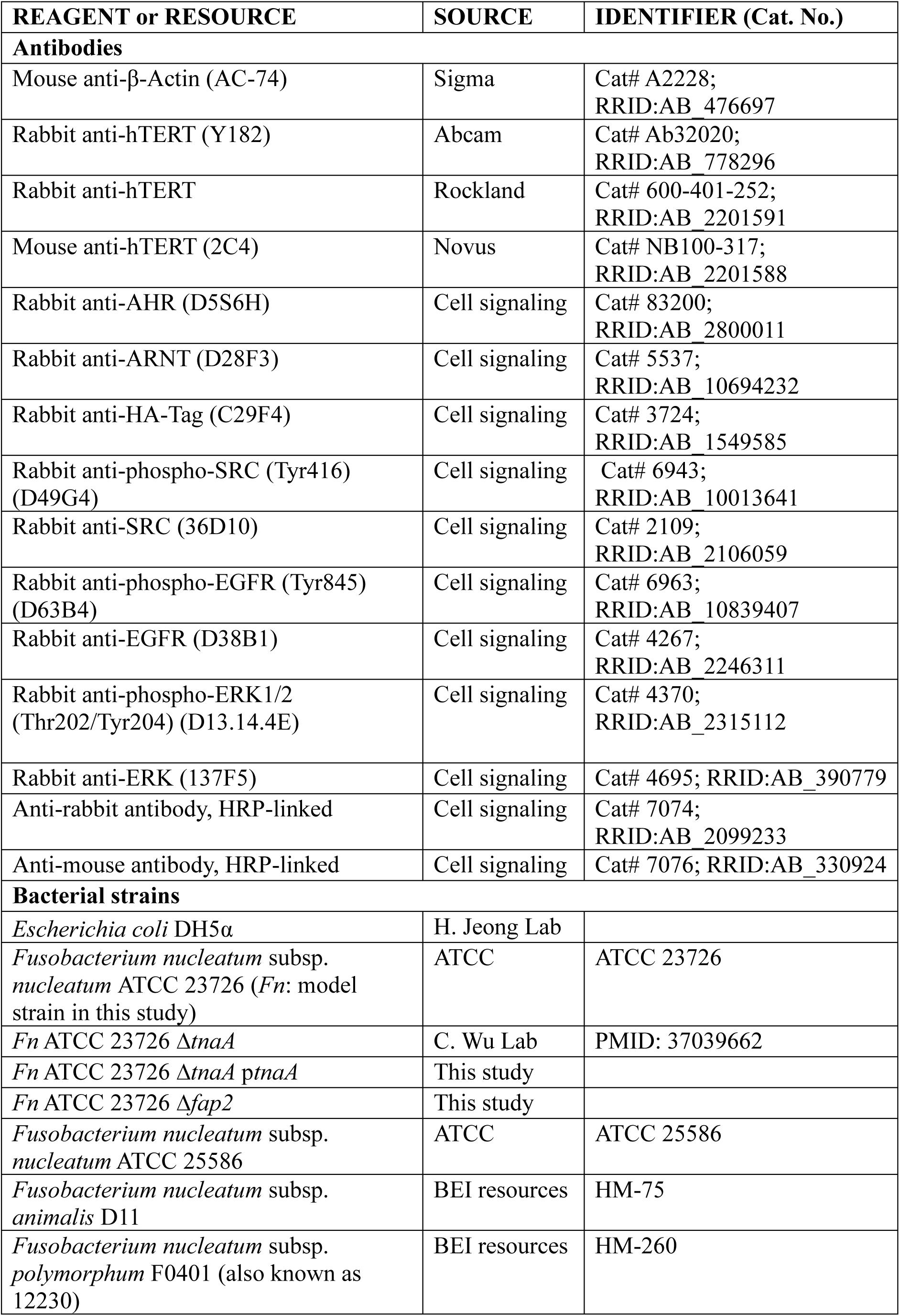
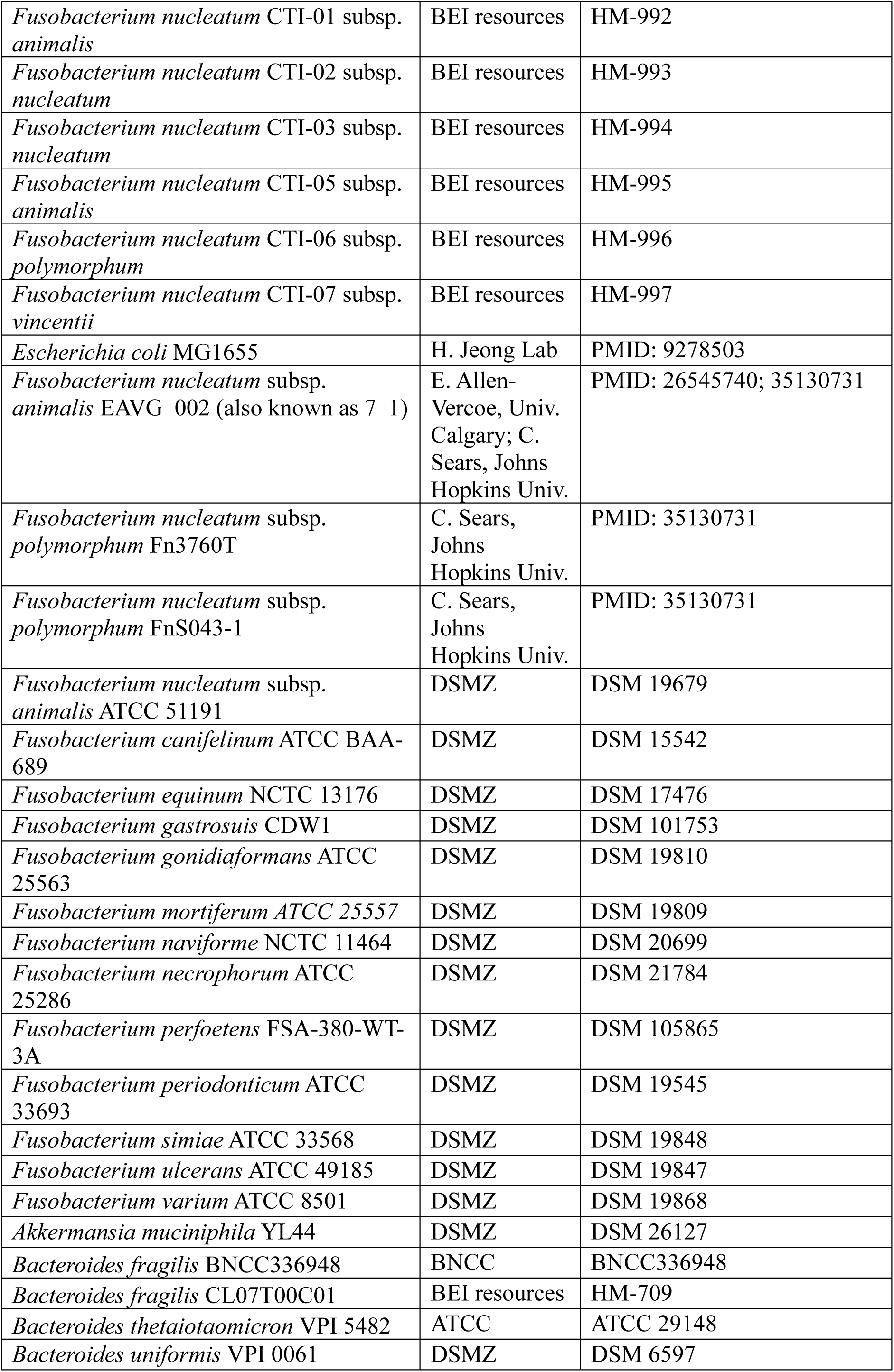
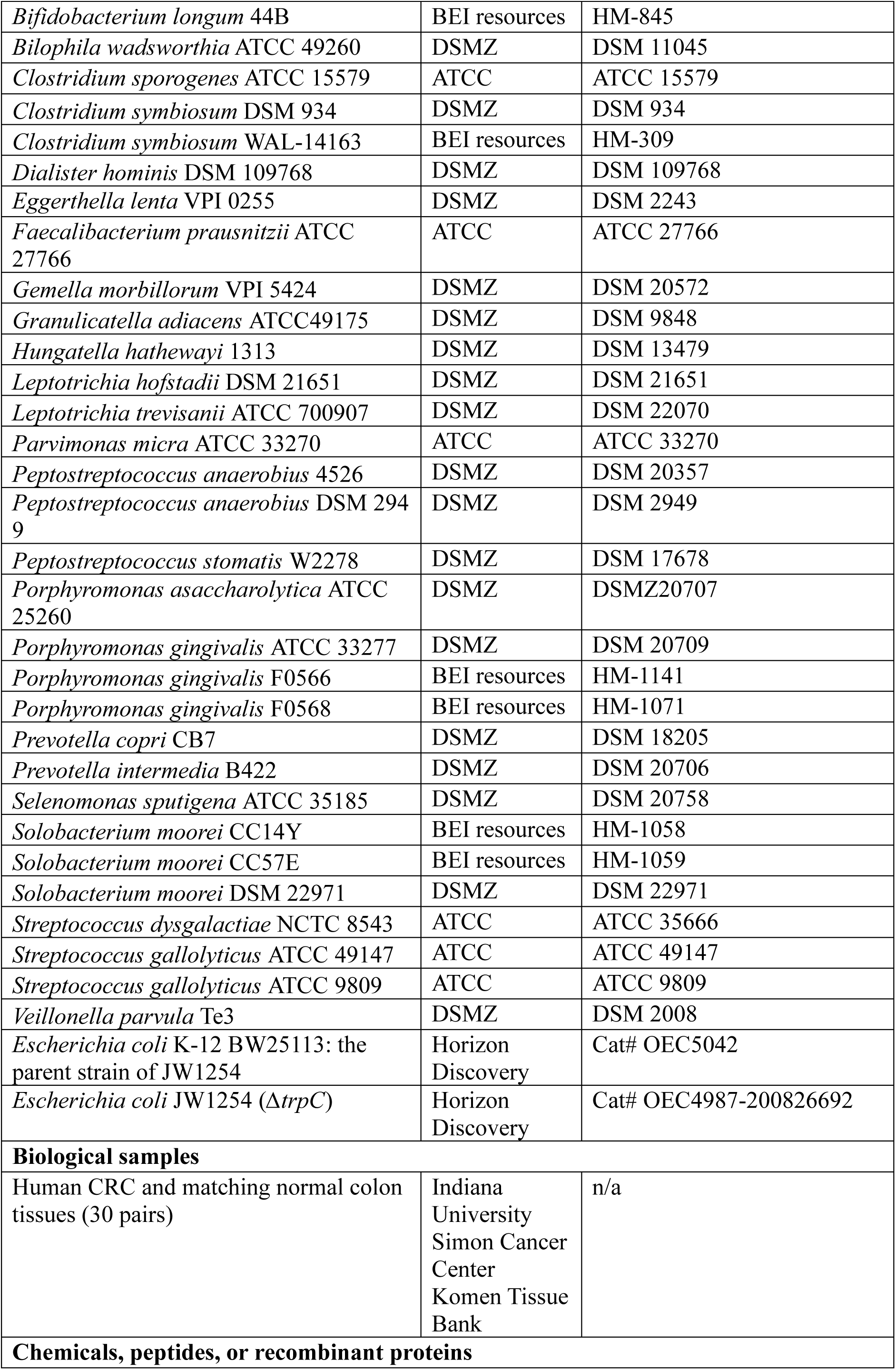
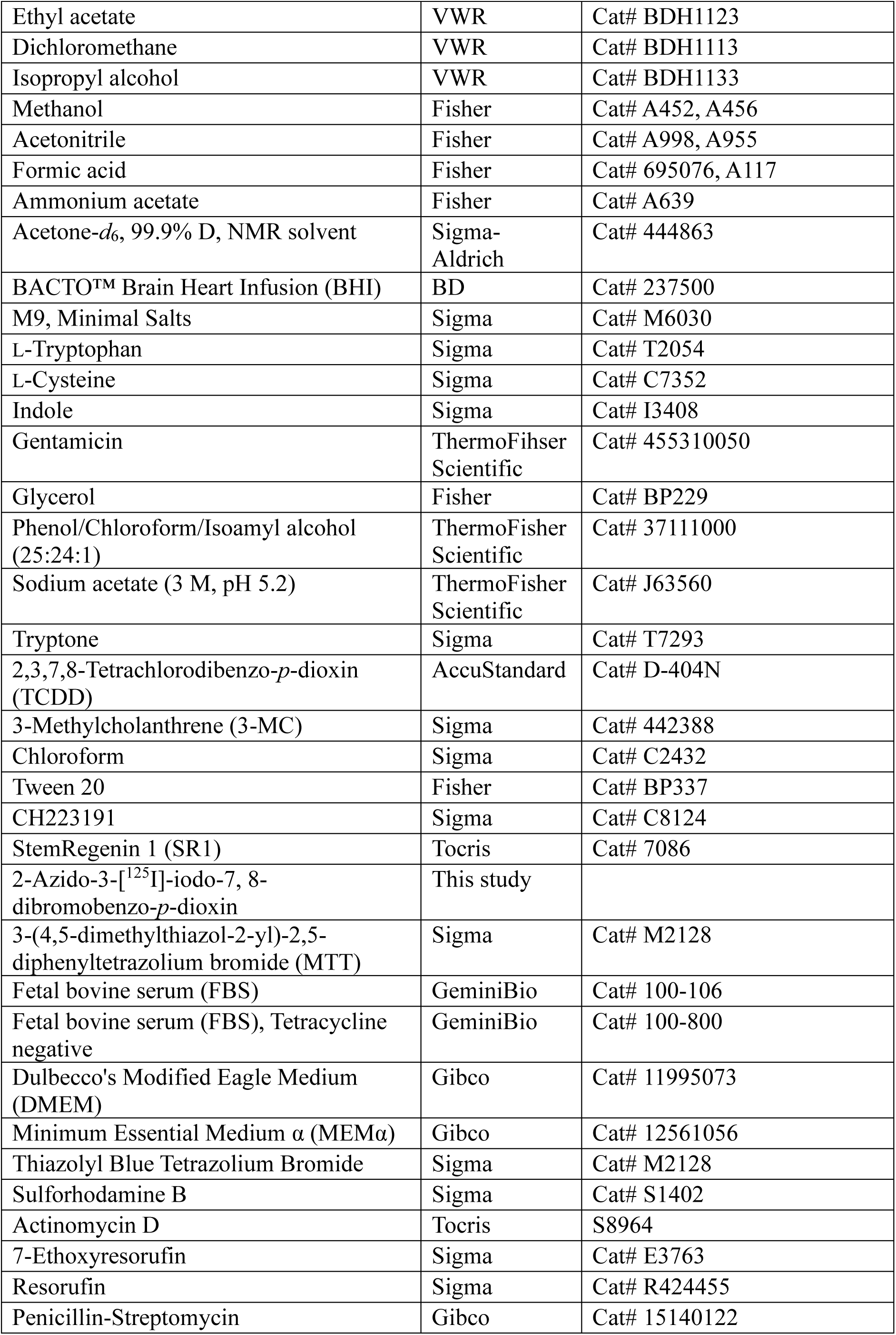
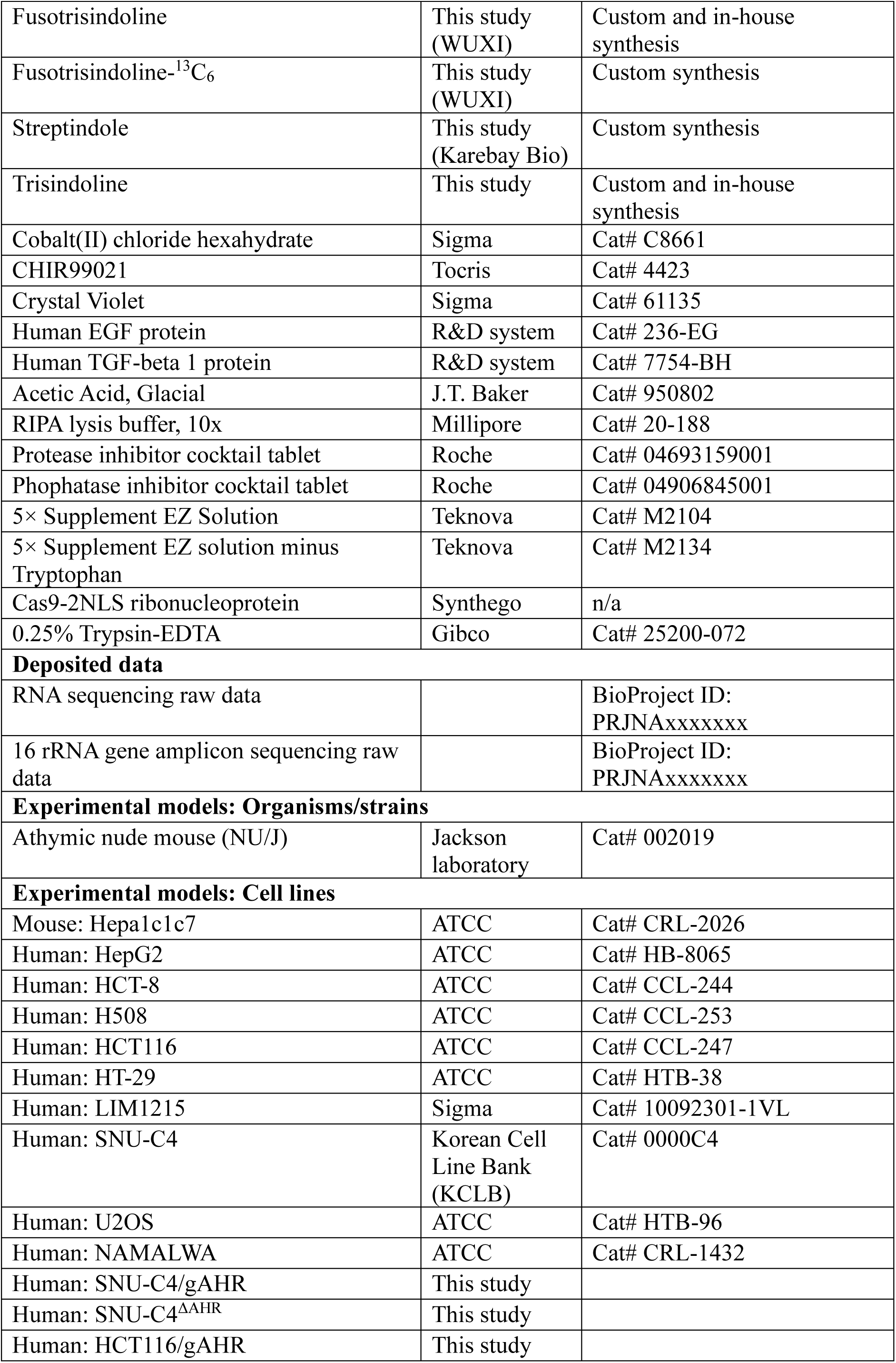
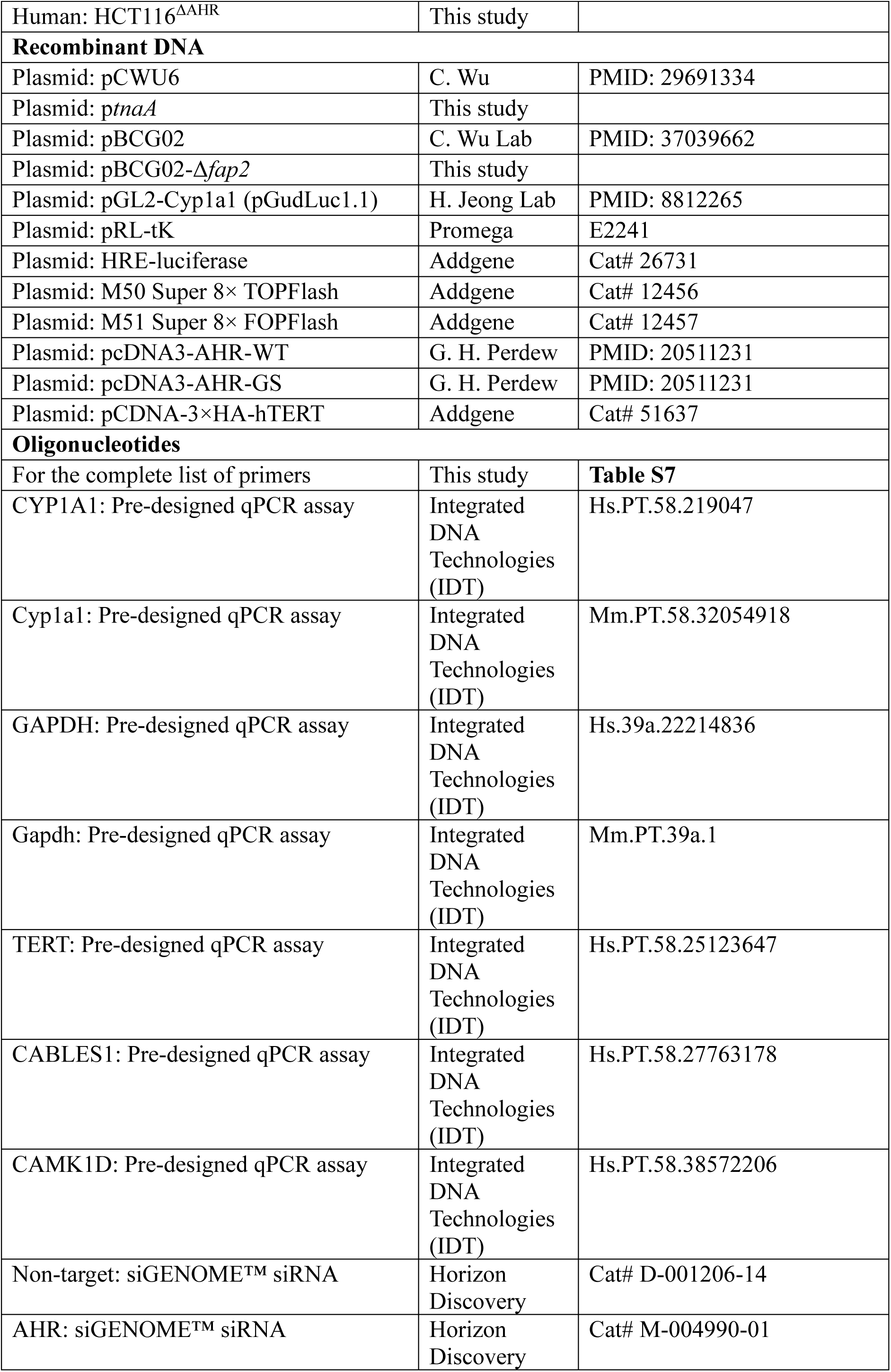
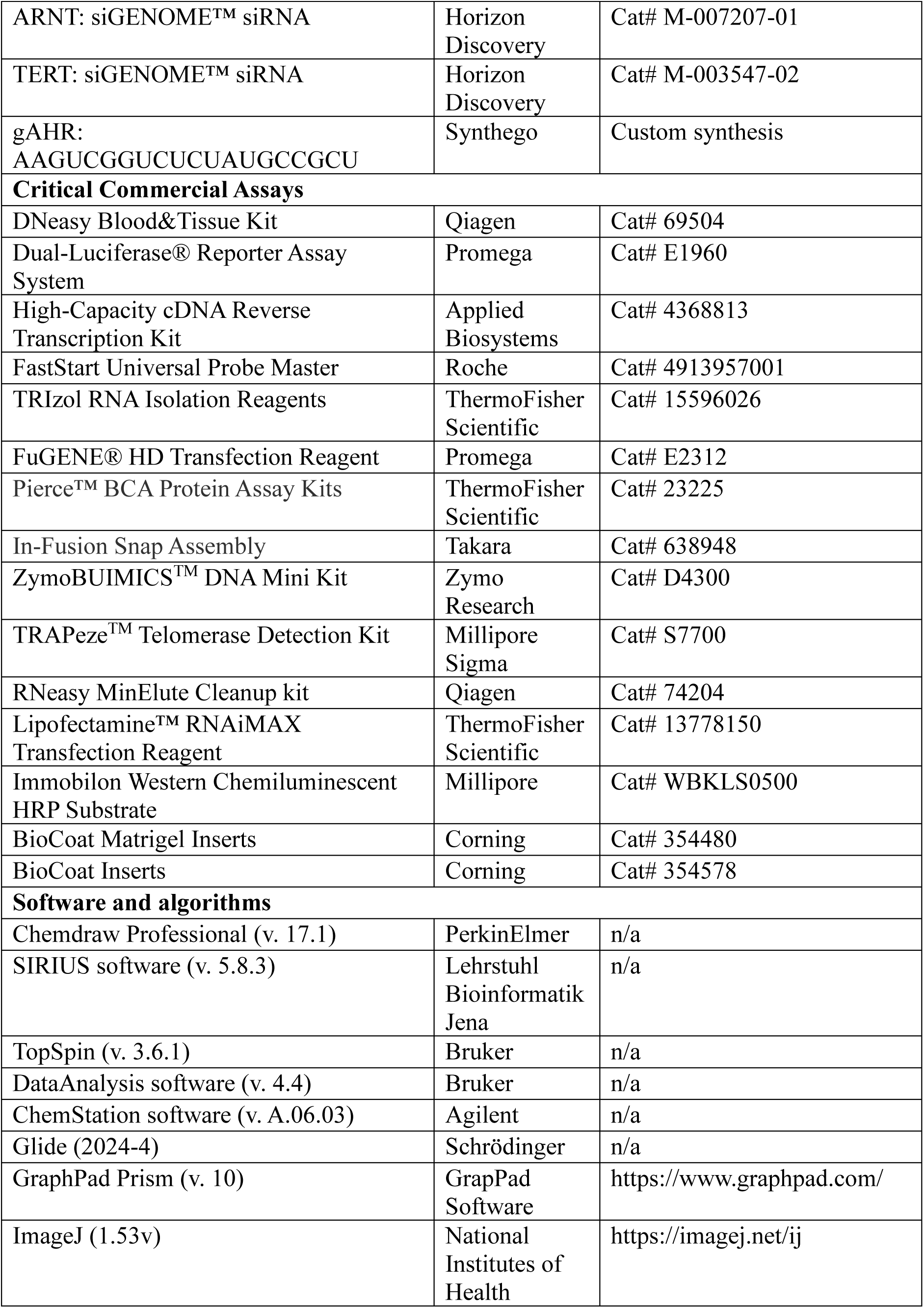

## Supplemental Information

***Fusobacterium nucleatum* produces previously unappreciated AHR-activating metabolites and promotes CRC cell proliferation via the AHR-TERT axis**

Kyoung-Jae Won, Pei-Ru Jin, Lydia Davis, Jimmy Orjala, Abigail T. Armstrong, Wen-Hung Wang, Harish Kothandaraman, Sagar M. Utturkar, Dawon Bae, Byeong-Seon Jeong, Chenggang Wu, Iain A. Murray, Gary H. Perdew, Chijian Xiang, Jianing Li, Hyunwoo Lee, Hyunyoung Jeong

**Figure S1.**
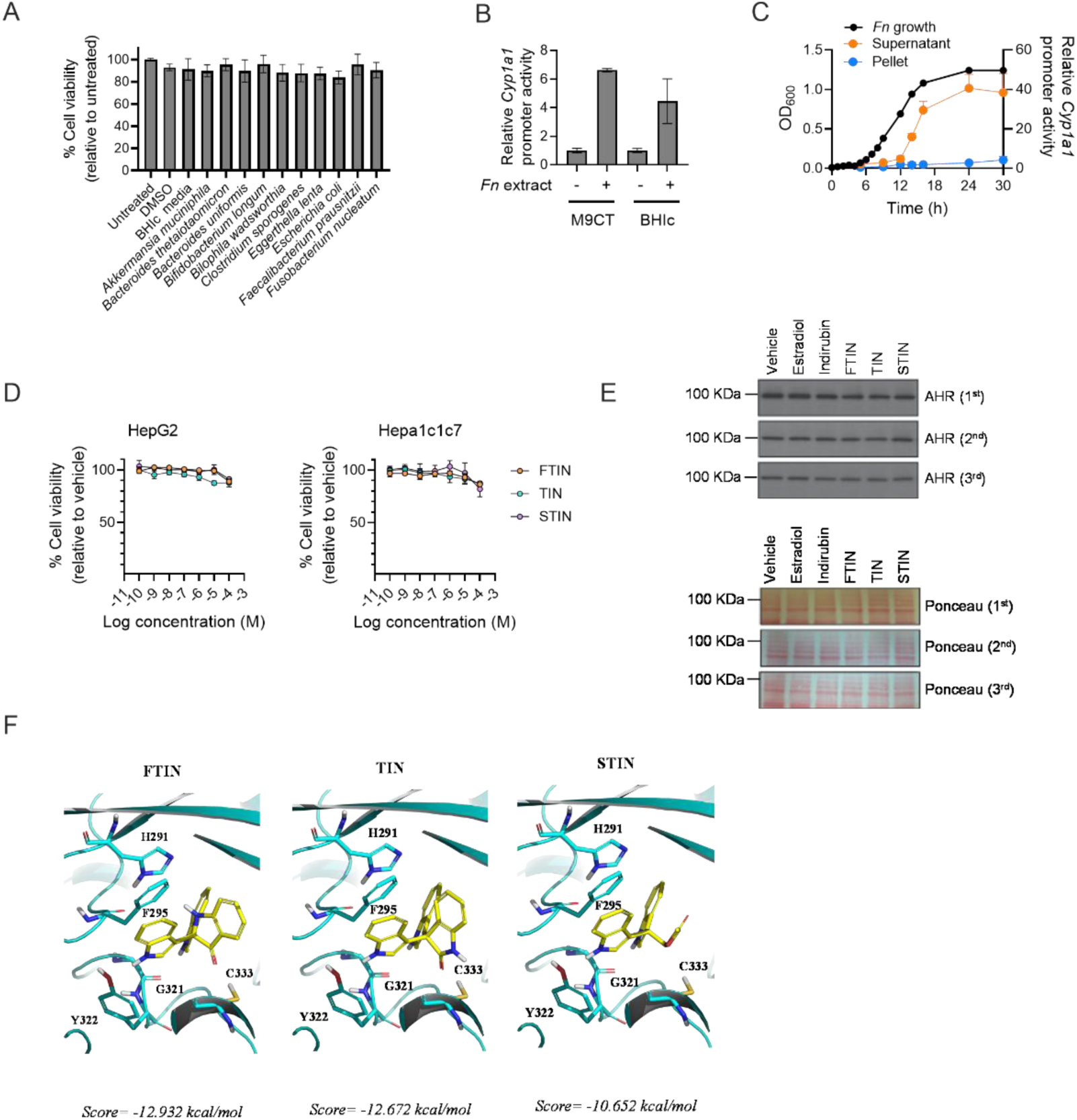
Characterization of three AHR-activating *Fn* metabolites. Related to Figure 1. (A) Respective bacterial extracts are not cytotoxic against HepG2 cells under conditions for the AHR reporter assays (related to Figure 1A). HepG2 cells were treated with respective bacterial extracts for 24 h, and cell viability was determined using the MTT assay (*n* = 3). (B) Organic extracts of *Fn* cultures grown in either BHIc or M9CT media activated AHR to a similar extent (*n* = 3) (related to Figure 1B and Supplemental Texts for the isolation and structure determination of *Fn* metabolites). (C) *Fn*-mediated AHR-inducing activity is present mainly in *Fn* culture supernatants, rather than cell pellets (related to Figure 1B and Supplemental Texts for the isolation and structure determination of *Fn* metabolites). HepG2 cells transfected with the AHR reporter system were treated with organic extracts of *Fn* cell pellet or M9CT culture supernatants prepared from the indicated time points (*n* = 3). *Fn* growth in M9CT was determined by measuring the optical density of the culture at 600 nm (OD_600_). (D) *Fn* metabolites (FTIN, STIN, and TIN) up to 1 µM were not cytotoxic against HepG2 or Hepa1c1c7 cells under conditions for the AHR reporter assay (related to Figures E and F). Respective cell lines transfected with the AHR reporter system were treated with each metabolite at varying concentrations for 4 h, and cell viability was determined using the SRB assay (*n* = 3). (E) Related to Figure 1H. Treatment of *Fn* metabolites and control compounds does not affect AHR expression and total protein levels in HN-30 cells (*n*=3). The expression of the AHR protein was assessed by western blotting, and total protein amounts loaded on the SDS- PAGE were visualized by Ponceau staining. (F) *in silico* models of the *Fn* metabolites (FTIN, STIN, and TIN) docked into the ligand- binding PAS domain of the AHR in the cryo-EM structure of the ternary AHR complex (Hsp90-XAP2-AHR) (PDB ID: 7ZUB). Data are represented as mean ± SD, where appropriate.

**Figure S2.**
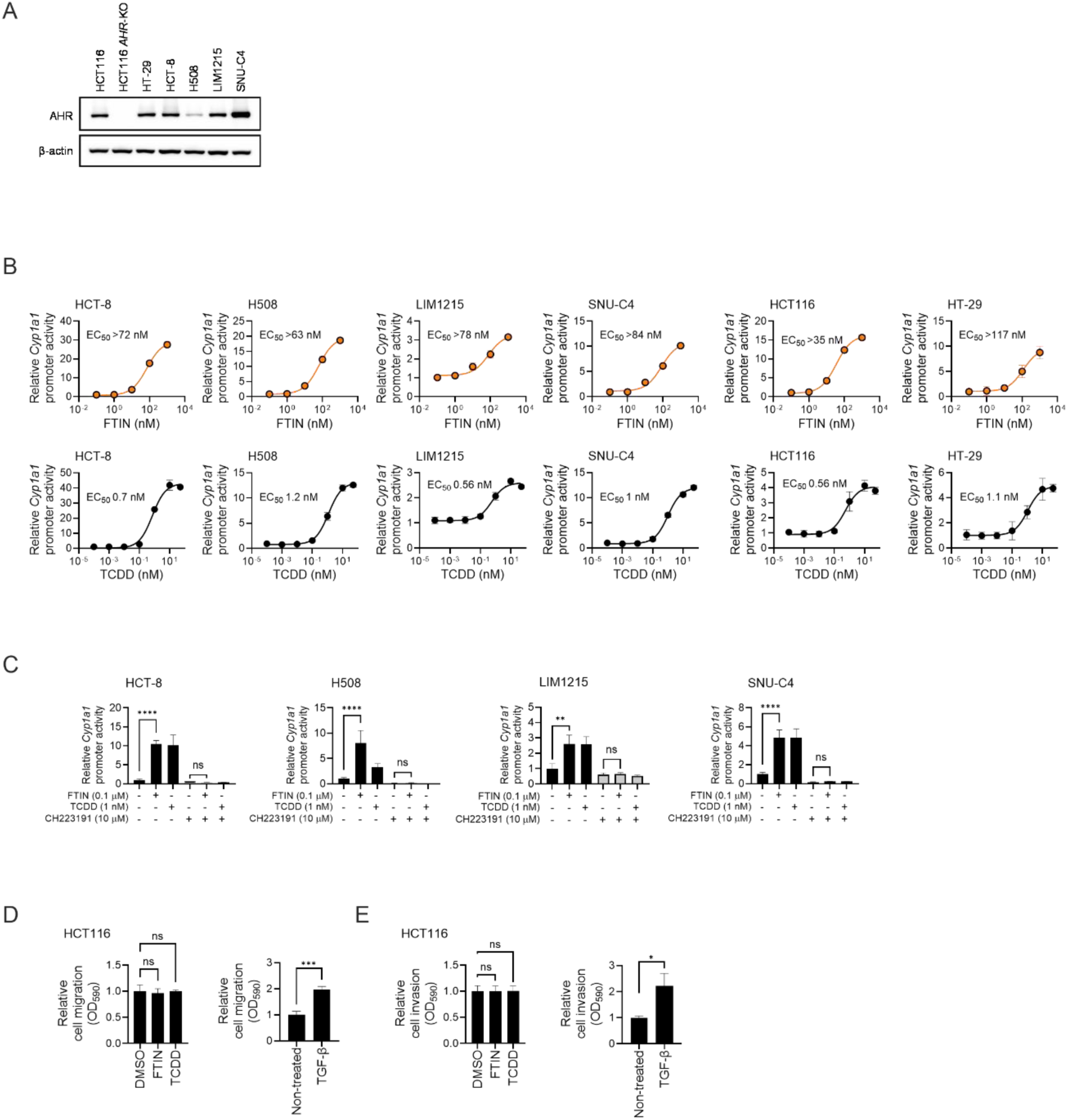
FTIN activates AHR in six different CRC cell lines. Related to Figure 2. (A) The expression of the AHR protein in six CRC cell lines used in Figure 2A was assessed by western blot, with β-actin as a loading control. (B) The induction of *Cyp1a1* promoter activity by FTIN or TCDD (positive control) in six different CRC cell lines (*n* = 3). Respective cell lines transfected with the AHR reporter system were treated with FTIN or TCDD at varying concentrations for 4 h. The AHR reporter assay was performed as described in Experimental Method Details. (C) The AHR antagonist CH223191 blocks the induction of *Cyp1a1* promoter activity by FTIN (*n* = 3). Respective cell lines were pretreated with CH223191 (10 µM), followed by treatment with FTIN (0.1 µM), TCDD (1 nM), or vehicle (DMSO) for 4 h. The AHR reporter assay was performed as described in Experimental Method Details. (D, E) Cell migration (D) and invasion (E) were determined using the Transwell assay (*n* = 3). Cells were pretreated with CH223191 (10 µM) for 1 h, followed by FTIN, TCDD, or vehicle (DMSO) treatment for 24 h. TGF-β was used as a positive control. Data are represented as mean ± SD, where appropriate. Statistical significance was assessed by ordinary one-way ANOVA with Tukey’s multiple comparisons test (C-D): *, *p* < 0.05; **, *p* < 0.01; ***, *p* < 0.001; ****, *p* < 0.0001; ns, not significant.

**Figure S3.**
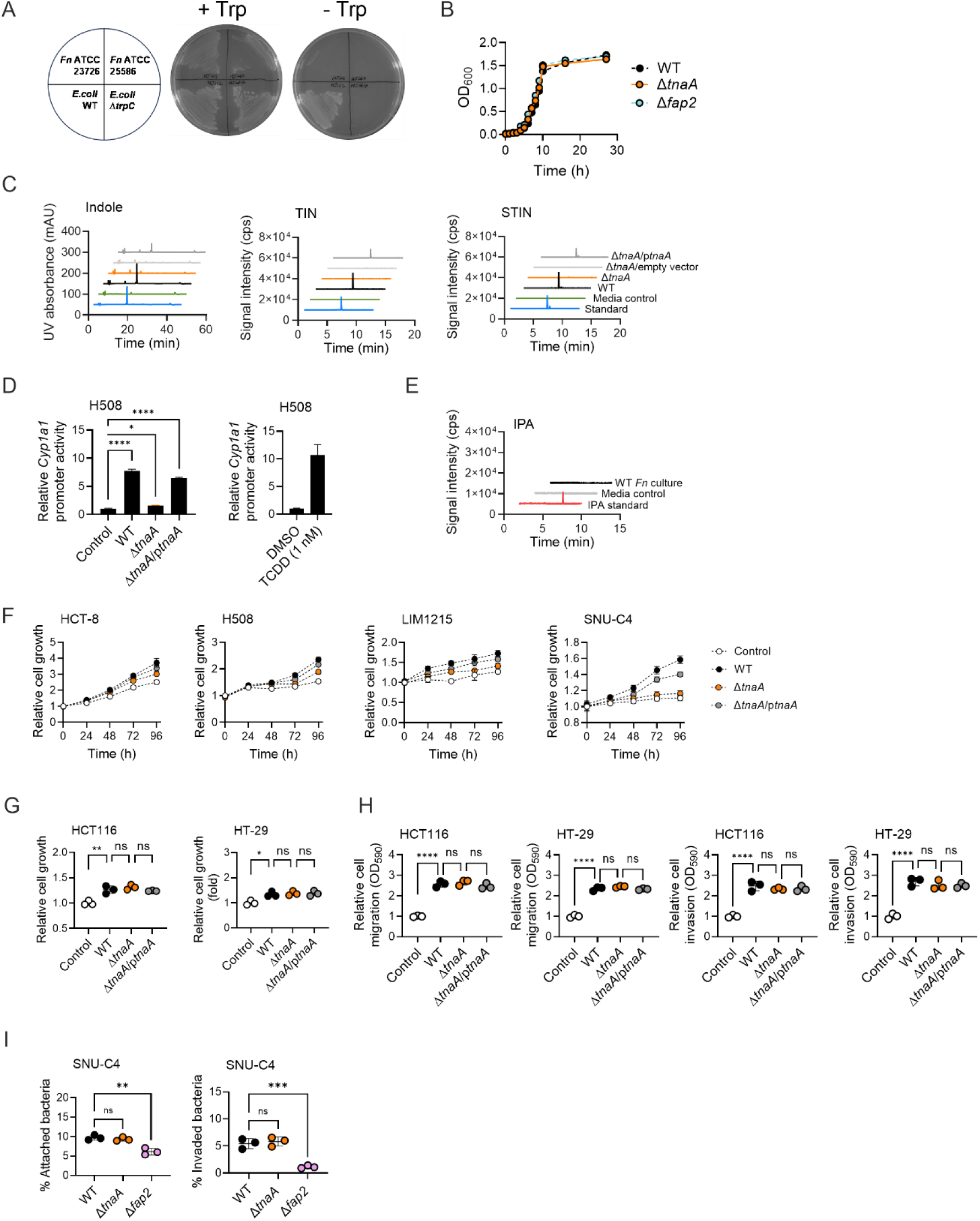
A Δ*tnaA* mutant defective in FTIN production induces distinct phenotypes between SNU-C4 and HCT116 CRC cell lines. Related to Figure 3. (A) Confirmation of L-tryptophan auxotrophy of *Fn* ATCC 23726 and *Fn* ATCC 25586 strains. Parent *E. coli* BW25113 and its isogenic Δ*trpC* strains were used as controls for L-tryptophan prototrophy and auxotrophy, respectively. Respective strains were streaked on M9-based agar medium with or without L-tryptophan. (B) Both Δ*tnaA* and Δ*fap2* mutants exhibit wild-type growth in BHIc medium. Shown is from a single experiment. OD_600_: optical density at 600 nm. (C) Related to Figure 3C, lack of TIN, STIN, and indole production in the culture of the Δ*tnaA* strain grown in BHIc. Shown is a representative of an HPLC-UV chromatogram of indole and LC/MS/MS spectra of TIN and STIN from two independent experiments with similar results. (D) The Δ*tnaA* strain is unable to induce *Cyp1a1* promoter activity in the H508 CRC cell line. CRC cells transfected with the AHR reporter system were co-incubated with respective *Fn* strains at an MOI of 300 for 4 h. AHR reporter assay was performed as described in Experimental Method Details. TCDD was used as a positive control for AHR activation. (E) LC/MS/MS spectra of 3-indolepropionic acid (IPA) measured in culture supernatants of wild-type *Fn* grown anaerobically in M9CT media for 24 h. Shown is a representative of two independent experiments with similar results. (F) The Δ*tnaA* strain was unable to promote the proliferation of four different CRC cell lines (HCT8, H508, LIM1215, and SNU-C4). Respective CRC cell lines were co-incubated with an *Fn* strain (wild-type, Δ*tnaA*, or Δ*tnaA*/p*tnaA* at an MOI of 300), serum-free media containing *Fn* were renewed every 24 h, and CRC cell growth was determined over time using the SRB assay. (G) The Δ*tnaA* strain retained wild-type growth-promoting activity for HCT116 and HT-29 CRC cell lines. Respective CRC cell lines were co-incubated with an *Fn* strain (MOI 300), serum-free media containing *Fn* were renewed every 24 h, and CRC cell growth was determined at 72 h using the SRB assay. (H) The Δ*tnaA* strain retained wild-type promoting activity for migration (left two) and invasion (right two) of HCT116 and HT-29 CRC cell lines. Cell migration and invasion were determined by the Transwell assay. HCT116 or HT-29 CRC cell line was co-incubated with respective bacterial strains for 24 h, and migration and invasion were determined as described in Experimental Method Details. (I) The Δ*tnaA* strain exhibits wild-type levels of attachment to and invasion into SNU-C4 cells (*n* = 3). Data are represented as mean ± SD, where appropriate. Statistical significance was assessed by ordinary one-way ANOVA with Dunnett’s multiple comparisons test (D, F-I): *, *p* < 0.05; **, *p* < 0.01; ***, *p* < 0.001; ****, *p* < 0.0001; ns, not significant.

**Figure S4.**
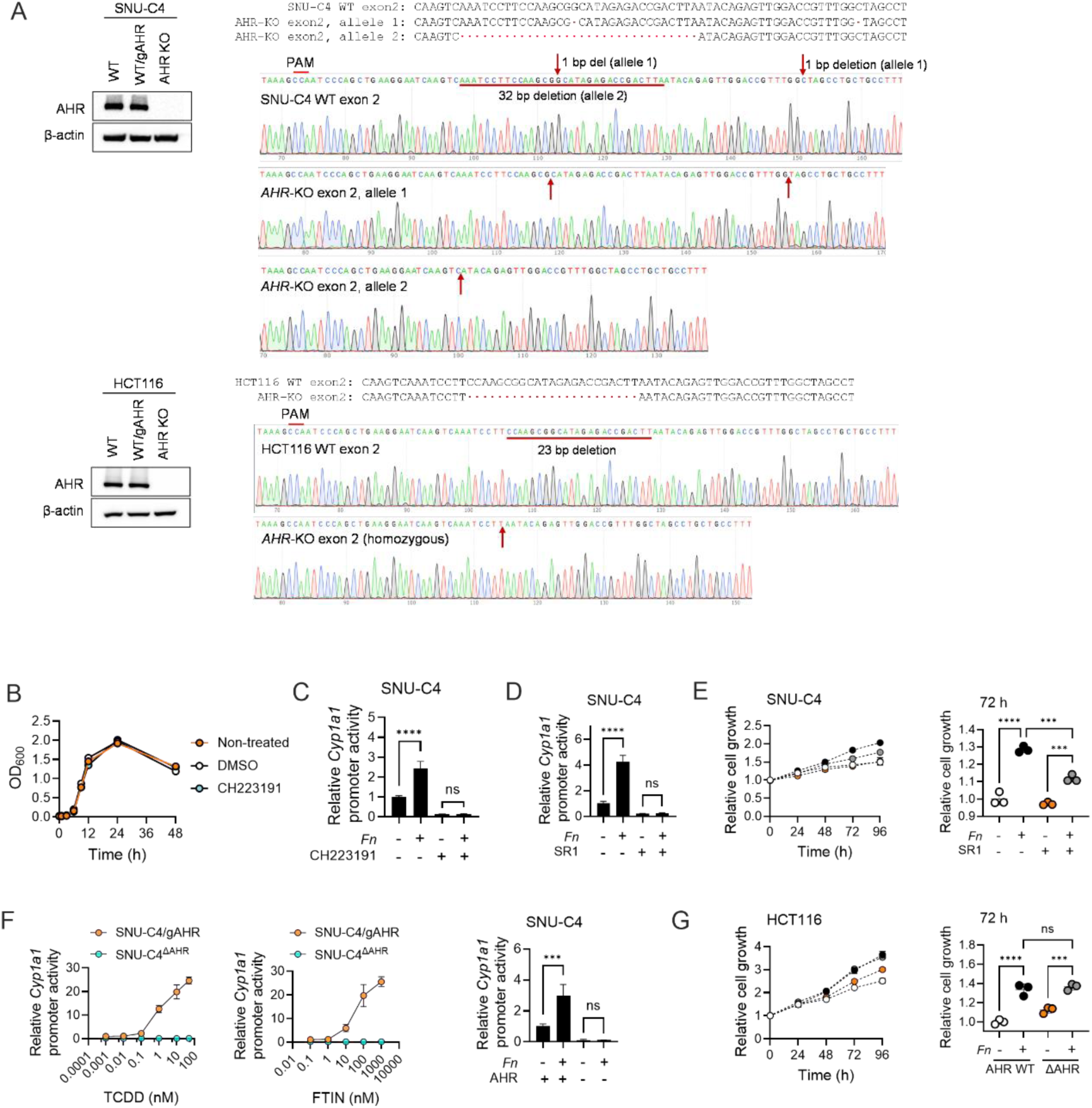
The effect of AHR inhibition or knockout on *Fn*-mediated AHR activation in CRC cells and on *Fn*-promoted CRC cell proliferation. Related to Figure 4. (A) Confirmation of AHR knockout by Sanger DNA sequencing and no AHR protein expression in SNU-C4^ΔAHR^ and HCT116^ΔAHR^ cells by western blotting (B) No growth-inhibitory effect of the AHR antagonist CH223191 (10 µM) or vehicle (DMSO) on *Fn* ATCC 23726 grown in BHIc. Shown is a representative of two independent experiments. (C and D) Both AHR antagonists, CH223191 (C) and SR1 (D), inhibit *Fn*-mediated induction of *Cyp1a1* promoter activity in SNU-C4 cells (*n* = 3). SNU-C4 cells transfected with the AHR reporter system were treated with CH223191 (10 µM), SR1 (1 µM), or DMSO in serum-free medium for 1 h and subsequently co-incubated with *Fn* (at MOI 300) for 4 h. The AHR reporter assay was performed as described in Experimental Method Details. (E) The AHR antagonist SR1 inhibits *Fn*-mediated promotion of SNU-C4 cell proliferation (*n* =3). SNU-C4 CRC cells were treated with SR1 (1 µM) for 1 h, and subsequently co-incubated with *Fn* (at MOI 300). Fresh serum-free medium containing Fn was renewed every 24 h, and CRC cell growth was determined using the SRB assay every 24 h up to 96 h. (F) Neither FTIN nor *Fn* induces *Cyp1a1* promoter activity in SNU-C4^ΔAHR^ cells (*n* = 3). TCDD was used as a positive control for the induction of *Cyp1a1* promoter activity. Respective CRC cell lines (the parent SNU-C4/gAHR and SNU-C4^ΔAHR^) transfected with the AHR reporter system were treated with TCDD or FTIN at varying concentrations, or co-incubated with *Fn* (at MOI 300) for 4 h. The AHR reporter assay was performed as described in Experimental Method Details. (G) *Fn* promotes the proliferation of the HCT116^ΔAHR^ cell line, as well as the parent HCT116/gAHR (*n* = 3). Respective CRC cell lines (parent HCT116/gAHR and HCT116^ΔAHR^) were co-incubated with *Fn* (at MOI 300) for up to 96 h. Fresh serum-free medium was renewed every 24 h, and cell growth was determined using the SRB assay every 24 h. Data are represented as mean ± SD, where appropriate. Statistical significance was assessed by ordinary one-way ANOVA with Dunnett’s multiple comparisons test: ***, *p* < 0.001; ****, *p* < 0.0001; ns, not significant.

**Figure S5.**
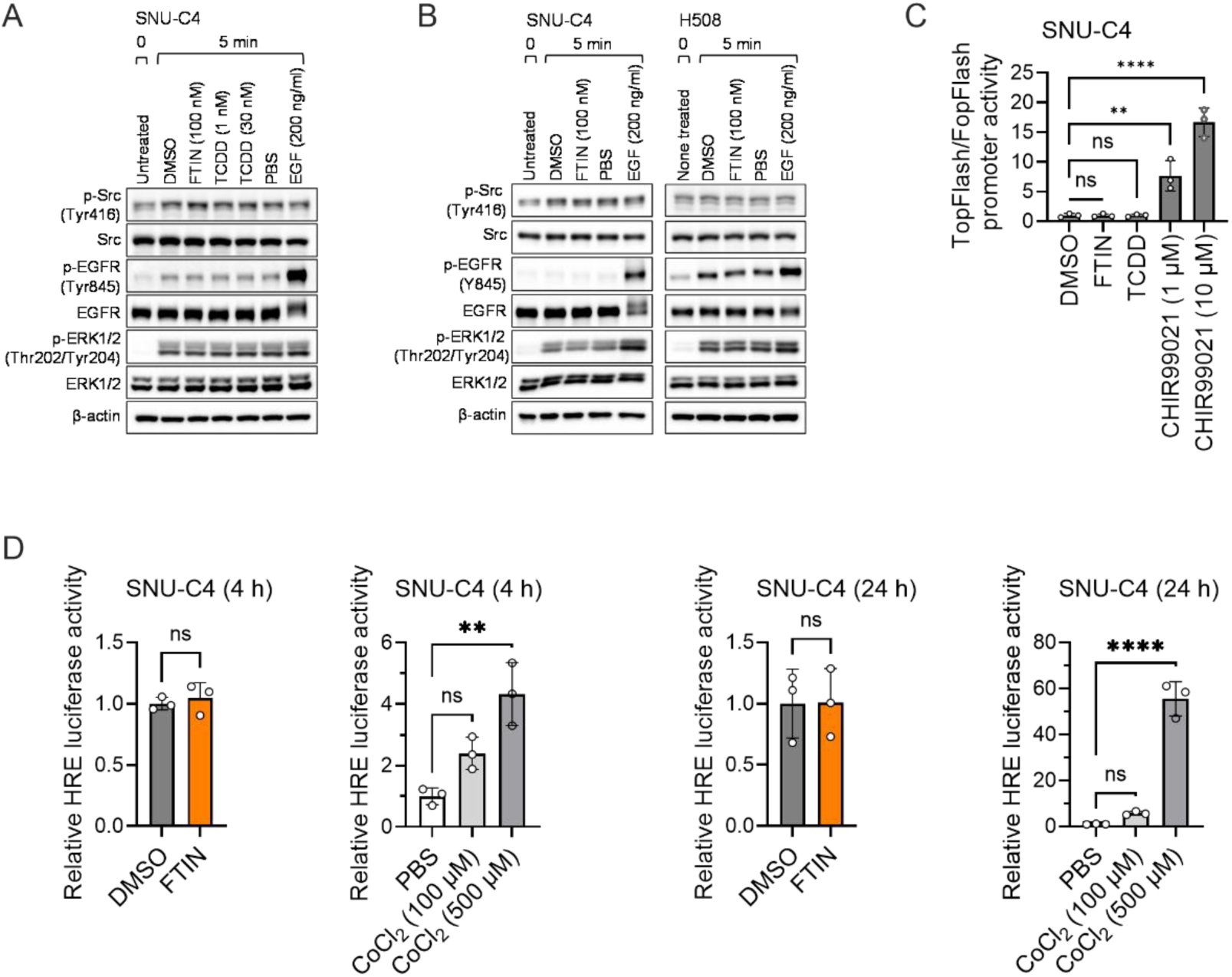
FTIN does not activate the Src kinase, β-catenin, or HIF-1 pathway. Related to Figure 5. (A) Neither TCDD nor FTIN activates Src kinase in the SNU-C4 cell line. (B) FTIN does not activate the Src kinase, neither in SNU-C4 CRC nor in H508 CRC cell lines. Respective CRC cell lines were serum-starved for 24 h and treated with FTIN (100 nM), EGF (200 ng/ml), or vehicle (DMSO or PBS) and harvested at the indicated time point. Protein and phosphorylation levels of Src, EGFR, and ERK1/2 were determined by western blot. β-actin was used as a loading control. EGF was used as a positive control for EGFR and/or ERK1/2 activation. Shown is a representative of more than three independent experiments. (C) Neither FTIN nor TCDD activates β-catenin (*n* = 3). SNU-C4 cells were transfected with a β-catenin reporter system for 24 h and treated with FTIN (0.1 µM), TCDD (1 nM), or vehicle (DMSO) in serum-free medium for 24 h. The GSK3 inhibitor CHIR99021 (at 1 µM and 10 µM) was used as a positive control for β-catenin activation. The β-catenin reporter assay was performed as described in Experimental Method Details. (D) FTIN does not activate the HIF-1 transcription factor (*n* = 3). SNU-C4 CRC cells transfected with a luciferase-based HIF-1 reporter system were treated with FTIN (100 nM) or vehicle (DMSO) in serum-free medium for 4 h or 24 h. Cobalt chloride (in PBS at 100 µM and 500 µM) was used as a positive control for HIF-1 activation. The HIF-1 reporter assay was performed as described in Experimental Method Details. Data are represented as mean ± SD, where appropriate. Statistical significance was assessed by ordinary one-way ANOVA with Tukey’s multiple comparisons test (C), Welch’s *t* test (DMSO vs. FTIN in the HIF-1 reporter assay), and ordinary one-way ANOVA with Tukey’s multiple comparisons test (PBS vs. CoCl_2_): **, *p* < 0.01; ****, *p* < 0.0001; ns, not significant.

**Figure S6.**
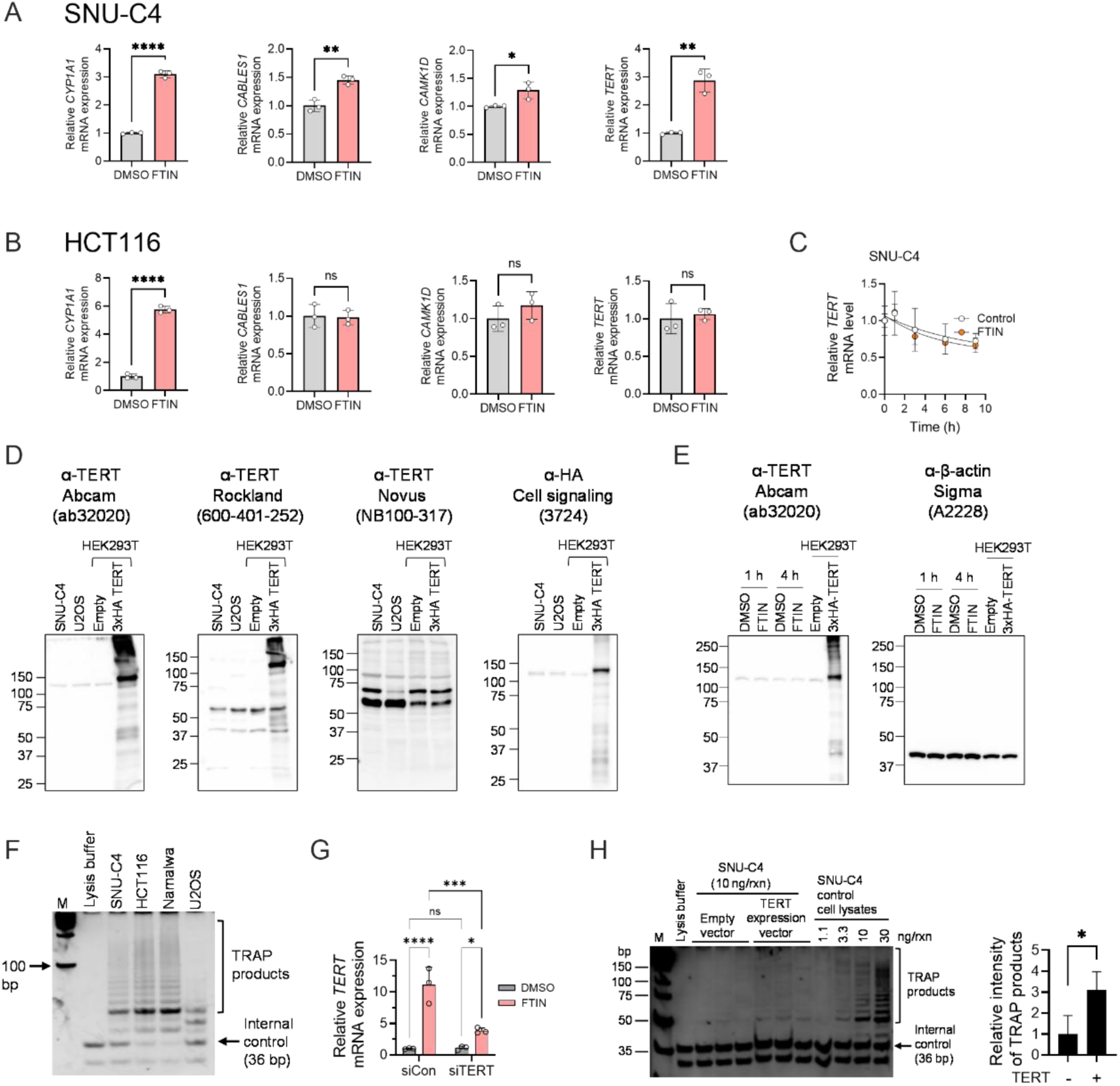
Additional information on FTIN-induction of TERT, anti-TERT antibodies, telomerase enzyme assays, and TERT overexpression in SNU-C4 cells. Related to Figure 6. (A) qRT-PCR validation of four FTIN-responsive genes (*CYP1A1*, *CABLES1*, *CAMK1D*, and *TERT*) identified in SNU-C4 CRC cells by RNA-seq (*n* = 3). (B) qRT-PCR validation of FTIN-responsive *CYP1A1* and three FTIN-nonresponsive genes (*CABLES1*, *CAMK1D*, and *TERT*) identified in HCT116 CRC cells by RNA-seq (*n* = 3). (C) No effect of FTIN on the half-life of *TERT* mRNA (*n* = 3). SNU-C4 CRC cells treated with FTIN (100 nM) or vehicle (DMSO) were treated with actinomycin D (5 µg/ml) or vehicle (DMSO) in serum-free medium for 9 h, and *TERT* mRNA levels were determined by qRT-PCR. (D) Three different commercial anti-TERT antibodies fail to detect endogenous TERT protein. Respective anti-TERT antibodies were tested to detect the TERT protein in cell lysates prepared with SNU-C4, U2OS, HEK293T/empty vector, and HEK293T/3×HA-TERT cells by western blot. The HEK293T/3×HA-TERT cell line overexpressing 3×HA-tagged TERT protein was used as a control. Anti-TERT antibodies from Abcam and Rockland, but not those from Novus, detect overexpressed 3×HA-tagged TERT protein, but none of the three anti-TERT antibodies were able to detect endogenous TERT protein. Shown is a representative of multiple attempts. (E) Anti-TERT antibody from Abcam (ab32020) could not detect the TERT protein in cell lysates prepared with SNU-C4 CRC cells treated with FTIN (100 nM) for 1 h or 4 h. Cell lysates prepared with HEK293T/3×HA-TERT cells were used as a control. Shown is the result of a single experiment. (F) Confirmation of telomerase activity in SNU-C4 CRC cells using the TRAP assay. HCT116 CRC and Namalwa cells were used as telomerase-positive controls, and U2OS cells as a telomerase-negative control. The TRAP assay was performed as described in Experimental Method Details. Shown is the result of a single experiment. (G) Confirmation of the efficacy of siRNA targeting *TERT* (siTERT) (*n* = 3). SNU-C4 cells were transfected with siTERT (20 nM) or control (siCon) for 72 h and subsequently treated with FTIN (100 nM) or vehicle (DMSO) for 4 h, and *TERT* mRNA levels were determined by qRT-PCR. (H) TRAP assay result showing increased telomerase activity in SNU-C4 CRC/3×HA-TERT cells overexpressing the 3×HA-tagged TERT protein as compared to SNU-C4/empty vector cells (*n* = 3). Celly lysates prepared from SNU-C4 cells maintained in DMEM medium supplemented with 10% FBS were used as controls. Data are represented as mean ± SD, where appropriate. Statistical significance was assessed by an unpaired *t*-test (A, B, H) and two-way ANOVA (G): *, *p* < 0.05; **, *p* < 0.01; ***, *p* < 0.001; ****, *p* < 0.0001; ns, not significant.

**Figure S7.**
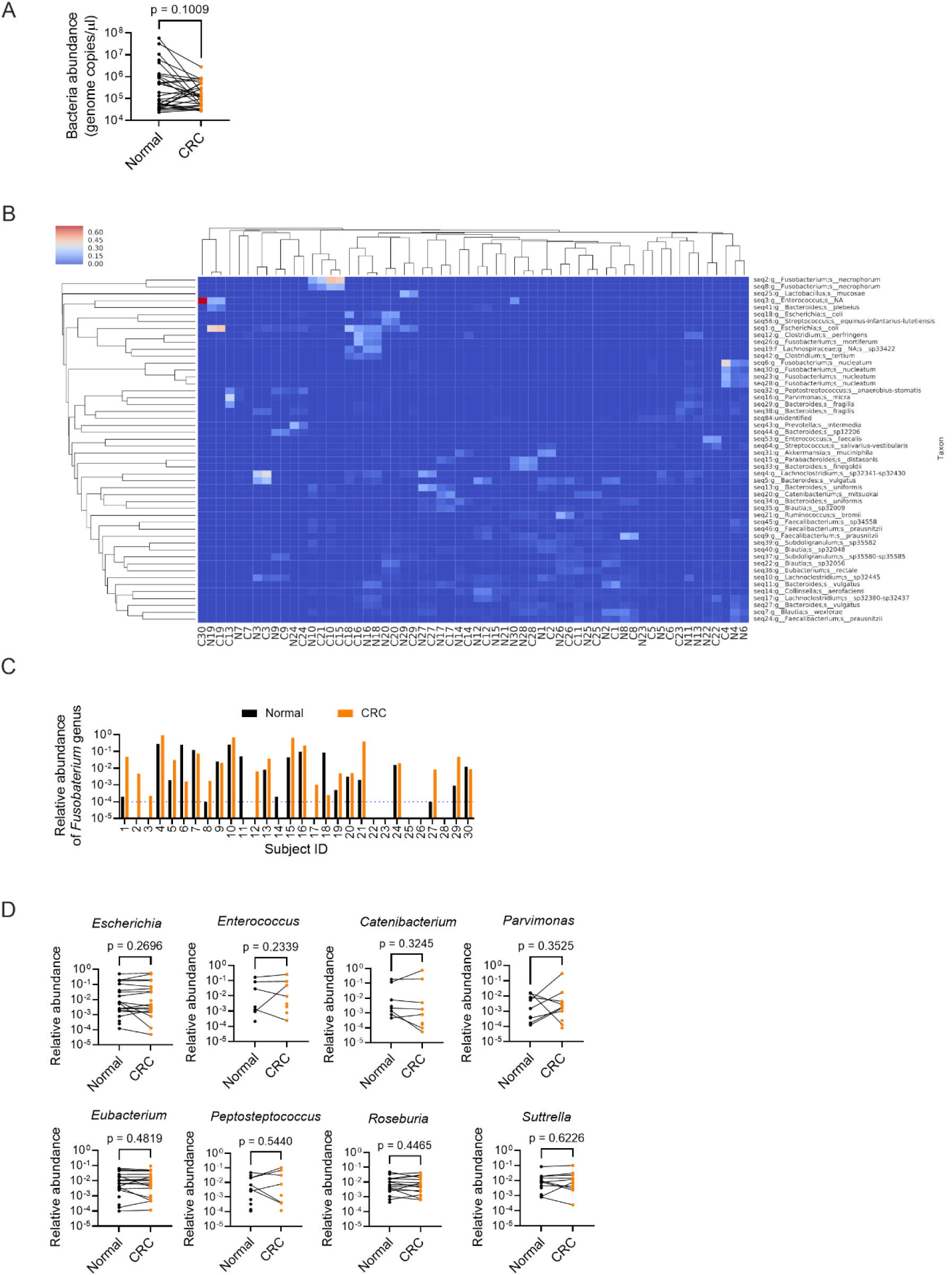
Additional analysis of 16S-seq data for 30 pairs of human CRC and matching normal colon tissues. Related to Figure 7. (A) Absolute abundance of bacterial genomes in the genomic DNA of 60 CRC and adjacent normal colon tissues. An absolute abundance of bacterial genomes was obtained as described in the Experimental Method Details (16S rRNA gene amplicon sequencing). Statistical significance was determined by a paired *t*-test. (B) Taxonomy abundance heatmap with sample clustering indicates that most pairs of CRC- normal colon tissues harbor similar bacterial compositions. The heatmap was created with the top 50 most abundant species identified. Each row represents the abundance for each taxon, with the taxonomy ID shown on the right, and each column represents the abundance for each sample, with the sample ID shown at the bottom. Samples were hierarchically clustered using Bray-Curtis dissimilarity, and taxa were also hierarchically clustered to group those with similar distributions (C) Relative abundance (≥ 0.01%) of the *Fusobacterium* genus in CRC tissues and matching normal colon tissues (30 paired tissues from 30 subjects). The dotted line denotes 0.01%. (D) Relative abundance (≥ 0.01%) of eight genera detected in paired tissues with FTIN levels above the limit of detection (0.01 nmol/g tissue) shows no difference between CRC tissues and matching normal colon tissues. Statistical significance was determined by a paired *t*-test.

**Table S1.** Complete RNA-seq data of four CRC cell lines_(F)TIN vs (D)MSO (Excel file)

**Table S2.**
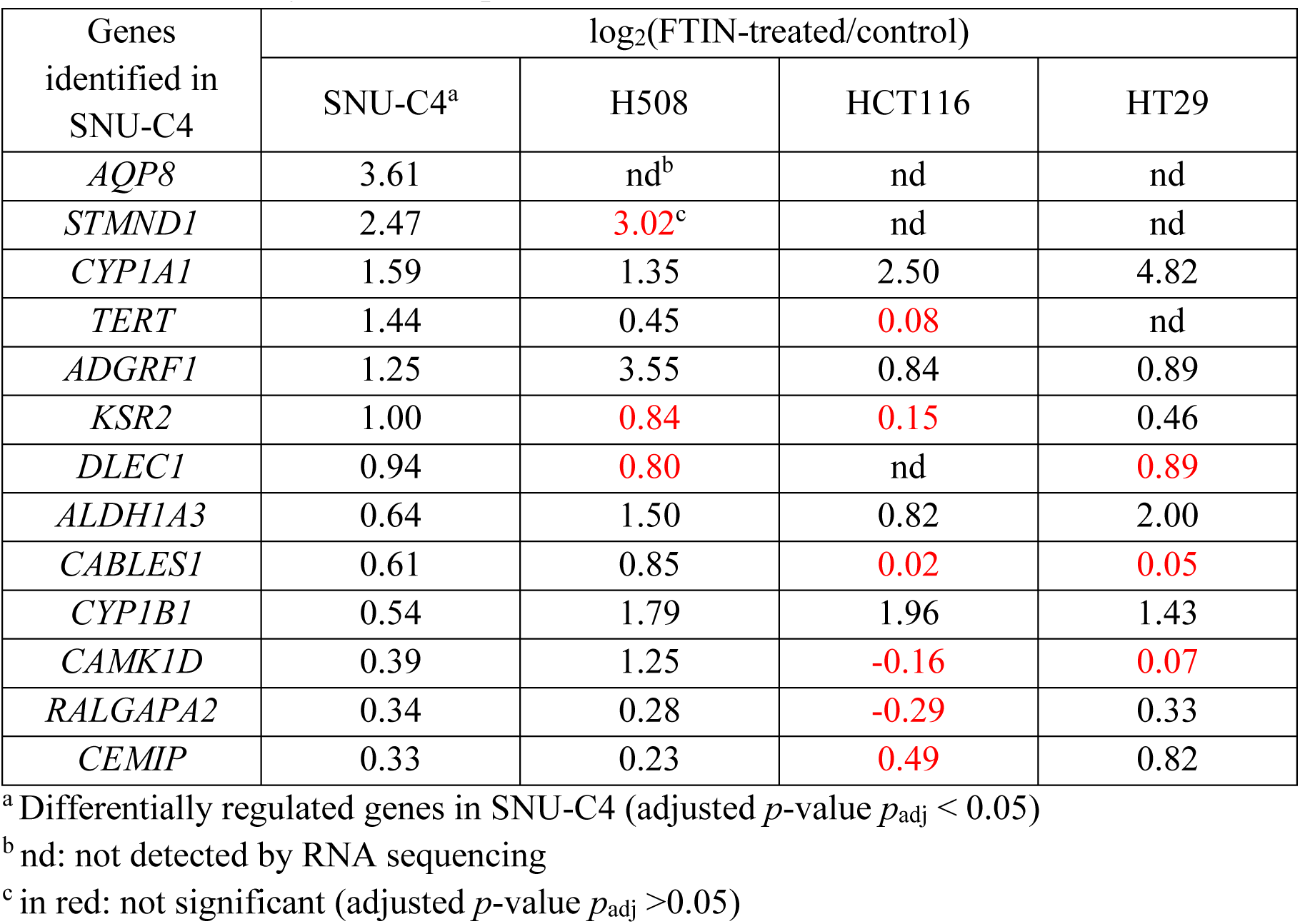
Summary of RNA-seq in four CRC cell lines

**Table S3.** Clinical data for CRC and adjacent normal colon tissues used in this study (Excel file)

**Table S4.**
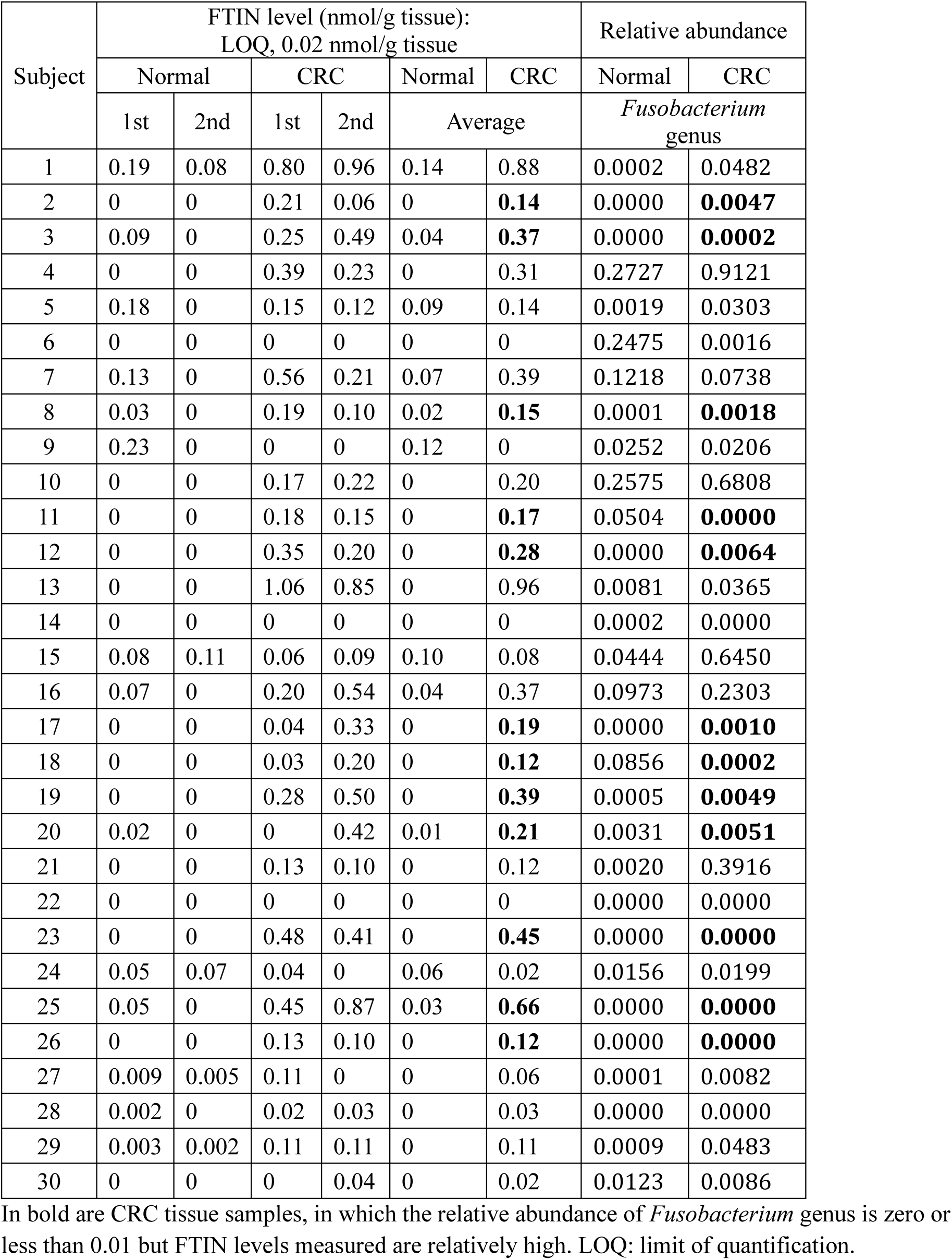
FTIN measurement in human CRC and normal colon tissues

**Table S5.** 16S-seq data showing the relative abundance of bacteria in CRC and normal colon tissues (Excel file)

**Table S6.** FTIN production in previously reported CRC-enriched bacteria

| Genus | Species | FTIN (ng/ml/OD<sub>600</sub>),<br>LOQ* = 0.02 ng/ml |  |
| --- | --- | --- | --- |
|  |  | 1st | 2nd |
| *Bacteroides* | *Bacteroides fragilis* BNCC336948 | < LOQ | < LOQ |
|  | *Bacteroides fragilis* CL07T00C01 | < LOQ | < LOQ |
|  | *Bacteroides thetaiotaomicron* VPI 5482 | 0.74 | 0.74 |
| *Clostridium* | *Clostridium symbiosum* DSM 934 | < LOQ | < LOQ |
|  | *Clostridium symbiosum* WAL-14163 | < LOQ | < LOQ |
| *Dialister* | *Dialister hominis* DSM 109768 | < LOQ | < LOQ |
| *Escherichia* | *Escherichia coli* MG1655 | 0.14 | 0.17 |
| *Fusobacterium* | *Fusobacterium nucleatum* ATCC 23726 | 42.99 | 37.01 |
| *Gemella* | *Gemella morbillorum* VPI 5424 | < LOQ | < LOQ |
| *Granulicatella* | *Granulicatella adiacens* ATCC 49175 | < LOQ | < LOQ |
| *Hungatella* | *Hungatella hathewayi* 1313 | < LOQ | < LOQ |
| *Leptotrichia* | *Leptotrichia hofstadii* DSM 21651 | < LOQ | < LOQ |
|  | *Leptotrichia trevisanii* ATCC 700907 | < LOQ | < LOQ |
| *Parvimonas* | *Parvimonas micra* ATCC 33270 | < LOQ | < LOQ |
| *Peptostreptococcus* | *Peptostreptococcus anaerobius* 4526 | < LOQ | < LOQ |
|  | *Peptostreptococcus anaerobius* DSM 2949 | < LOQ | < LOQ |
|  | *Peptostreptococcus stomatis* W2278 | 0.27 | 0.32 |
| *Porphyromonas* | *Porphyromonas asaccharolytica* ATCC 25260 | 11.03 | 7.24 |
|  | *Porphyromonas gingivalis* ATCC 33277 | 3.52 | 3.15 |
|  | *Porphyromonas gingivalis* F0566 | 5.28 | 5.58 |
|  | *Porphyromonas gingivalis* F0568 | 14.10 | 14.46 |
| *Prevotella* | *Prevotella copri* CB7 | 1.19 | 1.40 |
|  | *Prevotella intermedia* B422 | 0.12 | 0.16 |
| *Selenomonas* | *Selenomonas sputigena* ATCC 35185 | < LOQ | < LOQ |
| *Solobacterium* | *Solobacterium moorei* CC14Y | < LOQ | < LOQ |
|  | *Solobacterium moorei* CC57E | < LOQ | < LOQ |
|  | *Solobacterium moorei* DSM 22971 | < LOQ | < LOQ |
| *Streptococcus* | *Streptococcus dysgalactiae* NCTC 8543 | < LOQ | < LOQ |
|  | *Streptococcus gallolyticus* ATCC 49147 | < LOQ | < LOQ |
|  | *Streptococcus gallolyticus* ATCC 9809 | < LOQ | < LOQ |
| *Veillonella* | *Veillonella parvula* Te3 | < LOQ | < LOQ |

\*LOQ: limit of quantification.

**Table S7.**
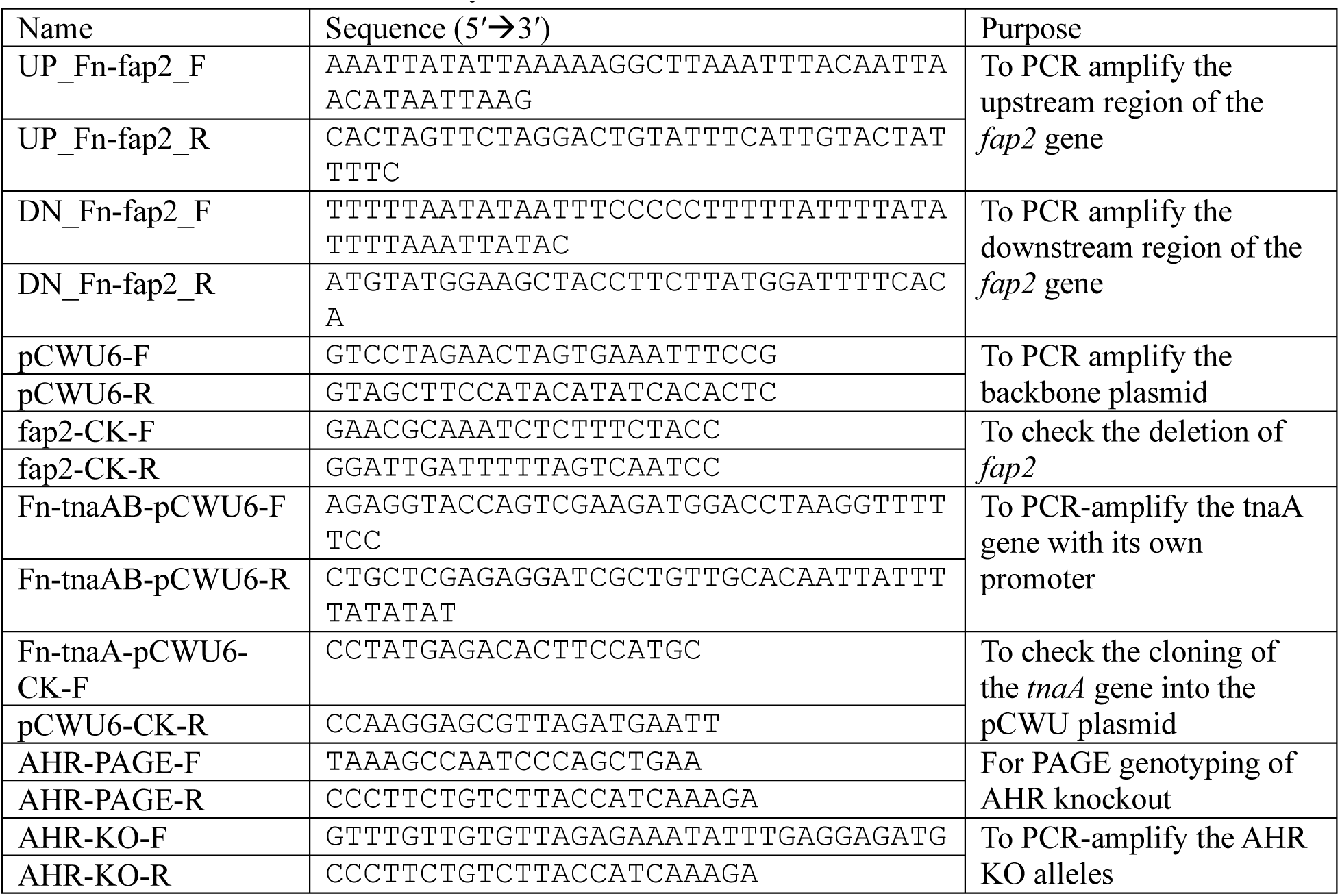
Primers used in this study

**Supplemental Texts**

## SUPPLEMENTAL RESULTS

**Isolation and identification of AHR-activating metabolites of *Fusobacterium nucleatum*** *Fusobacterium nucleatum* (*Fn*) was cultivated anaerobically in 2 L of M9CT medium for 72 h. At harvest, the culture supernatant was filtered and extracted by liquid-liquid partition using ethyl acetate (1:1). The organic extracts were fractionated by vacuum liquid chromatography (VLC) with Diaion HP-20SS resin using a step gradient consisting of isopropyl alcohol and H_2_O. The original extracts and VLC fraction F4 (70% isopropyl alcohol) demonstrated the highest AHR-activating activity in the AHR reporter assay. The bioactive VLC fraction (F4) was further fractionated by RP-HPLC using MeOH and re-evaluated in the same reporter assay. The bioactive subfractions with the greatest yield (SF7-11) were further purified using RP-HPLC using acetonitrile to yield an isolate that demonstrated potent bioactivity in the AHR reporter assay. The HPLC isolate was found to be a mixture of two major compounds (**1** and **2**) by HR-ESI-LC-MS/MS (**Supporting Figure S1**). The MS2 spectra for the ions corresponding to compound **1**, *m/z* 364.14 [M+H]^+^, and compound **2**, *m/z* 341.13 [M+Na]^+^ were selected for analysis and natural product database search (**Supplemental Method Details**, Dereplication using the SIRIUS workbench). For compound **1**, the best fit molecular formula, C_24_H_17_N_3_O, was calculated, and two natural products (>60% similarity match) were identified that correlated to the MS2 spectrum of **1**: 2,2-bis(1*H*-indol-3-yl)-1*H*-indol-3-one (**1**) with 66.8% similarity match and 3,3-bis(1*H*-indol- 3-yl)-1*H*-indol-2-one (trisindoline, **3**) with 68.3% similarity match. For compound **2**, the best fit molecular formula, C_20_H_18_N_2_O_2_, was calculated, and one potential match was identified, indicating the known compound, streptindole (similarity match, 70.7%). These results indicated the presence of two bisindole alkaloids **1** and **2** in *Fn* culture (**Chart 1**).

In previous studies, compound **1** was isolated from the syntrophic bacterium *Symbiobacterium thermophilum* and called BII as the abbreviation for 2,2-bis(3’-indolyl)indoxyl,^1^ and compound **2** was isolated from the facultative anaerobic bacterium *Streptococcus faecium* (reclassified as *Enterococcus faecium*^2^) and was given the traditional name streptindole.^3^ We named compound **1** as fusotrisindoline. For simplicity, we used the traditional names of compounds **1**, **2,** and **3** (trisindoline) hereafter.

Chart 1. Structures of compounds **1** and **2** isolated from *Fusobacterium nucleatum*, as well as **3**.

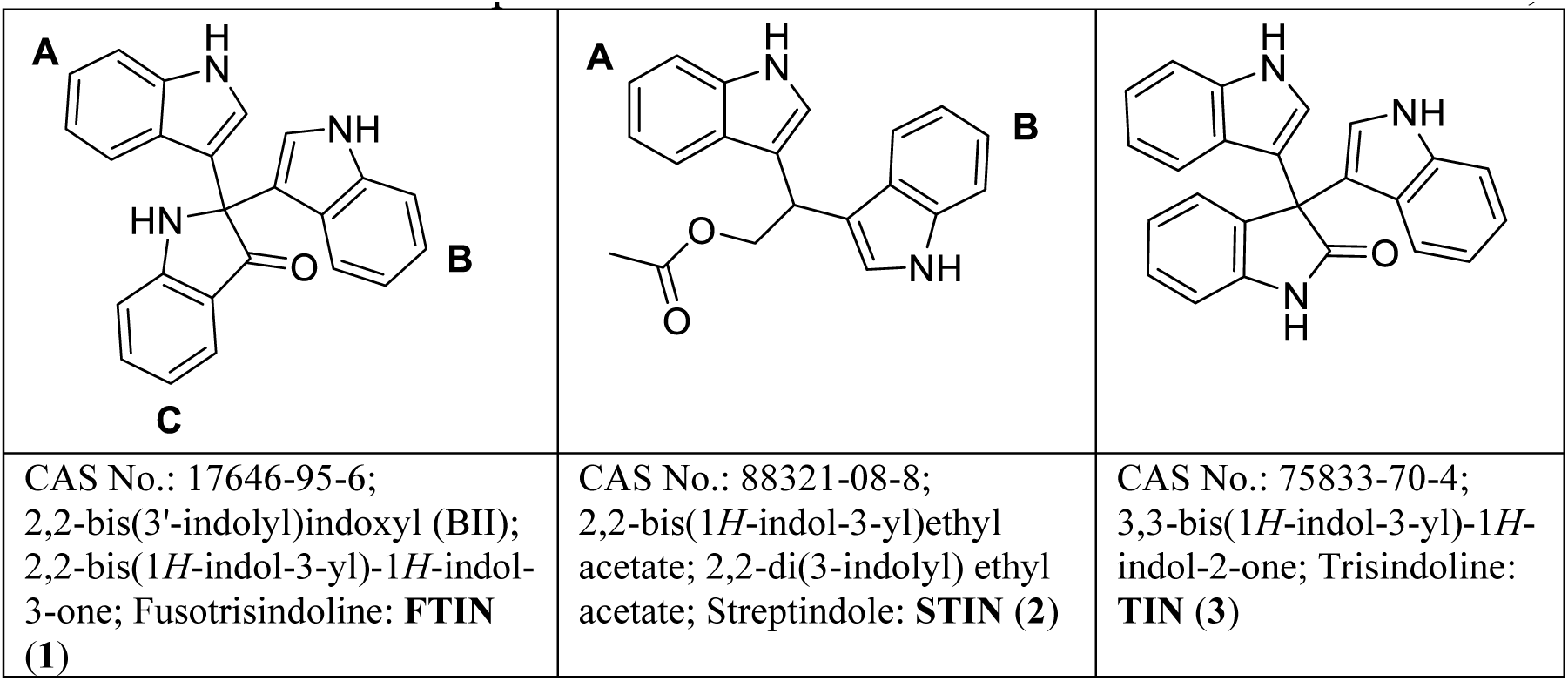

### Validation of FTIN and STIN structures

To confirm the identity of *Fn*-derived compounds **1** and **2**, the two positional isomers, fusotrisindoline (**1**, FTIN) and trisindoline (**3**, TIN), as well as streptindole (**2**, STIN), were custom and in-house synthesized (**Supporting Figures S2-S6**). Using analytical HPLC, we confirmed that the retention time (t_R_) of compound **1** in the *Fn*-derived HPLC isolate matches that of synthesized FTIN: synthesized FTIN (t_R_ 13.3 min.), synthesized TIN (t_R_ 12.4 min.), *Fn*-derived compound **1** (t_R_ 13.3 min) (**Supporting Figure S7**). The ^1^H NMR spectra for synthesized FTIN and STIN were identical to those of *Fn*-derived compounds **1** and **2** (**Supporting Figures S8-S13**), respectively, and matched previously reported ^1^H NMR signals in earlier studies (**Supporting** **Table 1**). All major signals in the ^1^H NMR spectrum of the *Fn*-derived HPLC isolate could be correlated to FTIN (**1**) and STIN (**2**) (**Supporting Figures S14** and **S15**). Using a standard ^1^H NMR experiment, FTIN (**1**) and STIN (**2**) in the *Fn*-derived HPLC isolate were found to be in an approximate 1:1 ratio (**Supporting Figure S16**). These results validate the chemical identities of two *Fn*-derived metabolites as FTIN and STIN.

### Identification of trisindoline as an *Fn* metabolite

TIN is a regioisomer of FTIN (**Chart 1** above). Our initial purpose in chemically synthesizing TIN was to ensure that *Fn*-derived compound **1** (i.e., FTIN) was not TIN during the validation of chemical identity. On the other hand, since TIN was previously shown to be produced, together with FTIN, by the same bacteria,^4–6^ we attempted to detect TIN while determining FTIN and STIN levels in *Fn* cultures using LC/MS. As a result, we found the presence of TIN in quantifiable amounts in *Fn* M9CT cultures (see **Main Text**, **Figure 3B**).

## SUPPLEMENTAL METHOD DETAILS

### Extraction and Fractionation

Vacuum liquid chromatography (VLC) was carried out using a Waters extraction manifold. High-performance liquid chromatography (HPLC) experiments were carried out using an Agilent 1100 HPLC system equipped with an Agilent 1100 quaternary pump, autosampler, column oven, photodiode array detector (PDA), and fraction collector. Chromatographic data were collected and analyzed using ChemStation software (Agilent). All fractions and HPLC isolates were dried *in vacuo* using a Savant Speedvac system (Thermo Fisher Scientific). A crude extract was prepared by a liquid-liquid partition with ethyl acetate using cell-free filtrate from the culture supernatant of *Fn*. The ethyl acetate extract was fractionated against Diaion HP-20SS using vacuum liquid chromatography (VLC) with a step gradient consisting of 20 mL fractions of isopropylalcohol (IPA):H_2_O, 0:100 (F1), 20:80 (F2), 40:60 (F3), 70:30 (F4), 90:10 (F5) and 100:0 (F6), followed by a wash using ethyl acetate and acetone, respectively. Fractions were evaporated to dryness *in vacuo*. The F4 (70% IPA, 10.8 mg) fraction demonstrated the greatest AHR activation in the AHR reporter assay and was further fractionated by reverse phase (RP)-HPLC. The F4 fraction was prepared using solid phase extraction (SPE) with a C_18_ cartridge (Agela Cleanert, C_18_, 100 mg, S181001-L) followed by syringe-driven ultrafiltration (0.22 μm filter, PTFE, Millex, SLLG025SS) prior to analysis. HPLC subfractionation was performed using a monolithic C_18_ column (Phenomenex, Onyx, 100 x 4.6 mm, P/N CH0-7643) operating at 37°C with a flow rate at 2.0 mL/min and a mobile phase beginning with a ramp from 35-50% B (A = 5% MeOH:H_2_O, B = 100% MeOH) from 0.0-0.5 min. and then a gradient consisting of 50-80% B over 9.5 min, ramping to 100% B from 10.0-11.0 min, holding at 100% B from 11.0-13.0 min., followed by re-equilibration to starting conditions. HPLC fractions were collected every 30 seconds between 0.5 and 12.5 min., yielding 24 subfractions. Due to low yield, subfractions 7-11 were combined and further purified using a C_18_ column (Phenomenex, Kinetex, 250 × 4.6 mm, P/N 00G-4633-E0) operating at 40°C and a flow rate of 1.0 mL/min with a gradient method consisting of an initial hold at 25% B (A = 5% CH_3_CN + 0.1% formic acid (FA), B = 95% CH_3_CN-H_2_O + 0.1% FA) from 0.0-3.0 min., followed by a gradient from 25-60% B between 3.0-56.0 min., followed by a hold at 60% B from 56.0-61.0 min, followed by a ramp to 95% B from 61.0-61.5 min., with a hold at 95% B from 61.5-70.5 min., followed re-equilibration to starting conditions. HPLC fractions were collected every 60 seconds from 19.0 to 45.0 min, yielding 26 subfractions. The major analytes coeluted in subfractions 10-12 (0.20 mg; grey solid; compound **1**, t_R_ = 29.3 min.; compound **2**, t_R_ = 30.0 min.) and were combined for further analysis.

### High-Resolution Mass Spectrometry and Nuclear Magnetic Resonance Spectroscopy

The HR-ESI-LC-MS/MS (high-resolution electrospray ionization-liquid chromatography-mass spectrometry) analysis of the HPLC isolates was conducted at UIC using an ImpactII (Bruker) Qq- TOF mass spectrometer equipped with an electrospray ionization source operating in positive ionization mode, connected by a capillary to a Nexera X2 multimodule UHPLC system (Shimadzu). Chromatographic separation was achieved using a C_18_ column (Phenomenex, Kinetex, 50 X 2.1 mm, P/N 00B-4475-AN) operating at 40°C with a flow rate of 0.3 mL/min using the following chromatographic program: initial hold at 10% B from 0.0-0.5 min., followed by a gradient from 10-100% B from 0.5-5.5 min., with a hold at 100% B from 5.5-7.0 min., followed by re-equilibration to starting conditions.

The MS method was calibrated before and during analysis using a solution of sodium formate (VWR). MS1 detection was optimized for ions with a mass to charge ratio (*m/z*) of 50- 2000 with a sampling rate of 12.0 Hz. At the electrospray source, the end plate offset was set to 250 V, capillary voltage of 4500 V, nebulizer at 4.0 bar, drying gas at 12 L/min with a drying temperature of 225°C. Transfer energy (RF) was set at 350.0 Vpp for ion funnel 1 and 2. The hexapole operated at 80.0 Vpp, and the quadrupole ion energy was set at 4.0 eV with a low mass cut-off of 50.00 *m/z*. At the collision cell, the collision energy (CE) was 5.0 eV with a pre-pulse storage of 7.0 μs. Fixed data acquisition was set to 0.68 Hz, and tandem spectra were collected using a stepping program: base *m/z* 500 (charge state 1), CE=35.0 eV; base *m/z* 1000 (charge state 1), CE=50.0 eV; base *m/z* 2000 (charge state 1), CE=70.0 eV. The auto MS/MS setting was used to collect MS2 spectra, collecting data for five precursor ions per duty cycle, requiring an absolute threshold of 316 cts. Ions were actively excluded after collecting three spectra, reconsidering the precursor ion if the current intensity was 2.0× greater than the previous intensity. The software, DataAnalysis (Bruker, version 4.4) was used for post-run calibration and analysis of MS data.

NMR data were collected using a Bruker Avance AVIII 600 MHz spectrometer equipped with a 5 mm CP DCH probe operating at 600.15 MHz for ^1^H. NMR experiments were acquired using Bruker pulse programs. All chemical shifts were referenced to the residual solvent signal (acetone-*d*_6_, δ_H_ 2.05).

### Dereplication using the SIRIUS workbench

The post-run calibrated datafile (.d) was exported from DataAnalysis (Bruker, version 4.4) as an mzXML file and was imported into the SIRIUS workbench (version 5.8.3). Two major analytes were observed in the LC-MS chromatogram of the HPLC isolate; therefore, the MS2 spectra for the ions corresponding to **1**, *m/z* 364.14 [M+H]^+^, and **2**, *m/z* 341.13 [M+Na]^+^ were selected for analysis. All default parameters were applied, including the SIRIUS molecular formula identification module,^7^ CSI:FingerID fingerprint prediction,^8^ and structure database search modules (with the COCONUT,^9^ GNPS,^10^ Natural Products,^11^ and PubChem^12^ databases selected), as well as the CANOPUS compound class prediction module.^13^

### Synthesis of FTIN, FTIN-^13^C_6_, STIN, and TIN

FTIN (>96% purity) and stable-isotope labeled FTIN (FTIN-^13^C_6_, >99% purity) were custom synthesized by WUXI, and STIN (>95% purity) by KareBay Bio. FTIN and TIN (>95% purity) were also in-house synthesized (see **Main Method Details**).

#### Chromatographic comparison of synthesized standards and *Fn*-derived isolates

Approximately 50 μg of synthesized FTIN and TIN, as well as 3.75 μg of *Fn*-derived HPLC isolate containing **1** and **2,** were analyzed by analytical RP-HPLC using a C_18_ column (Phenomenex, Kinetex, 250 x 4.6 mm, P/N 00G-4633-E0) operating at a flow rate of 1.0 mL/min and a temperature of 40°C. A gradient analysis was performed with an initial hold at 20% B (A = 5% MeCN, B = 95% MeCN) from 0.0-3.0 min., followed by a gradient of 20-100% B between 3.0-33.0 min., a wash at 100% B between 33.0-39.0 min., followed by re-equilibration to starting conditions. The compounds had the following retention times: synthesized FTIN (t_R_ 13.3 min.), synthesized TIN (t_R_ 12.4 min.), *Fn*-derived **1** (t_R_ 13.3) (**Supporting Figure S7**).

---

**Supporting Table 1.** Comparison of ^1^H NMR data obtained in this study with previously reported data for compounds **1** (FTIN) and **2** (STIN)

| Position | Observed in this study | Synthesized in this study | Reported (Watsuji et al., 2007)<sup>1</sup> | Observed in this study | Synthesized in this study | Reported (Osawa & Namiki, 1983)<sup>3</sup> |
| --- | --- | --- | --- | --- | --- | --- |
|  | As in *Fn* mixture of **1** and **2** |  |  | As in *Fn* mixture of **1** and **2** |  |  |
|  | 2,2-di(3-indoyl)-3-indolone (**1**), Acetone-*d*<sub>6</sub>, 600 MHz | 2,2-di(3-indoyl)-3-indolone (**1**), Acetone-*d*<sub>6</sub>, 600 MHz | 2,2-di(3-indoyl)-3-indolone (**1**), Acetone-*d*<sub>6</sub>, 500 MHz | Streptindole (**2**), Acetone-*d*<sub>6</sub>, 600 MHz | Streptindole (**2**), Acetone-*d*<sub>6</sub>, 600 MHz | Streptindole (**2**), Acetone-*d*<sub>6</sub>, 360 MHz |
|  | δ<sub>H</sub>, ppm | δ<sub>H</sub>, ppm | δ<sub>H</sub>, ppm | δ<sub>H</sub>, ppm | δ<sub>H</sub>, ppm | δ<sub>H</sub>, ppm |
| A-1, NH | 10.17 (1H, bs) | 10.23 (1H, s) | 10.20 (1H, bs) | 10.07 (1H, bs) | 10.05 (1H, s) | 10.06 (1H, s) |
| A-2 | 7.24 (1H, dd, J=2.6 Hz) | 7.24 (1H, d, J=2.6 Hz) | 7.22 (1H, d, J=2.6 Hz) | 7.22 (1H, d, J=2.3 Hz) | 7.22 (1H, d, J=2.3 Hz) | 7.23 (1H, s) |
| A-3 | - | - | - | - | - | - |
| A-3a | - | - | - | - | - | - |
| A-4 | 7.45 (1H, d, J=8.2 Hz) | 7.46 (1H, dd, J=8.1, 1.0 Hz) | 7.43 (1H, dd, J=8.1, 10.0 Hz) | 7.60 (1H, d, J=7.9 Hz) | 7.60 (1H, d, J=7.8 Hz) | 7.62 (1H, d, J=7.8 Hz) |
| A-5 | 6.84 (1H, ddd, J= 7.0, 8.0, 1.0 Hz) | 6.85 (1H, ddd, J=7.0, 8.1, 1.0 Hz) | 6.82 (1H, ddd, J=7.0, 8.1, 10.0 Hz) | 6.95 (1H, ddd, J=7.0, 8.0, 1.0 Hz) | 6.95 (1H, ddd, J=6.9, 7.9, 1.0 Hz) | 6.96 (1H, dd, J=6.8, 7.8 Hz) |
| A-6 | 7.04 (1H, ddd, J=8.0, 7.0, 1.0 Hz) | 7.05 (1H, ddd, 7.0, 8.0, 1.0 Hz) | 7.04 (1H, ddd, J=7.0, 8.1, 10.0 Hz) | 7.06 (1H, ddd, 7.0, 8.2, 1.0 Hz) | 7.07 (1H, ddd, J=8.3, 7.0, 1.0 Hz) | 7.08 (1H, dd, J=8.3, 6.8 Hz) |
| A-7 | 7.39 (1H, d, J=8.0 Hz) | 7.39 (1H, dd, J=8.1, 1.0 Hz) | 7.35 (1H, dd, J=8.1, 10.0 Hz) | 7.38 (1H, d, J=8.2 Hz) | 7.38 (1H, d, J=8.3 Hz) | 7.40 (1H, d, J=8.3 Hz) |
| A-7a | - | - | - | - | - | - |
| B-1, NH | 10.17 (1H, bs) | 10.23 (1H, s) | 10.20 (1H, bs) | 10.07 (1H, bs) | 10.05 (1H, s) | 10.06 (s) |
| B-2 | 7.24 (1H, dd, J=2.6 Hz) | 7.24 (1H, d, J=2.6 Hz) | 7.22 (1H, d, J=2.6 Hz) | 7.22 (1H, d, J=2.3 Hz) | 7.22 (1H, d, J=2.3 Hz) | 7.23 (1H, s) |
| B-3 | - | - | - | - | - | - |
| B-3a | - | - | - | - | - | - |
| B-4 | 7.45 (1H, d, J=8.2 Hz) | 7.46 (1H, dd, J=8.1, 1.0 Hz) | 7.43 (1H, dd, J=8.1, 10.0 Hz) | 7.60 (1H, d, J=7.9 Hz) | 7.60 (1H, d, J=7.8 Hz) | 7.62 (1H, d, J=7.8 Hz) |
| B-5 | 6.84 (1H, ddd, J= 7.0, 8.0, 1.0 Hz) | 6.85 (1H, ddd, J=7.0, 8.1, 1.0 Hz) | 6.82 (1H, ddd, J=7.0, 8.1, 10.0 Hz) | 6.95 (1H, ddd, J=7.0, 8.0, 1.0 Hz) | 6.95 (1H, ddd, J=6.9, 7.9, 1.0 Hz) | 6.96 (1H, dd, J=6.8, 7.8 Hz) |
| B-6 | 7.04 (1H, ddd, J=8.0, 7.0, 1.0 Hz) | 7.05 (1H, ddd, J=7.0, 8.0, 1.0 Hz) | 7.04 (1H, ddd, J=7.0, 8.1, 10.0 Hz) | 7.06 (1H, ddd, J=7.0, 8.2, 1.0 Hz) | 7.07 (1H, ddd, J=8.3, 7.0, 1.0 Hz) | 7.08 (1H, dd, J=8.3, 6.8 Hz) |
| B-7 | 7.39 (1H, d, J=8.0 Hz) | 7.39 (1H, dd, J=8.1, 1.0 Hz) | 7.35 (1H, dd, J=8.1, 10.0 Hz) | 7.38 (1H, d, J=8.2 Hz) | 7.38 (1H, d, J=8.3 Hz) | 7.40 (1H, d, J=8.3 Hz) |
| B-7a | - | - | - | - | - | - |
| C-1, NH | 7.14 (1H, bs) | 7.20 (1H, bs) | 7.10 (1H, bs) | - | - | - |
| C-2 | - | - | - | - | - | - |
| C-3 | - | - | - | - | - | - |
| C-3a | - | - | - | - | - | - |
| C-4 | 7.56 (1H, d, J=7.7 Hz) | 7.56 (1H, d, J=7.7 Hz) | 7.53 (1H, dd, J=1.3, 7.7 Hz) | - | - | - |
| C-5 | 6.80 (1H, ddd, J=7.0, 8.0, 1.0 Hz) | 6.80 (1H, ddd, J=8.1, 7.0, 1.0 Hz) | 6.77 (1H, ddd, J=0.8, 7.1, 7.7 Hz) | - | - | - |
| C-6 | 7.52 (1H, ddd, J=7.0, 8.0, 1.3 Hz) | 7.51 (1H, ddd, J=1.3, 7.1, 8.1 Hz) | 7.48 (1H, ddd, J=1.3, 7.0, 8.1, 10.0 Hz) | - | - | - |
| C-7 | 7.05 (1H, dd, J=8.3, 0.87 Hz) | 7.04 (1H, dd, J=0.8, 8.4 Hz) | 7.02 (1H, dd, J=0.8, 8.4 Hz) | - | - | - |
| C-7a | - | - | - | - | - | - |
| Methine (C8) | - | - | - | 4.94 (1H, t, J=7.0 Hz) | 4.94 (1H, t, J=7.0 Hz) | 4.93 (1H, t, J=6.9 Hz) |
| Methylene (C9) | - | - | - | 4.72 (2H, d, J=7.0 Hz) | 4.72 (2H, d, J=7.0 Hz) | 4.72 (2H, d, J=6.9 Hz) |
| Acetyl CH<sub>3</sub> (C10) | - | - | - | 1.92 (3H, s) | 1.92 (3H, s) | 1.92 (3H, s) |

## SUPPORTING FIGURES

**S1.**
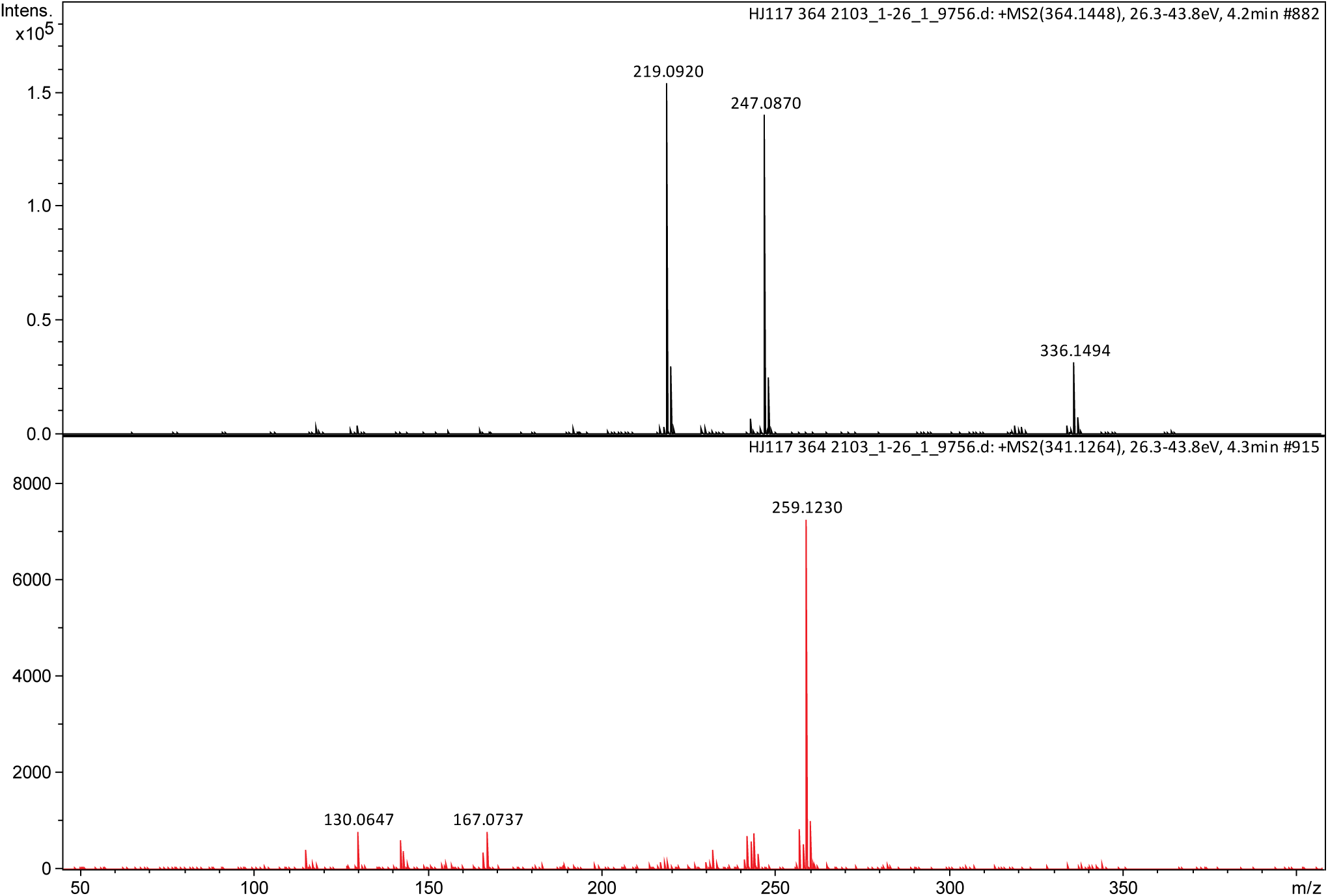
HR-ESI-LC-MS/MS spectra of *Fn*-derived compounds **1** and **2**.

**S2.**
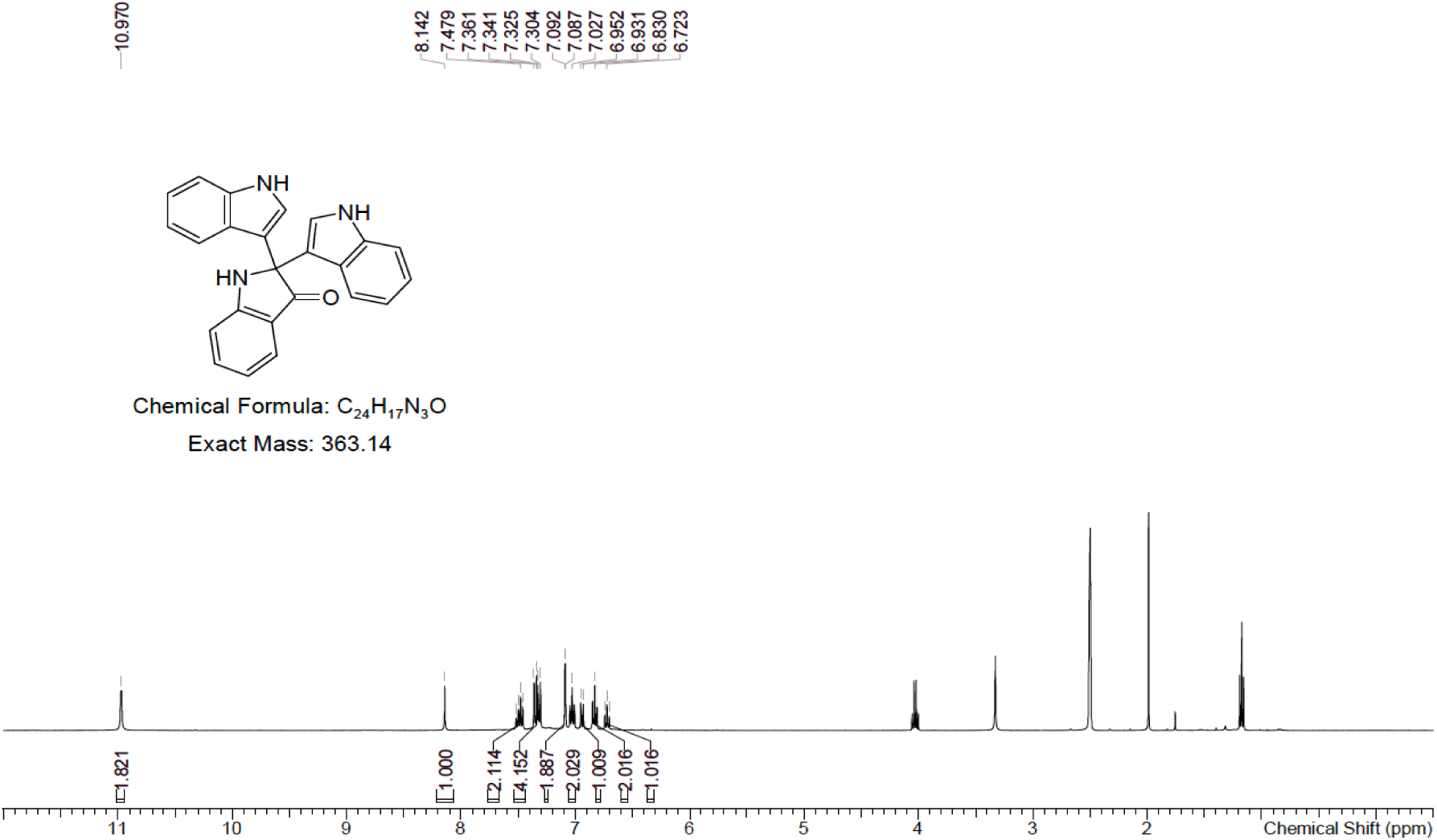
^1^H NMR spectrum (DMSO-*d*_6_) of custom-synthesized FTIN (**1**) provided by the company WUXI.

**S3.**
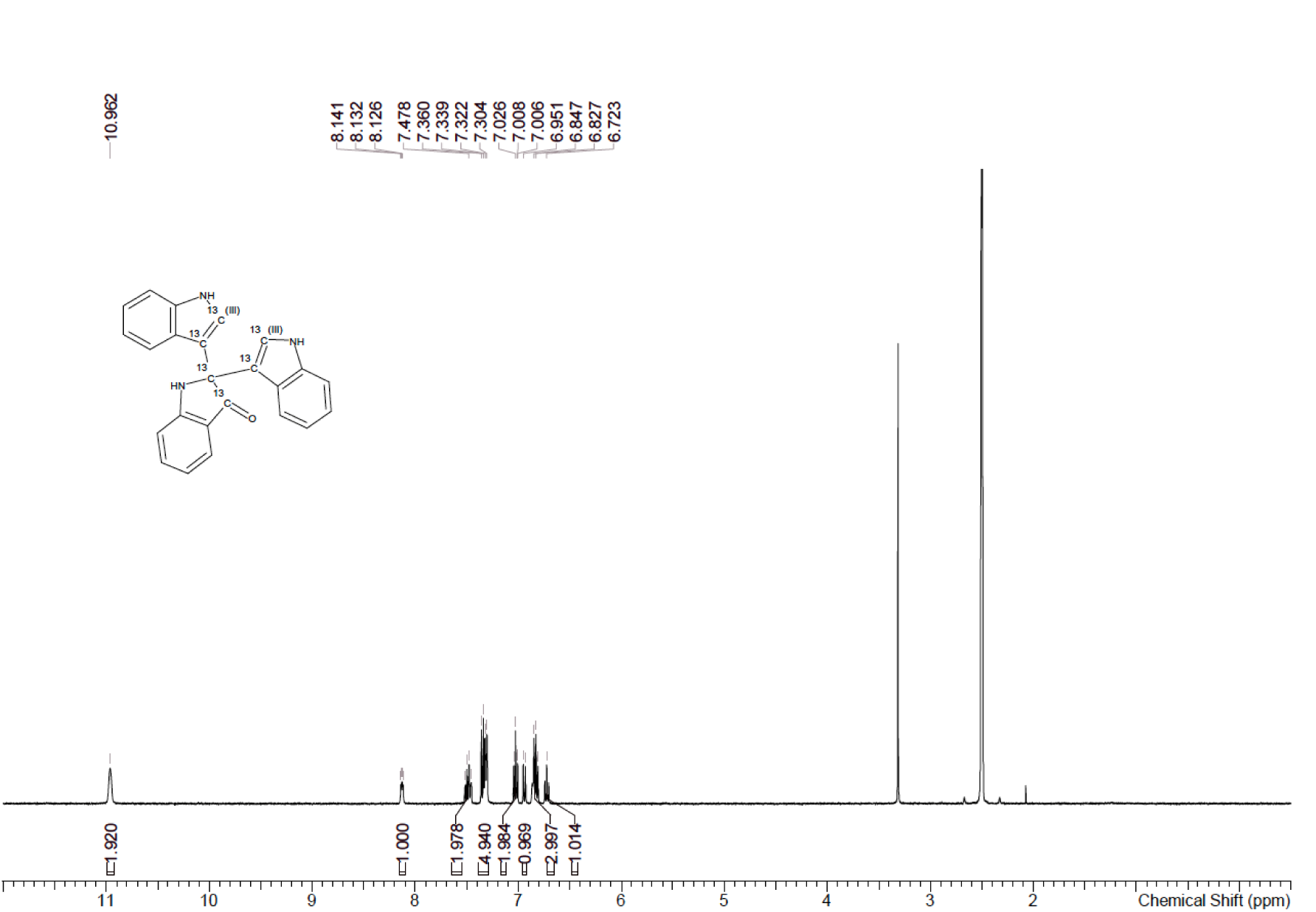
^1^H NMR spectrum (DMSO-*d*_6_) of custom-synthesized isotope-labelled FTIN-^13^C_6_ (**1**) provided by the company WUXI

**S4.**
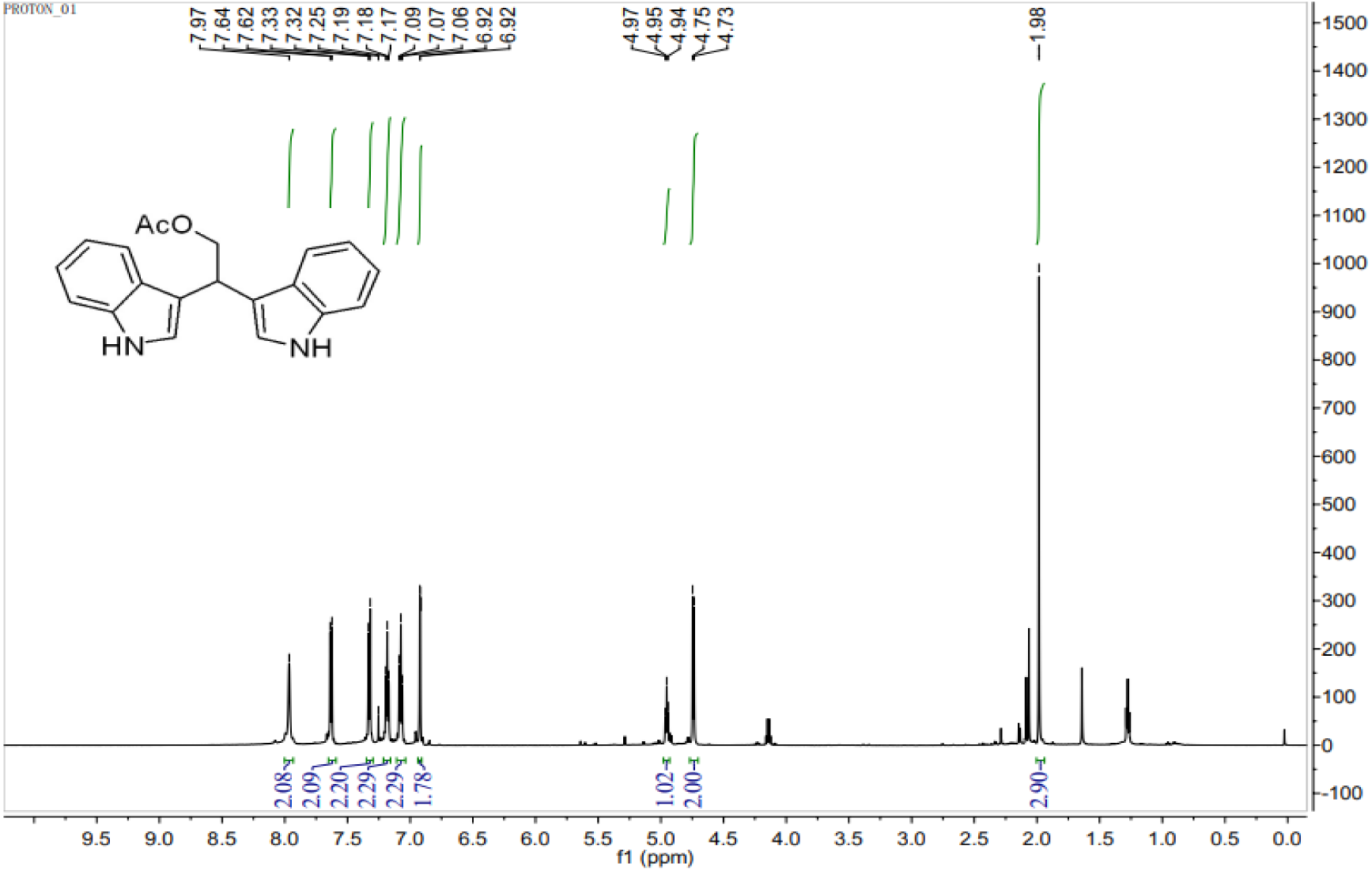
^1^H NMR spectrum (DMSO-*d*_6_) of custom-synthesized STIN (**2**) provided by the company KareBay Bio.

**S5.**
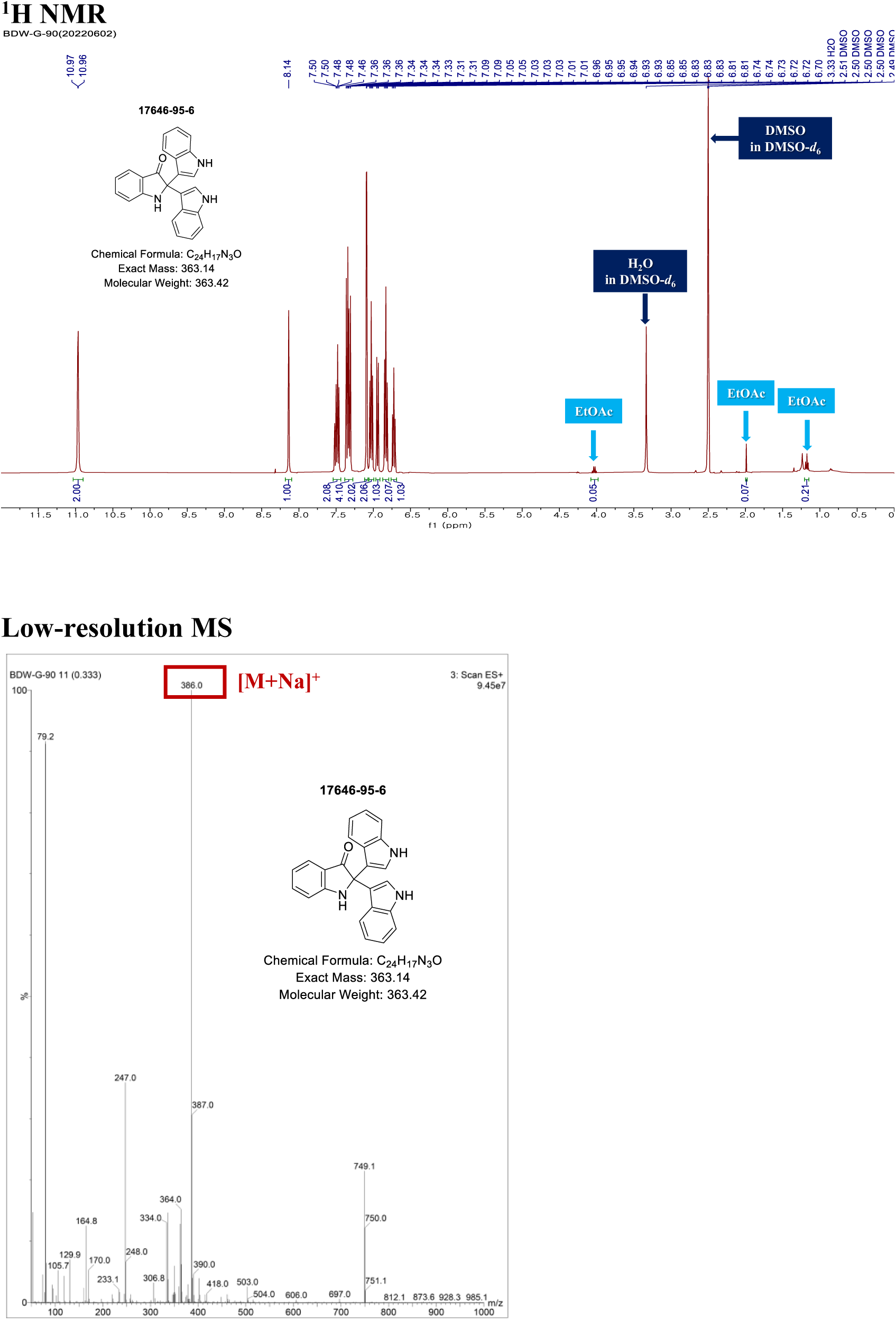
^1^H NMR spectrum (DMSO-*d*_6_) and low-resolution mass spectrometry of in-house synthesized FTIN (**3**).

**S6.**
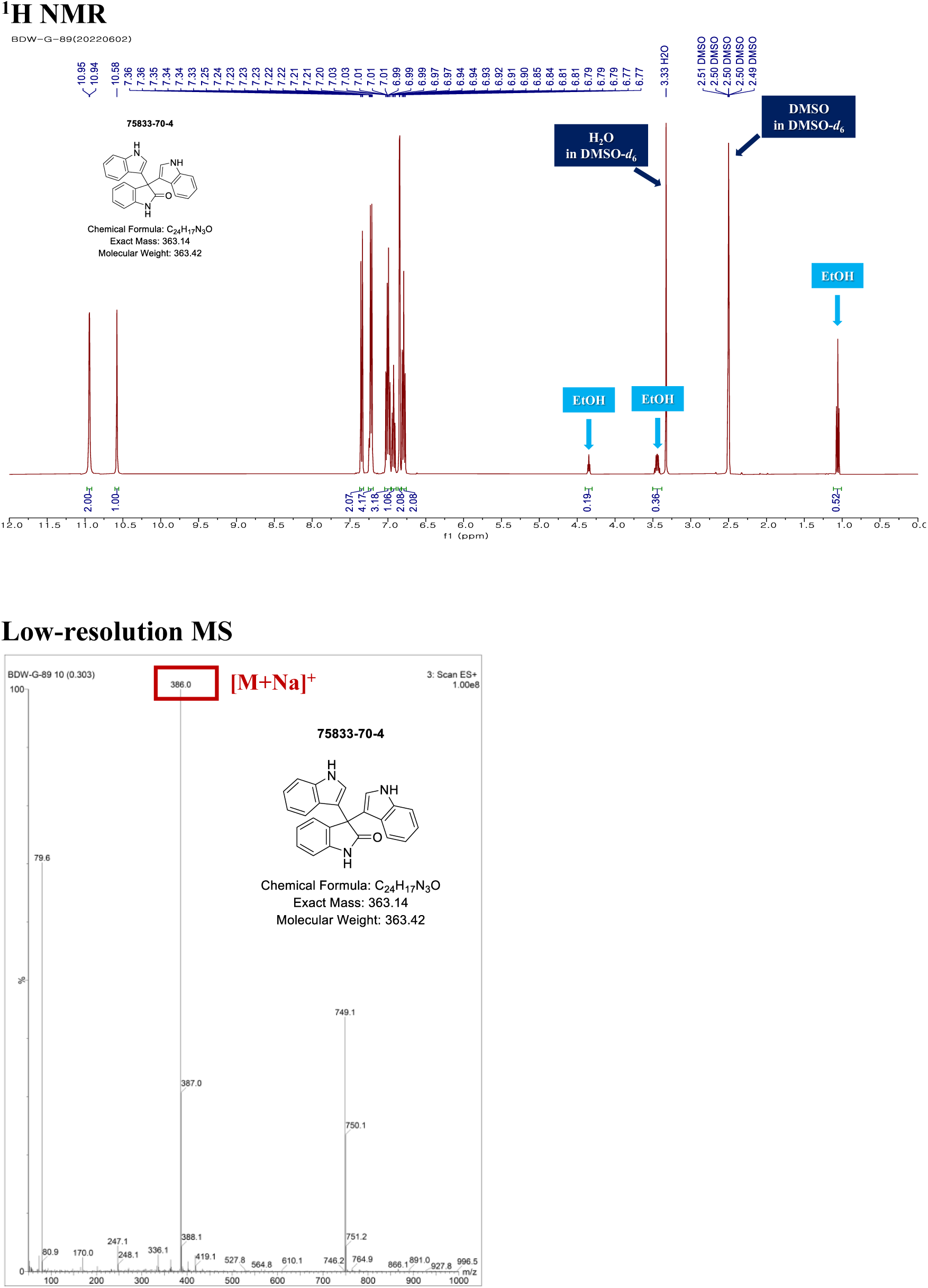
^1^H NMR spectrum (DMSO-*d*_6_) and low-resolution mass spectrometry of in-house synthesized TIN (**3**).

**S7.**
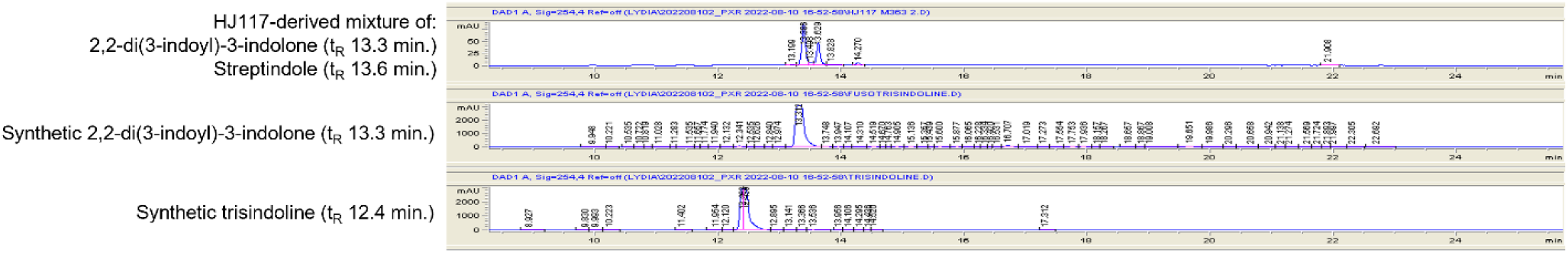
HPLC chromatograms comparing t_R_ of *Fn*-derived **1** to synthesized FTIN and TIN.

**S8.**
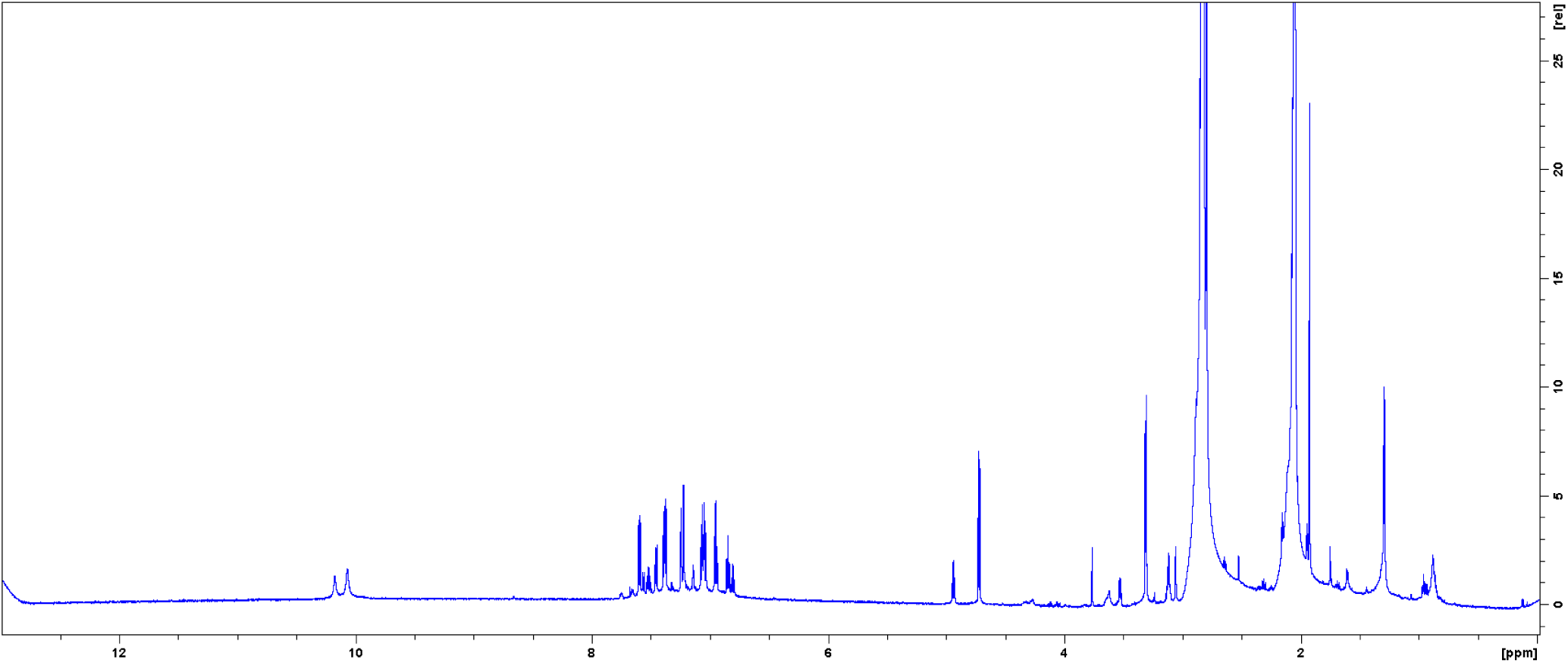
^1^H NMR spectrum (acetone-*d*_6_) of *Fn*-derived mixture of **1** and **2**.

**S9.**
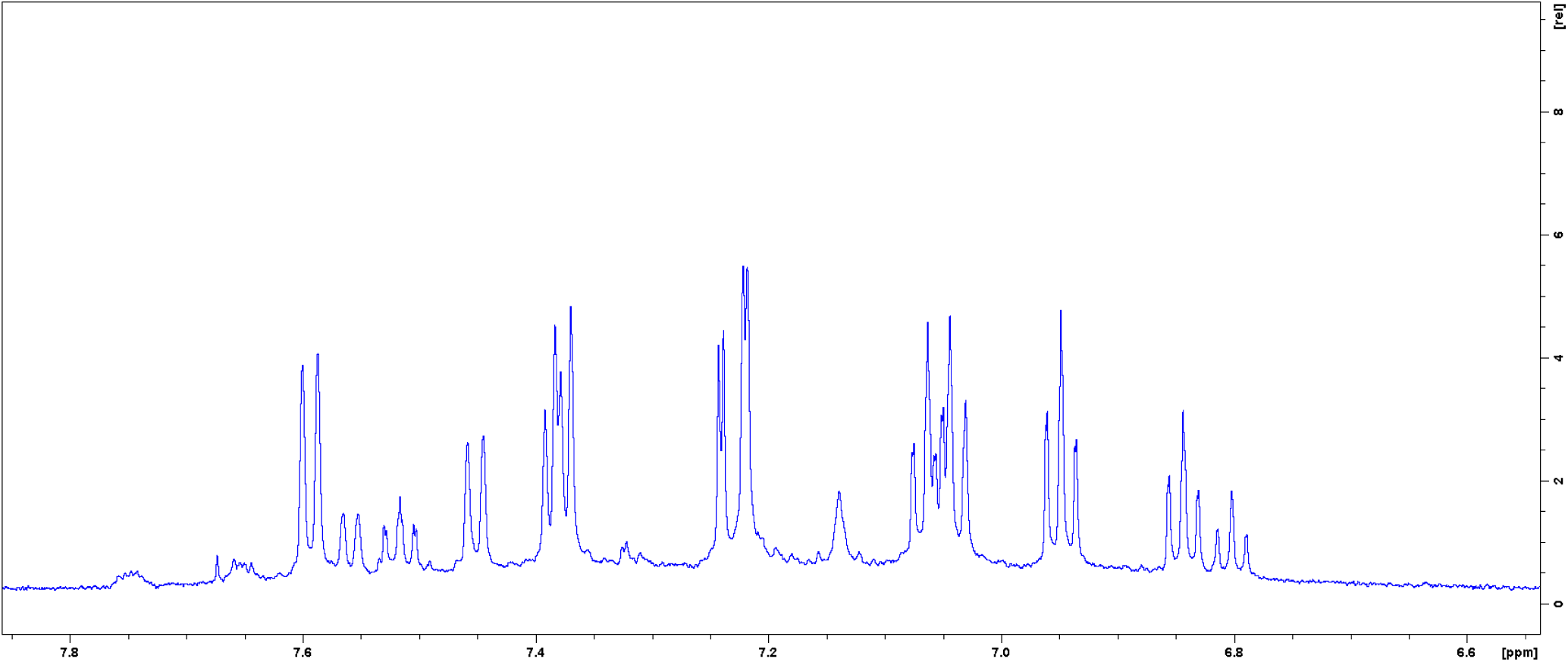
^1^H NMR spectrum (acetone-*d*_6_) of *Fn*-derived mixture of **1** and **2**, 6.6-7.8 ppm.

**S10.**
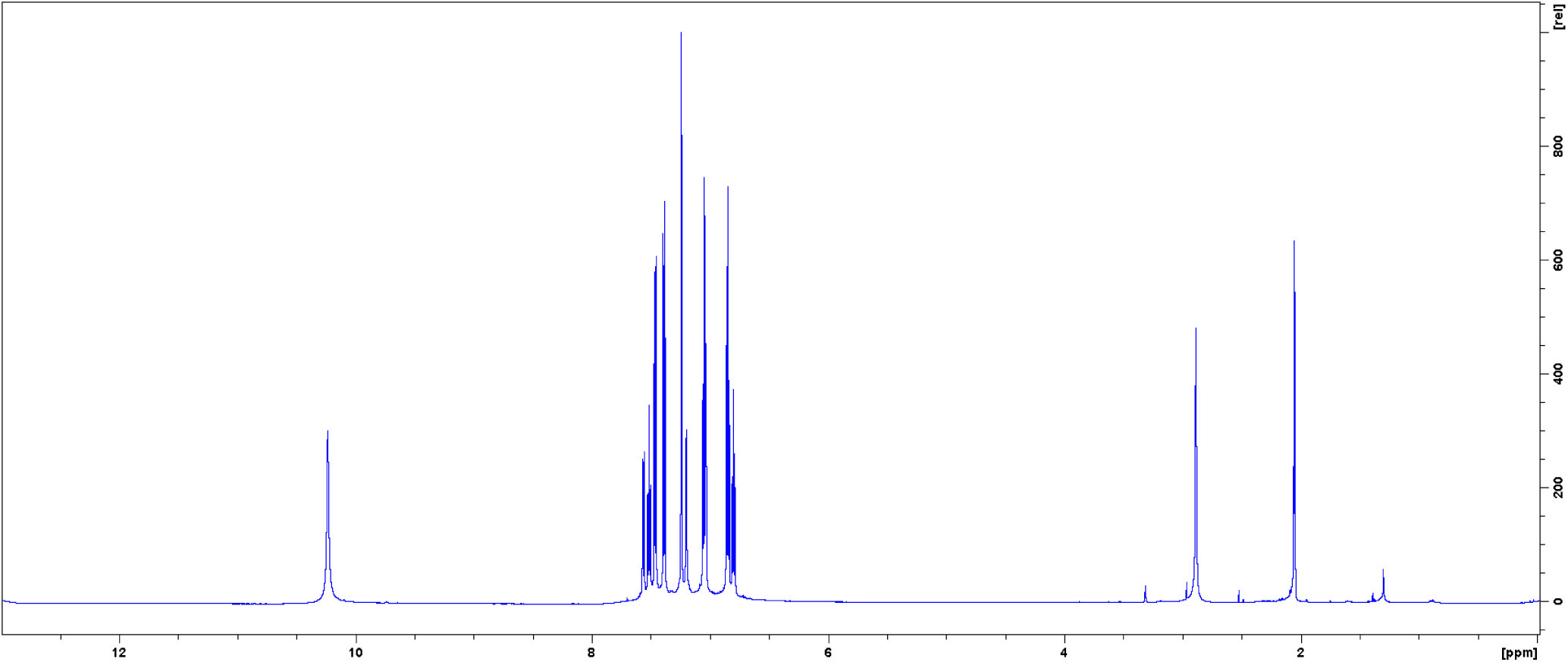
^1^H NMR spectrum (acetone-*d*_6_) of in-house synthesized FTIN (**1**).

**S11.**
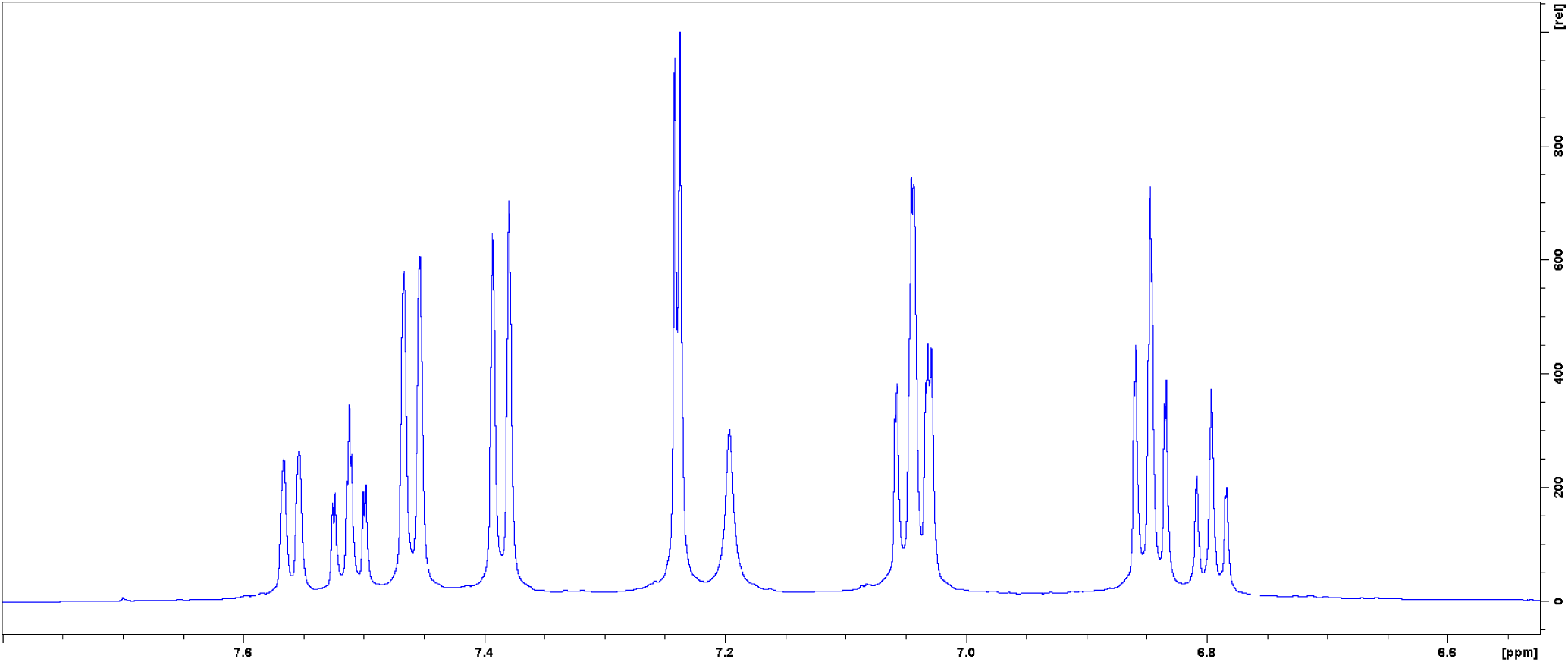
^1^H NMR spectrum (acetone-*d*_6_) of in-house synthesized FTIN (**1**), 6.6-7.8 ppm. S12.

**S12.**
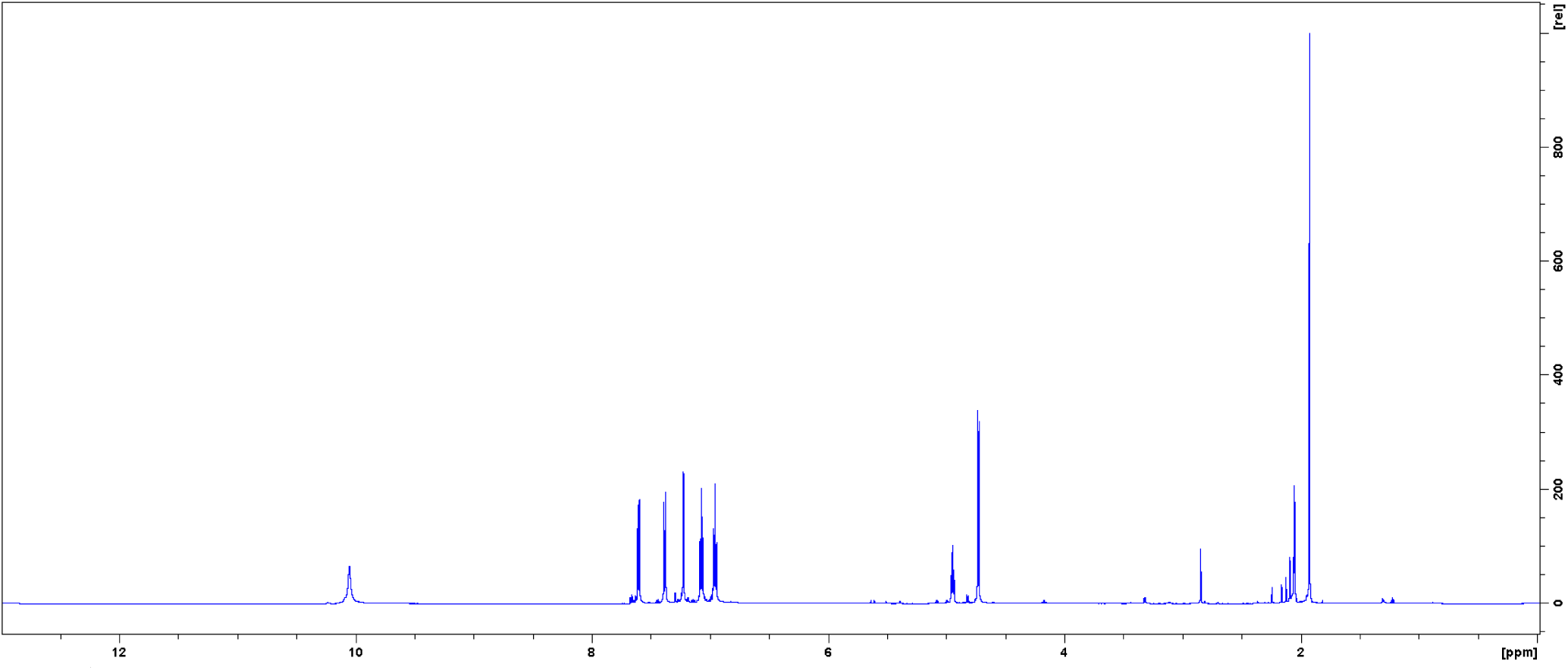
^1^H NMR spectrum (acetone-*d*_6_) of custom-synthesized STIN (**2**).

**S13.**
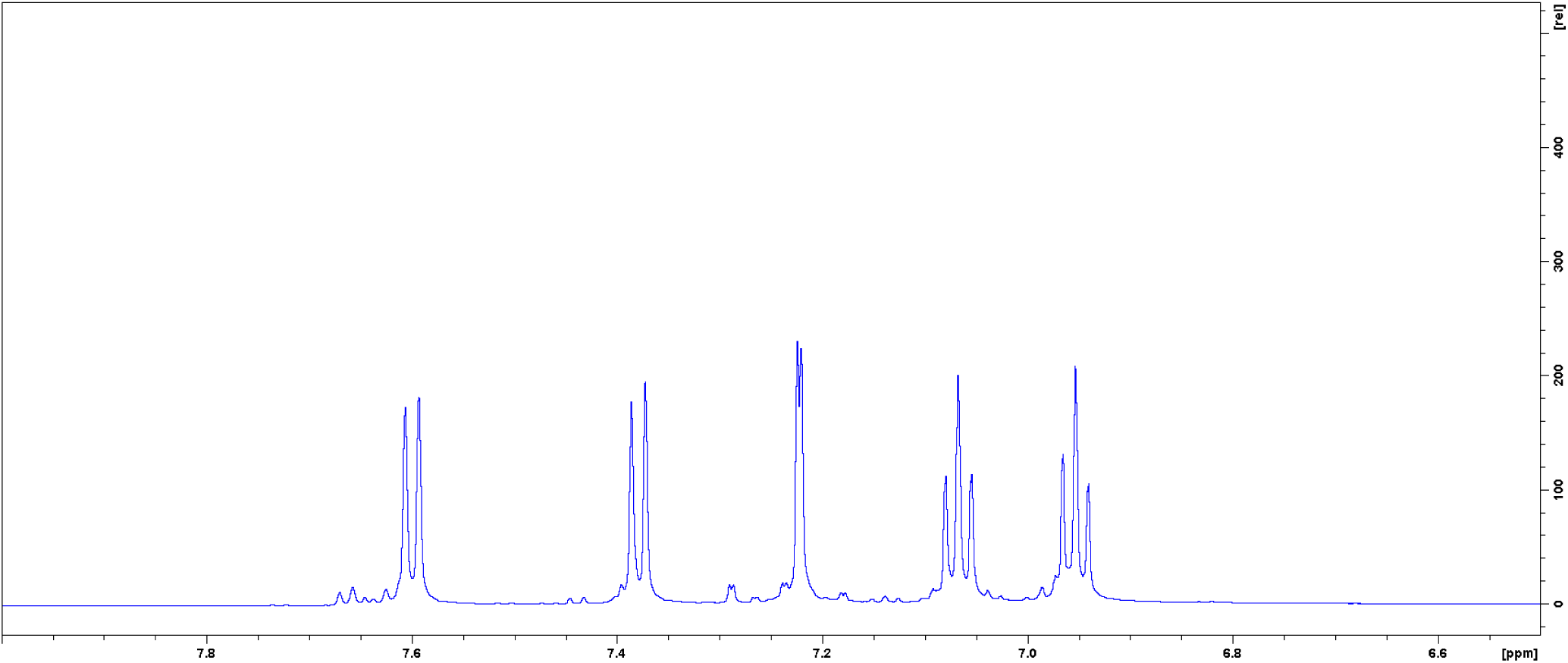
^1^H NMR spectrum (acetone-*d*_6_) of custom-synthesized STIN (**2**), 6.6-7.8 ppm.

**S14.**
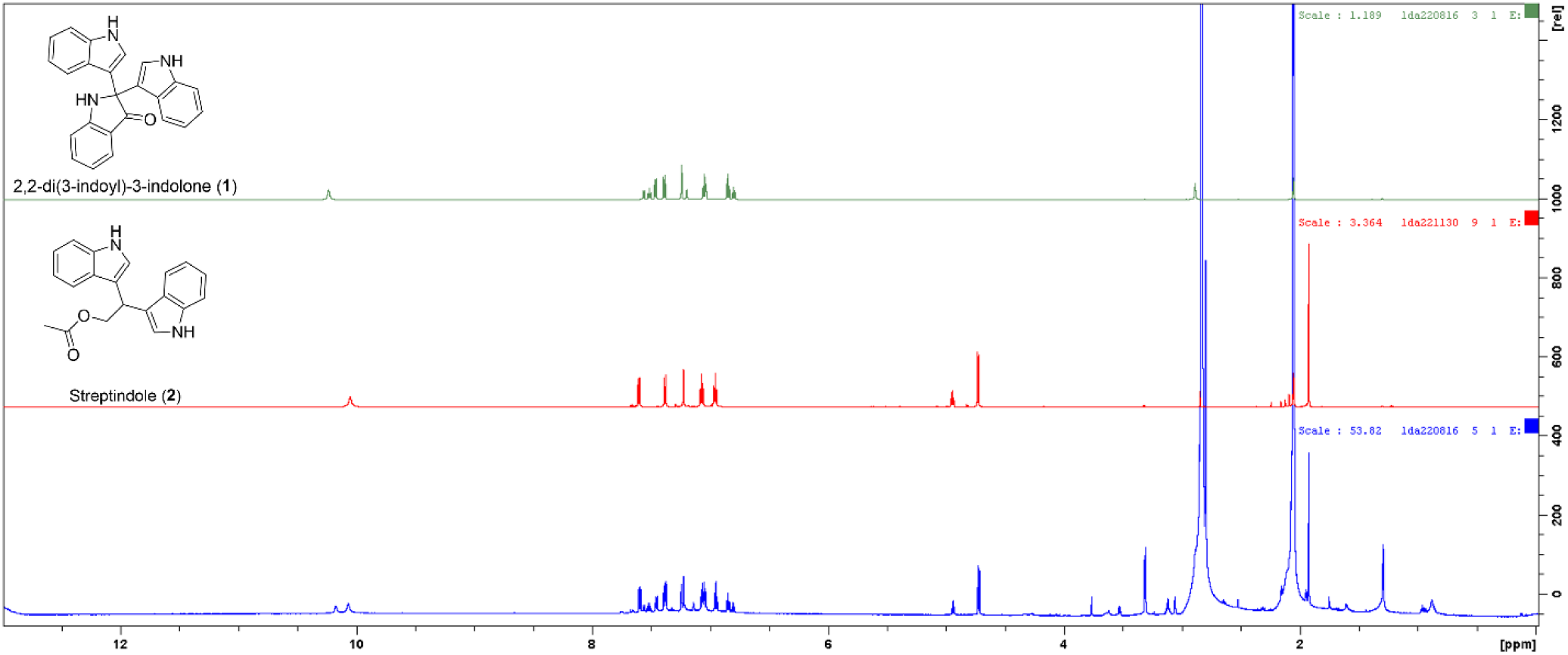
Stacked ^1^H NMR spectra (acetone-*d*_6_) of *Fn*-derived mixture of **1** and **2** (bottom, blue) compared to synthesized FTIN (top, red) and STIN (middle, green).

**S15.**
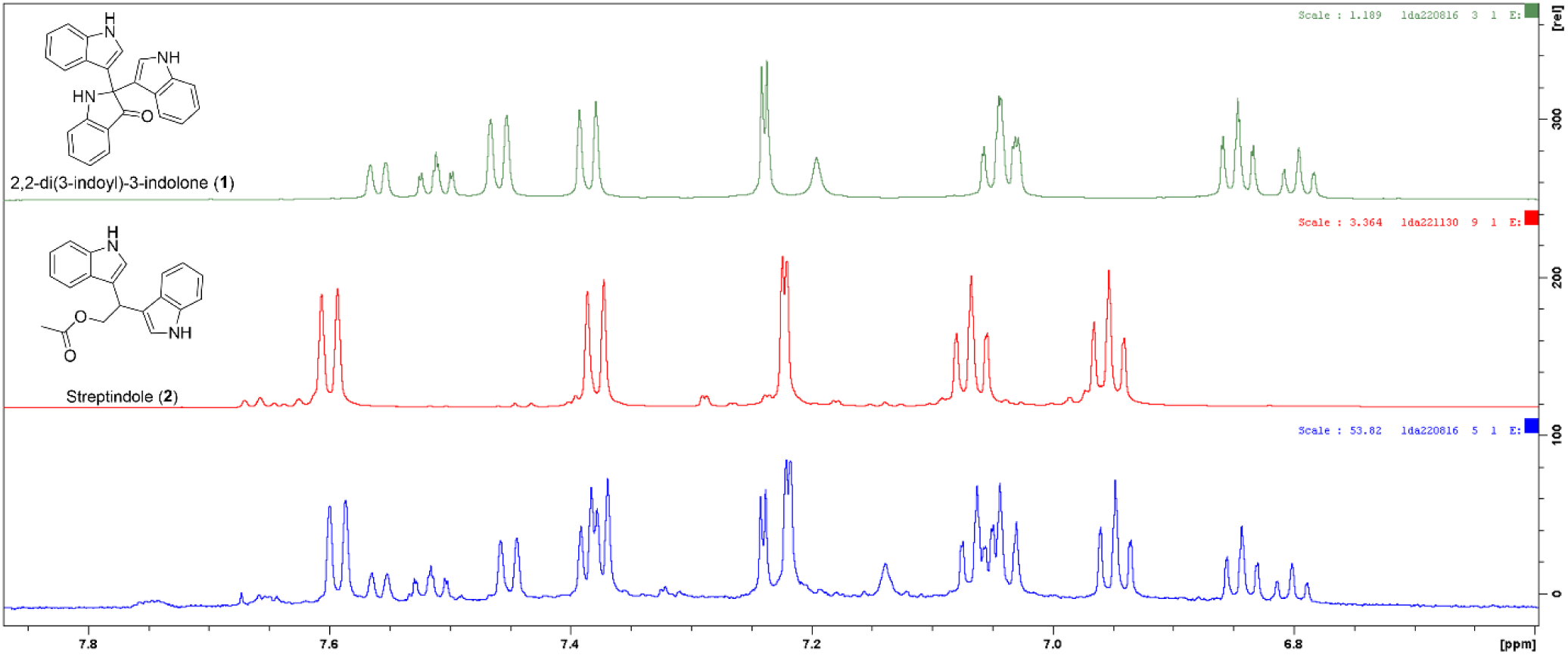
Stacked ^1^H NMR spectra (acetone-*d*_6_) of *Fn*-derived mixture of **1** and **2** (bottom, blue) compared to synthesized FTIN (top, red) and STIN (middle, green), 6.6-7.8 ppm.

**S16.**
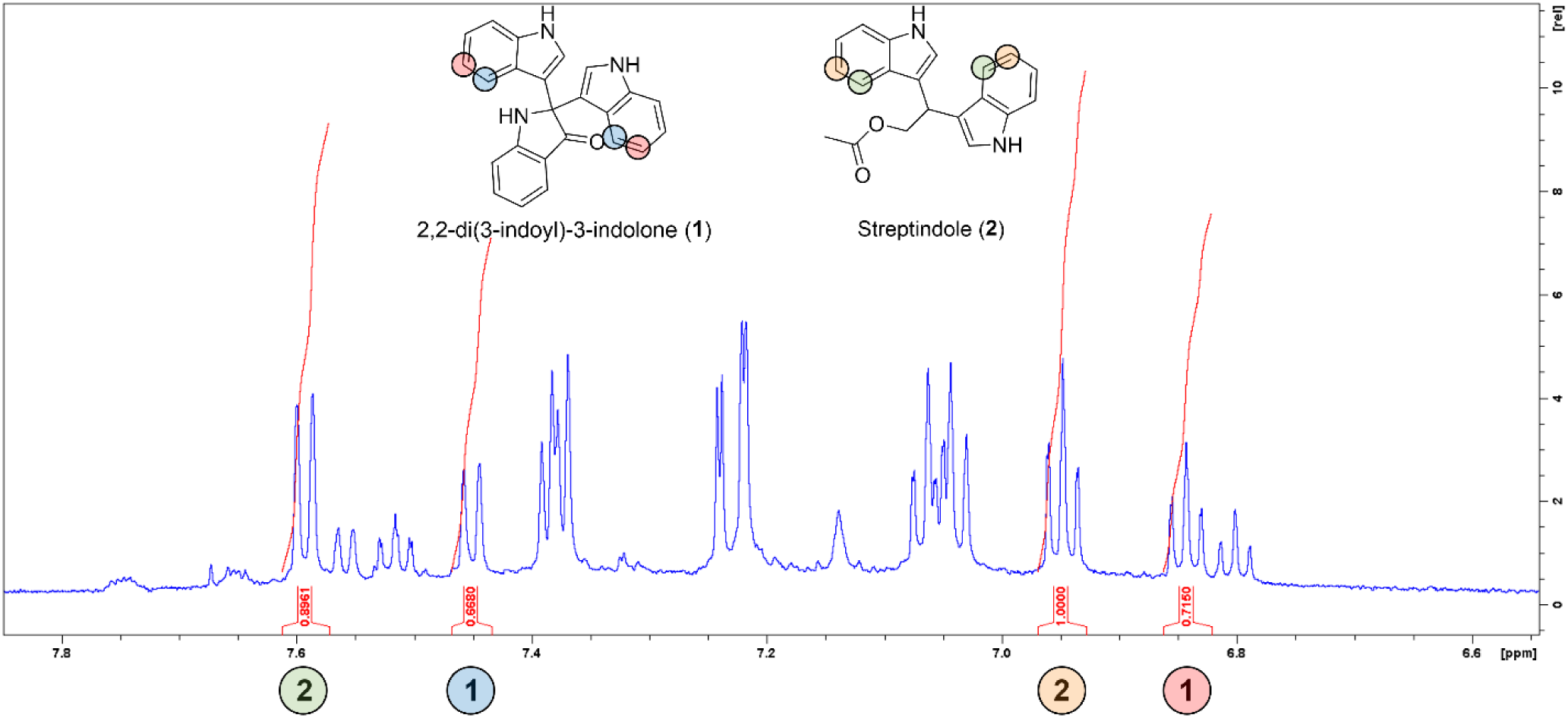
Annotated ^1^H NMR spectrum (acetone-*d*_6_) of *Fn*-derived mixture of **1** and **2** with signal integration labeled, indicating a 1:1 ratio.

